# A phylogenomic analysis of *Nepenthes* (Nepenthaceae)

**DOI:** 10.1101/680488

**Authors:** Bruce Murphy, Félix Forest, Timothy Barraclough, James Rosindell, Sidonie Bellot, Robyn Cowan, Michal Golos, Matthew Jebb, Martin Cheek

## Abstract

Nepenthaceae is one of the largest carnivorous plant families and features ecological and morphological adaptations indicating an impressive adaptive radiation. However, investigation of evolutionary and taxonomic questions is hindered by poor phylogenetic understanding, with previous molecular studies based on limited loci and taxa. We use high-throughput sequencing with a target-capture methodology based on a 353-loci, probe set to recover sequences for 197 samples, representing 151 described or putative *Nepenthes* species. Phylogenetic analyses were performed using supermatrix and maximum quartet species tree approaches. Our analyses confirm five Western outlier taxa, followed by *N. danseri*, as successively sister to the remainder of the group. We also find mostly consistent recovery of two major Southeast Asian clades. The first contains common or widespread lowland species plus a Wallacean–New Guinean clade. Within the second clade, sects. *Insignes* and *Tentaculatae* are well supported, while geographically defined clades representing Sumatra, Indochina, Peninsular Malaysia, Palawan, Mindanao and Borneo are also consistently recovered. However, we find considerable conflicting signal at the site and locus level, and often unstable backbone relationships. A handful of Bornean taxa are inconsistently placed and require further investigation. We make further suggestions for a modified infra-generic classification of genus *Nepenthes*.

## 1. Introduction

*Nepenthes* L. is the sole genus of Nepenthaceae, one of the largest families of carnivorous plants and part of the Caryophyllales. Nearly 300 species names have been published in the genus and there are an estimated 160–180 extant species (Clarke et al., 2018b; appendix 1). Carnivorous plants have always attracted considerable attention from scientists and the public alike, with *Nepenthes* in particular having some of the most dramatic and intriguing adaptations (Chase et al., 2009; Darwin, 1875; Ellison and Adamec, 2018; Fukushima et al., 2017; Juniper and Joel, 1989; Thorogood et al., 2018). Unlike their closest carnivorous relatives in Droseraceae, *Nepenthes* catch their prey within a fluid-filled pitcher that attaches to the leaf blade by a tendril. Although clearly circumscribed as a group by the universal presence of this unique pitcher organ, as well as a fundamentally homogeneous floral morphology, remarkable variation in pitcher form has been observed across the genus, from the giant, occasionally rodent-trapping pitchers of *N. attenboroughi*, to the delicate, rimless cups of *N. inermis* (Danser, 1928; Robinson et al., 2008). Furthermore, in recent years a range of studies have demonstrated various ecological adaptations related to nutrient acquisition and sometimes strongly linked to this morphological variation, such as the pitchers of *Nepenthes rajah*, adapted to catch the faeces of tree-shrews (Bauer et al., 2008; Ulrike Bauer et al., 2012b; Bazile et al., 2015; Clarke et al., 2009; Gaume et al., 2017; Lim et al., 2014; Merbach et al., 2002; Moran et al., 2012; Pavlovič et al., 2011; Scharmann et al., 2013; Scholz et al., 2010). These, along with variations of substrate and altitude, mark them the genus as a good putative example of an adaptive radiation (Bauer et al., 2012a; Clarke and Moran, 2016; Gaume et al., 2016; Thorogood et al., 2018). Many interesting questions remain about the evolution of key adaptations and their correlation with rates of diversification, the role of ecological divergence in speciation, and the importance of introgression and gene flow in the origin of new species or the maintenance of a wider pool of genetic diversity for variation (Thorogood et al., 2018).

Although the importance of ecological variation has been increasingly recognised for *Nepenthes*, geography is also likely to have strongly influenced speciation and diversification. Nepenthaceae is endemic to the palaeo-tropics, where its range extends from Madagascar to New Caledonia and from the Cape York Peninsula of northeastern Australia to southern China and the Patkai Hills in northeastern India (Jebb and Cheek, 1997; McPherson, 2009). However, the vast majority of species in the genus are endemic to one of three main regions within Malesia, i.e. Borneo, Sumatra and the Philippines, which are respectively home to around 44, 38 and 51 extant species, of which 37, 33 and 50 are endemics. The biogeography of the family have long been a focus for biologists, especially because of the centrality of the region to the history of this discipline (Wallace, 1869, 1860). Danser (1928) was a pioneering advocate of Wegener’s (1922) theory of continental drift and suggested that the handful of outlying western *Nepenthes* species in Madagascar, the Seychelles, Sri Lanka and India revealed an origin of the group on the ancient landmass of Gondwanaland. This theory has gained some support from molecular studies which have tended to place these species in varying order as the earliest-diverging lineages in the genus and successively sister to the rest of the genus (Alamsyah and Ito, 2013; Meimberg et al., 2001; Meimberg and Heubl, 2006; Mullins, 2000). However, the homogeneity of basic form and the evidence for recent, rapid diversification in at least some of the group put this ancient origin in doubt (Clarke et al., 2018a; Merckx et al., 2015; Schwallier et al., 2016). However, further investigation of these questions and of the effect of ongoing climate change on the future of the genus (Cannon et al., 2009; Schwallier et al., 2016) require a well resolved and accurate phylogeny.

The taxonomy and systematics of the group have also suffered from confusion and disagreement partly as a result of phylogenetic uncertainty (Clarke et al., 2018a). Relationships remain unclear at all taxonomic levels, from species delimitations and infra-generic relationships to the placement of Nepenthaceae in relation to other families. At the family level, Nepenthaceae has been repeatedly resolved in the Caryophyllales, within a partly carnivorous clade also containing Ancistrocladaceae, Dioncophyllaceae, Droseraceae and Drosophyllaceae (Heubl et al., 2006; Walker et al., 2018; Yao et al., 2019). However, relationships between these five families have been variously resolved, with only the sister relationship of Ancistrocladaceae and Dioncophyllaceae consistently recovered (Brockington et al., 2009; Heubl et al., 2006; Meimberg et al., 2000; Renner and Specht, 2011; Soltis et al., 2011; Walker et al., 2018; Yang et al., 2018, 2015; Yao et al., 2019). The carnivorous clade has usually been resolved as sister to the Frankeniaceae, Tamaricaceae, Plumbaginaceae and Polygonaceae (FTPP) clade, with this combined clade as sister to the rest of the Caryophyllales (Brockington et al., 2009; Cuenod et al., 2002; Smith et al., 2014, 2017; Soltis et al., 2011; Yang et al., 2018, 2015; Yao et al., 2019; but see Walker et al., 2018).

Uncertainty regarding evolutionary relationships has also limited the wide-scale adoption of subgeneric categories within *Nepenthes* itself. Danser (1928) suggested six infra-generic sections (sects. *Insignes, Montanae, Nobiles, Regiae, Urceolatae, Vulgatae)* but acknowledged that they did not all reflect evolutionary relationships. More recently, there have been two main conflicting approaches to adapting this classification in the light of molecular studies and taxonomic work. Cheek and Jebb (2015; 2017; 2016a, 2016b, 2015) have published an additional five sections *(Alatae, Micramphorae, Pyrophytae, Tentaculatae* and *Villosae)*, plus subsection *Poculae-ovis*, and the informal *Danseri* group, while rearranging many taxa. Clarke et al. (2018b) recognise only three of these new sections and also drop Danser’s *Nobiles* which was lectotypified by Cheek and Jebb (2015, 1999). These differences are summarised in Table 1, with full details of taxa in appendix 1. For both Cheek and Jebb (2015, 2009) and Clarke et al. (2018b) Danser’s *Montanae* and *Regiae* remain large groups, although their interpretations of these groups differ. In particular, Clarke et al. (2018b) restricted sect. *Montanae* to Sumatra and Java, placing all taxa from Peninsular Malaysia and Indochina (plus *N. albomarginata, N. hemsleyana, N. rafflesiana* and *N. reinwardtiana* which occur also in Borneo) in an extended sect. *Pyrophytae* The latter group was erected by Cheek and Jebb (2016b) for Indochinese species adapted to seasonally dry, fire-prone habitats. Cheek and Jebb’s (2013a, 2014, 2015) large sect. *Alatae*, containing many Philippine (mainly Mindanaoan) taxa, is not recognised by Clarke et al. (2018b), who placed most of these taxa in sections *Regiae* or *Villosae*. Cheek and Jebb’s small sect. *Micramphorae* (Cheek and Jebb, 2013b) is also not recognised but sects. *Insignes* and *Tentaculatae* are similarly defined by both groups of authors. Both of these small sections are notable for extending across Wallace’s line. Clarke et al. (2016b) rename Danser’s (1928) *Vulgatae* as sect. *Nepenthes* on nomenclatural grounds and move *N. gracilis, N. mirabilis* and *N. papuana* to the small *Urceolatae* group of other common, widespread lowland species. A small number of species, including limestone specialists *N. campanulata, N. northiana* and *N. mapuluensis*, defy confident taxonomic placement in any of the above groups (Table 1).

**Table 1.**
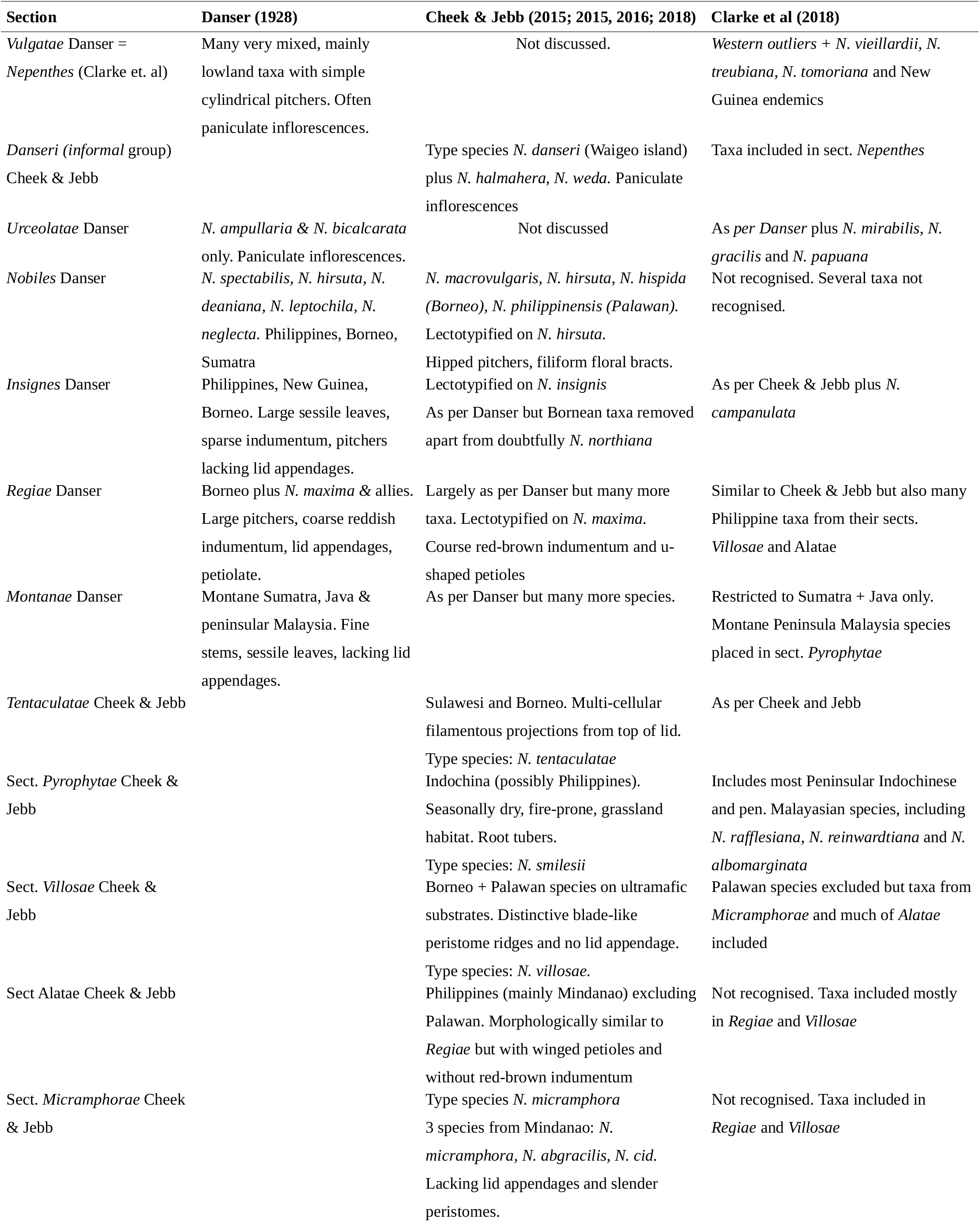
Summary of infrageneric classification of Nepenthes by three authorities.

At a lower level, species delimitation is also controversial. Clarke et al. (2018b) for example recognise only 127 species, with many recently described species particularly from the Philippines and Indochina considered synonyms of other taxa. In cases where one or more species have been segregated from an existing species there is potentially a question about the coherence of the remaining species, as for example with the recent separation of *N. orbiculata, N. parvula, N. rowanae* and *N. tenax* from *N. mirabilis* (Catalano, 2018; Clarke and Kruger, 2005, 2006; Wilson and Venter, 2016). There have also been few instances of multiple individuals being sequenced in previous molecular studies (but see Kurata et al. (2008). Therefore, it is unclear whether large geographical populations of widespread species form clades, or to what extent such populations are connected by gene flow. These uncertainties are not unique to *Nepenthes* but various factors make them important in this group: the frequency of natural hybrids and apparent lack of intrinsic reproductive barriers between taxa, the extent of intraspecific morphological variation and the reliance by taxonomists on the pitchers.

Three research groups using Sanger sequencing methods have produced the major molecular analyses of *Nepenthes* to date: Mullins’s (2000) nuclear and plastid study; the (2001) plastid study of Meimberg et al. (later supplemented with more species and a small nuclear dataset by Meimberg and Heubl, 2006); and Alamsyah and Ito’s (2013) study of the nuclear ITS marker. Although Mullins’s (2000) tree was well-sampled at the time, all these phylogenetic analyses lack a great many of the currently known species, are limited to just one or two markers and often show low branch support. Further efforts have sought to supplement these studies with more taxi or loci (Bunawan et al., 2017; Golos, 2012; Merckx et al., 2015; Renner and Specht, 2011; Schwallier et al., 2016) but major inconsistencies remain. Conflict between markers has been particularly noted by Mullins (2000). These studies have informed the infrageneric classifications described above but have also left room for different interpretations. High-throughput sequencing offers an opportunity to achieve greater resolution through sequencing more loci and thus resolve remaining inconsistencies. It also enables investigation of any conflicting signal between loci that may reveal a history of introgression, as suggested by Mullins (2000), or Incomplete Lineage Sorting (ILS). DNA target capture and sequencing is an increasingly popular next generation sequencing approach that makes it possible to sequence many loci in many species at a cheaper cost than whole genome sequencing and at a higher efficiency than genome skimming (Weitemeir et al., 2014). This approach has been used with success at the inter- and infra-specific level (Villaverde et al., 2018) but has never been used in *Nepenthes* where there is no available probe kit specific to the genus. However, a set of probes to capture 353 low-copy nuclear genes has been recently designed to work across angiosperms (Johnson et al., 2018). The variable flanking regions potentially also recovered with this approach may be of particular value at this taxonomic level given the likelihood of rapid recent diversification in at least some clades (Johnson et al., 2016; Schwallier et al., 2016).

In this study we test the performance of the angiosperm probe set to resolve infra-generic relationships in Nepenthaceae, taking advantage of the many *Nepenthes* species now available for sequencing following a rapid burst of species descriptions in the last two decades (Clarke et al., 2018a). We construct phylogenetic trees of 151 species and show that the 353 nuclear loci and their flanking regions are suitable to resolve most species into major clades. We interpret our findings in the context of previous molecular and taxonomic studies and suggest several taxonomic changes to make the infra-generic classification of the genus better reflect its evolutionary history.

## 2. Materials and Methods

### 2.1 Sampling

Leaf samples were collected in silica gel from the tropical nursery at the Royal Botanic Gardens, Kew and from the National Collection of *Nepenthes* at Chester Zoo. Additional specimens were sampled from the nursery of Andreas Wistuba (Germany), the private collection of Michal Golos (UK) and from wild specimens collected during field work in Indonesia. The remaining samples were taken from herbarium specimens housed at RBG Kew. Details of all sampled material are provided in Appendix 2. Several plants in cultivated collections were too small to take voucher herbarium specimens but living collection accession details are provided. Wherever possible, we also provide details of original collectors and localities., While the main aim of the sampling strategy adopted here was to represent as many putative species as possible, multiple samples were included for widespread species to represent different populations. Some intra-population samples were also included.

### 2.2 Extraction and library preparation

DNA was extracted from dried leaf tissue using a modified cetyl-tri-methylammonium bromide (CTAB) method, with chloroform:isoamyl alcohol (“Sevag”) separation and isopropanol precipitation at −20C (Doyle and Doyle, 1987). Additional cleaning was performed using Nucleospin silica columns (Machery-Nagel, Düren, Germany) or, for some herbarium specimens, Ampure XP paramagnetic beads (Beckman Coulter, High Wycombe, UK). Fragment size was estimated using an Agilent 4200 TapeStation (Agilent Technologies, Palo Alto, California, USA) for a few reference samples along with gel electrophoresis. Samples with fragment sizes >400 bp (all silica-dried samples) were then sonicated in Covaris AFA Fiber Pre-Slit Snap-Cap microTUBEs with a Covaris E220 Focused-ultrasonicator (Woburn, Massachusetts, USA) for 40 seconds with peak power set to 50W and duty factor at 20% to reduce most fragments closer to a target size of ca. 400 bp. Most herbarium samples were not sonicated.

Sequencing libraries were created from the fragmented genomic DNA using the NEBNext Ultra kit (New England Biolabs) with Ampure beads for size selection at 550 bp. Most libraries were prepared in half volumes to maximise reagents and indexes, with 200 ng (or minimum 50 ng) of fragmented DNA concentrated to a volume of 25 μL. Eight cycles of PCR amplification were used initially for all libraries, with further cycles, up to a total of 12, attempted for some herbarium samples. All library concentrations were quantified using a Quantus fluorometer (Promega Corporation, Madison, Wisconsin, USA) and their size measured using the TapeStation.

Up to 50 ng of library DNA was pooled in batches of up to 48 samples for hybridisation to the biotinylated probes using the Angiosperms 353 v 1 target capture kit (Johnson et al., 2018) available from Arbor Biosciences (Arbor Biosciences, Ann Arbor, Michigan, USA). Hybridisation was performed for 24 hours at 65 °C, followed by 12 cycles of PCR. Sequencing was performed at the Royal Botanic Gardens, Kew on an Illumina MiSeq (Illumina San-Diego, California. USA) with v 3 reagent chemistry (2 × 300 bp paired-end reads) or at Genewiz (Takeley, UK) on an Illumina HiSeq to produce 2 × 150 bp paired-end reads.

Technical replicate samples were included for one sample by using the same extraction of genomic DNA to create two separate libraries with different index combinations, which were then pooled and sequenced together. These were used to ascertain the potential sequence variability arising due to differential allele amplification or artefacts from the library preparation steps.

### 2.3 Sequencing and read trimming

Raw sequencing reads were cleaned with Trimmomatic (Bolger et al., 2014) in Illuminaclip palindrome mode to remove adapters using the paired end Illumina Tru-seq3 adapter file. The following settings were used based on experimentation: seedMismatches=1; palindomeClipThreshold=30; simpleClipThreshold=7; minAdapterLength=2; keepBothReads=TRUE. The keepBothReads option was used to retain post-trim forward and reverse reads. For general quality trimming, the MAXINFO option was used in preference to SLIDINGWINDOW as recommended in the application manual. The following settings were used with the MAXINFO option: targetLength=40, strictness=0.85. Finally, the MINLENGTH setting was also used with length set to 36 to remove shorter reads that might not be uniquely positionable against over sequences.

### 2.4 Recovery of target loci and flanking regions

The HybPiper pipeline v. 3 (Johnson et al., 2016) was used to recover coding sequences (CDS) for each locus from the trimmed reads, using a multi-species target file of amino acid gene sequences matching the probes from the 353 gene set of Johnson et al. (2018). HybPiper was run using the BLAST option (Altschul et al., 1990), in preference to BWA (Li and Durbin, 2009), as this was found to produce longer sequences. The optional pipeline module intronerate.py was used to recover “intron” sequences (concatenated fragments of introns and intergenic DNA) and “supercontig” sequences (CDS + concatenated intron fragments) from reads that extended beyond exon boundaries. After an initial run, the “supercontig” sequences from the sample *Nepenthes benstonei* were added to the target file and the pipeline run again. This samples was chosen because it had the highest number of reads of all ingroup samples produced from early sequencing batches and good sequence recovery (Table S1). We used the “--exclude” option to ensure that this additional target was only used for initial sorting of reads and not by exonerate for marking exon boundaries. The SPADES coverage cut-off option was set to four for this second run to maximise recovery. The pipeline was then used to retrieve CDS sequences for all samples into separate files for each of the 353 loci. An additional 351 files were created for intron loci, with two intron loci not recovered for any samples.

### 2.5 Alignment and approach to paralogs and missing data

Sequences < 25 bp were first removed from all intron files. Paralog warnings produced by the HybPiper pipeline for samples with multiple long contigs were further investigated by running the supplementary HybPiper scripts paraloginvestigator.py and paralogretriever.py to recover additional sequences for all putative locus copies (Johnson et al., 2016). We then generated alignments and trees with the additional sequences included as separate samples, and examined whether these grouped with each other or with different taxa on the resulting gene trees. Most warnings were produced for outgroup samples and for these the putative paralogous sequences did form clades, suggesting these were either not paralogs or were insufficiently diverged and unlikely to cause problems downstream. For the ingroup samples, gene trees were too inconsistent or poorly resolved to judge paralogs by this method. However, two loci, g6227 and g5919 were removed because they produced paralogs warnings for respectively 92 and 13 ingroup samples, while most warnings were for only 1 or 2 loci (see section 3.2), In addition, we aligned and compared the sequence identity of our two technical replicates (see sections 3.2, 3.11) for each locus. As a result we removed a further two loci which had outlying levels of pairwise variation between these samples, based on the rationale that they may represent paralogs.

Sequences were aligned using PASTA (Mirarab et al., 2015) which performs an iterative alignment process informed by tree generation steps between cycles of alignment. We used the MAFFT L-ins-i and other default settings within PASTA (Katoh and Standley, 2013). All alignments were subsequently “trimmed” using TrimaAl (Capella-Gutiérrez et al., 2009) to remove regions with gaps for many sequences or too much disparity between sequences. We used the “-automated1” option in TrimAl, which is optimised for downstream maximum likelihood (ML) tree reconstruction and chooses between a trimming method based purely on gap distribution and a method that also uses an automatically selected site similarity threshold. Outgroup sequences were removed from the intron files because they could not be aligned unambiguously. For the CDS sequences we created sequence files with and without the outgroups. All CDS alignments were concatenated to produce a supermatrix (hereafter called “AllCDS-supermatrix”) using AMAS.py (Borowiec, 2016). The same was done with ingroup sequences only and this version is referred to as “AllCDS-ingroup-supermatrix”.

Because missing data has been found to pose various problems for phylogenomic analyses (Hosner et al., 2016; Sayyari et al., 2017), we created a dataset with greater taxon representation comprising only the CDS ingroup loci that were recovered in at least 177 taxa. This cut-off was chosen because the number of taxa per loci dropped off steeply beyond this number, therefore allowing us to remove loci with a high amount of missing data while keeping many loci. The remaining 266 loci were concatenated to produce a supermatrix (hereafter called “177-supermatrix”) using AMAS.py (Borowiec, 2016). The same process was performed with the outgroup samples also included, using this time a threshold of 199 taxa per loci which resulted in 255 retained loci (hereafter “199-supermatrix”. These datasets were used in preference to the all-loci datasets to produce the supermatrix trees we discuss.

Datasets of individual locus alignments (Table 3) were also selected to be used for gene tree inference and maximum quartet support (MQS) species tree reconstruction with ASTRAL-III, which is statistically consistent under the multi-species coalescent model (Zhang et al., 2018). Taxon completeness of individual gene trees is arguably less important for this approach than maximising loci (Sayyari et al., 2017; Zhang et al., 2018). Therefore, for our main ingroup MQS dataset (hereafter “MQS-all-loci”) we included all CDS and intron loci, except for the putative paralogs mentioned above and five loci (three CDS, two intron) which were identified as high-end outliers for the proportion of parsimony informative characters, and therefore suspected of containing erroneous signal. This MQS-all-loci dataset comprised 338 CDS alignments and 339 intron alignments. For the analysis with outgroups (“MQS-CDS-alltaxa)”, we used CDS sequences only, resulting in 349 loci.

**Table 2.**
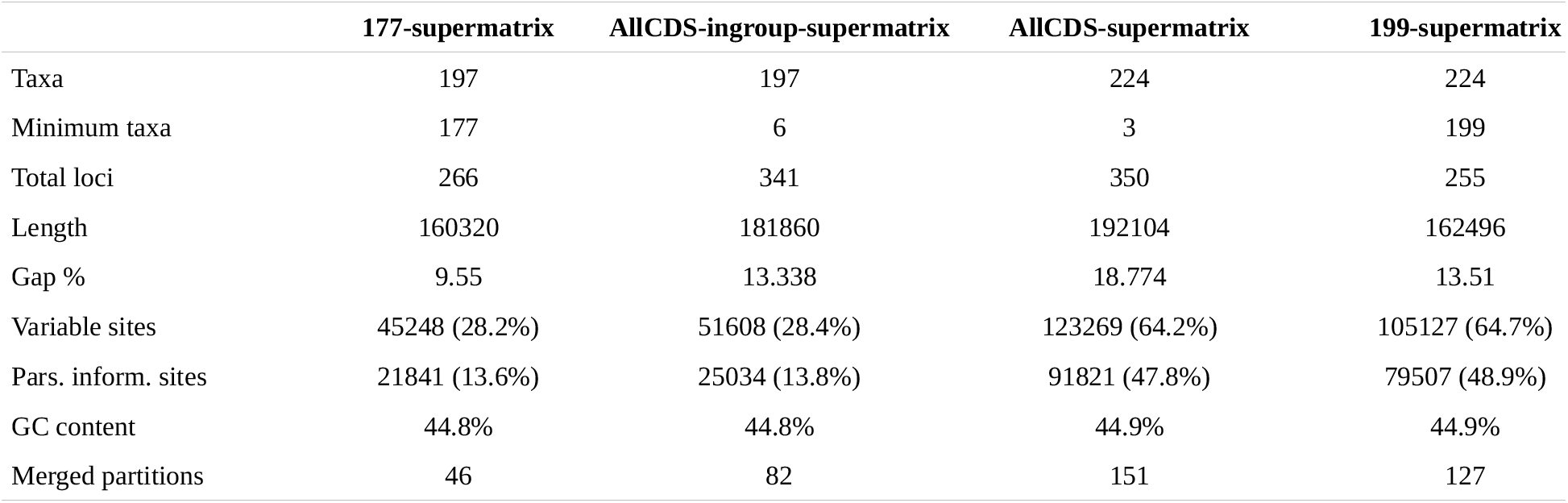
Summary statistics for concatenated matrices of trimmed CDS alignments. Merged partitions shows the revised number of partitions chosen by the partitionfinder software in Iqtree for ML tree reconstruction.

|  | **177-supermatrix** | **AllCDS-ingroup-supermatrix** | **AllCDS-supermatrix** | **199-supermatrix** |
| --- | --- | --- | --- | --- |
| Taxa | 197 | 197 | 224 | 224 |
| Minimum taxa | 177 | 6 | 3 | 199 |
| Total loci | 266 | 341 | 350 | 255 |
| Length | 160320 | 181860 | 192104 | 162496 |
| Gap % | 9.55 | 13.338 | 18.774 | 13.51 |
| Variable sites | 45248 (28.2%) | 51608 (28.4%) | 123269 (64.2%) | 105127 (64.7%) |
| Pars. inform. sites | 21841 (13.6%) | 25034 (13.8%) | 91821 (47.8%) | 79507 (48.9%) |
| GC content | 44.8% | 44.8% | 44.9% | 44.9% |
| Merged partitions | 46 | 82 | 151 | 127 |

**Table 3.** Median values (except where otherwise stated) for trimmed CDS and intron individual locus alignments. Median BS values are for ML gene-trees.

|  | **MQS-all-loci**<br>**(ingroup only)** | **MQS-CDS-loci**<br>**(ingroup only)** | **MQS-Intron-loci**<br>**(ingroup only)** | **MQS-super-loci**<br>**(ingroup only)** | **MQS-CDS-alltaxa**<br>**(all taxa)** |
| --- | --- | --- | --- | --- | --- |
| Total loci | 677 | 338 | 339 | 318 | 349 |
| Median no. taxa | 188 | 191 | 186 | 194 | 212 |
| Min no. taxa | 6 | 6 | 6 | 6 | 7 |
| Max no. taxa | 197 | 197 | 197 | 197 | 224 |
| Median length | 553 | 443 | 822 | 1843 | 443 |
| Max length | 6368 | 2740 | 6368 | 7251 | 2739 |
| Min length | 38 | 78 | 38 | 370 | 72 |
| Median gaps % | 6.26 | 3.8 | 9.212 | 12.52 | 6.71 |
| Max gaps % | 58.12 | 8.22 | 57.57 | 43.865 | 58.96 |
| Min gaps % | 0 | 0 | 0.171 | 1.132 | 0 |
| Median variable sites | 208 (44%) | 113.5 (25%) | 494 (63%) | 950 (52%) | 280 (64%) |
| Max % variable sites | 87.1% | 65.9% | 87.1% | 81.9% | 90.2% |
| Min % variable sites | 3.8% | 3.80% | 5.90% | 5.40% | 2.2% |
| Median pars. inform. sites | 102 (19.7%) | 54 (11.7%) | 239 (31%) | 496.5 (26.5%) | 203 (47.7%) |
| Max % pars. inform. sites | 61.8% | 49.8% | 61.8% | 55.5% | 76.6% |
| Min % pars. Inform. sites | 0.4% | 1.9% | 0.4% | 2.0% | 11% |
| GC content | 39.7% | 44.5% | 34.8% | 38.0% | 44.6% |
| Median BS | 57 | 48 | 64 | 72 | 54 |
| Median BS after collapsing<br>nodes <10 | 68 | 62 | 71 | 75 | 66 |

Loci with insufficient signal can be problematic for gene tree reconciliation methods such as ASTRAL because all input gene trees have equal weight and are assumed to be true (McCormack et al., 2013; Mirarab and Warnow, 2015; Zhang et al., 2018a). In an attempt to overcome the difficulties of short loci with limited signal, we also produced a set of longer alignments (hereafter “MQS-super-loci”) by concatenating individual trimmed CDS alignments with their corresponding trimmed intron alignments (Table 3). This achieves a similar purpose to “binning” loci (Bayzid et al., 2015) and is likely open to some of the same criticisms (Adams and Castoe, 2019), but although exons and introns of the same CDS can recombine, our approach may be more likely to respect the assumption of an absence of recombination inside the superlocus than binning loci located far apart in the genome. To ensure a relatively even representation of taxa across the length of the superlocus and therefore avoid large amounts of missing data in super-locus alignments, we only kept super-loci for which the difference in taxon number between the CDS and intron alignments was less than ten. This resulted in a total of 318 super-loci, ranging in taxon representation from 6 to 197 (median 194).

### 2.6 Gene-tree and species tree reconstruction

ML gene trees were generated for CDS, intron and super-loci alignments with IQTREE (Nguyen et al., 2015), using the Modelfinder program (Kalyaanamoorthy et al., 2017) to select an appropriate model for site frequency and rate heterogeneity. For the super-locus trees, a partitioned analysis (Chernomor et al., 2016) was used to accommodate different patterns of site and rate variation between the intron and CDS loci. The built-in ultrafast bootstrap (BS) algorithm (Hoang et al., 2018) was used to obtain 1000 BS replicates per locus. To evaluate gene tree resolution, we used custom scripts to calculate the mean BS value for each gene tree and for the whole dataset (Table S2). Gene tree nodes with a BS support value less than 10 were collapsed using the “nw_ed” function of Newick utilities (Junier and Zdobnov, 2010) to avoid arbitrary topologies detrimentally influencing the MQS tree. A more stringent cut-off value for collapsing nodes was not used because of evidence that this may sometimes be detrimental (Zhang et al., 2018b). MQS species trees were generated using ASTRAL-III (Zhang et al., 2018b) with the “-t3” option to calculate local posterior probabilities (LPP) (Sayyari and Mirarab, 2016).

To infer species trees from the 177-supermatrix and the AllCDS-supermatrix, we also used IQTREE v.1.6.9 with 1000 ultrafast bootstrap replicate (Hoang et al., 2018). We used the built-in partitionfinder2 option (Lanfear et al., 2016) to select the best partitioning scheme from an initial partition file based on individual loci, resulting in 46 and 127 partitions for the 177-supermatrix and the 199-ingroup-supermatrix respectively. Model selection was then automated for each partition using the built-in ModelFinder (Kalyaanamoorthy et al., 2017) with the free-rate model option included.

MQS and supermatrix trees were rooted using Phyutility (Smith and Dunn, 2008) to the clade consisting of all outgroups excluding the FTPP and carnivorous clades. Ingroup-only trees were re-rooted to *N. pervillei*. Trees were drawn using custom python scripts utilising the ETE toolkit (Huerta-Cepas et al., 2016). Gene and site concordance factors (GCF and SCF) were calculated using IQTREE based on the individual gene trees and alignments (Figs. 2, S1-2; Table S3-4; Ane et al., 2006; Gadagkar et al., 2005; Minh et al., 2018).

## 3. Results and Discussion

### 3.1 Sequencing results

Over the multiple sequencing runs, the number of trimmed reads for the in-group samples had mean 1,633,508, median 1,159,088, maximum 15,486,656 and minimum 390,316 (Table S1). Outgroup taxon reads were generated separately by the Plant and Fungal Tree of Life Project at RBG Kew; the number of trimmed reads had mean 1,273,127, median 1,027,119, maximum 4,972,230 and minimum 278,152. Reads mapped on target ranged from 0.84% to 11.6% for the ingroup (median 6.37%). Across all samples, the number of genes producing sequences at least 25%, 50% and 75% the length of the target had median 296, 227 and 130. Eight genes were never recovered for ingroup taxa but all of these produced sequences for at least some of the outgroup samples. Across all samples and genes, sequences averaged 59% of target length.

### 3.2 Paralog warnings

Du et al. (2019) have recently found that paralogs did not have a detrimental impact on species tree inference with ASTRAL. However, they could potentially introduce noise into supermatrix trees. Out of the total 67,584 CDS sequences produced for all samples and loci, the HybPiper pipeline produced paralog warnings for 289 sequences in total, of which 149 were from ingroup taxa. HybPiper gives paralog warnings when multiple long contigs are produced by SPADES (Johnson et al., 2016). We considered these in relation to taxa and in relation to genes. In the first case, taxa with warnings for many genes may have undergone whole genome duplication (WGD). In the second case, warnings from many different samples for a particular gene may indicate a gene duplication (paralogous gene). There were 135 taxa (114 ingroup taxa) with warnings for one or more sequences. Despite previous indications of WGD in *Nepenthes* (Walker et al., 2017; Knapek 2012) we found no evidence of recent WGD in the ingroup, with a maximum of 3 warnings for any *Nepenthes* taxon. In the outgroups, several taxa had higher numbers of warnings, with a maximum of 35 warnings for *Frankenia laevis*. With regard to individual genes, eighty-five genes produced paralog warnings for one or more sequences but only 26 genes produced warnings for any ingroup sequences. For most of these genes, warnings were given for only 1 or 2 ingroup sequences but two genes were excluded from subsequent analyses for having 13 and 92 putatively affected ingroup sequences, thus removing over two thirds of the 149 ingroup sequences with warnings.

To test if the warnings were actually caused by paralogs or had other explanations such as heterozygosity, we produced gene trees with both alternative sequences included for the 85 affected genes (Johnson et al., 2016). We found little sign of ancient gene duplication, with most trees showing sequences from the same sample grouped together, or a general lack of resolution.

However, gene g6227 with 92 affected in-group sequences was the exception. We then mapped the CDS sequences with paralog warnings for this gene on to the species tree (Fig S3), where they suggest a gene duplication somewhere on the branch leading from the Insignes clade to the rest of clade 2. To further check that recovered genes were homologous between samples, we also calculated pairwise variation between our replicate samples. We found alignments of these two samples were non-identical after gap-removal for 93 out of 209 CDS sequences recovered for both samples, although only 5 of these genes produced warnings in HybPiper. Again, heterozygosity or noise is likely responsible for most of this variation but two samples with >5% pairwise variation pairwise difference between replicates were removed from the analyses.

### 3.3 Alignment summary statistics

The CDS 177-super-matrix alignment of ingroup taxa retained 266 loci of the 341 total CDS loci and reduced the missing data from 13.4% to 9.6% with a reduction in length of 21540 bp and 3193 parsimony-informative sites (Table 2). The proportion of variable and parsimony-informative sites remained essentially the same, as did the GC content. With outgroups included, the proportion of variable and parsimony informative sites was much higher.

In the all-locus MQS tree dataset there was a median of 188 of the total 197 taxa included per locus, with a minimum representation of 6 taxa (Tables 3, S5). The median locus length was 553 bp, with 208 (43.9%) variable sites, 102 (19.7%) parsimony-informative sites and 6.26% gaps. The intron loci greatly increased the number of parsimony informative sites but at a cost of significantly more missing data. Our super-loci of combined intron and CDS sequences had a median of 496.5 parsimony informative sites, nearly five times greater than for the all-locus set. This resulted in much higher median bootstrap scores in the gene trees (Tables 3, S2).

### 3.4 Family-level relationships

In contrast to Walker et al. (2018), our MQS and supermatrix trees (Fig. 1) replicated the family-level relationships found by previous studies, resolving with strong bootstrap (BS) and LPP support the carnivorous clade as sister to the FTPP clade, and this combined clade as sister in turn to the rest of the Caryophyllales. Within the carnivorous clade, Droseraceae was recovered as sister to the other families and Nepenthaceae as sister to the clade containing Drosophyllaceae, Ancistrocladaceae and Dioncophyllaceae. This relationship was also recovered by Walker et al. (2017, 2018) although Dioncophyllaceae was not sampled there. Our similarity to these studies may reflect the phylogenomic scope of both studies and the influence of the nuclear portion of the genome since a recent plastome study (Yao et al., 2019) as well as earlier studies using plastid markers and fewer nuclear genes found different relationships (Brockington et al., 2009; Soltis et al., 2011). The topology found here accords with the topology and interpretation of carnivorous adaptations suggested by Renner and Specht (2011, 2013). Fleischmann et al. (2018) argue that features such as echinate pollen tetrads and the presence of both plumbagin and its isomer 7-methyljuglone in Droseraceae and Nepenthaceae support a topology (based on APGIV data (2016)) with a clade of those two families sister to the other families. If the topology presented here is correct, either the three other families must have reverted back to monads, or tetrads evolved independently in Droseraceae and Nepenthaceae. However, if the chemical features represent a symplesiomorphy with Plumbaginaceae, as suggested by Fleischmann et al. (2018), then the topology of Fig. 1 is arguably no less consistent with these characters than theirs.

**Fig. 1.**
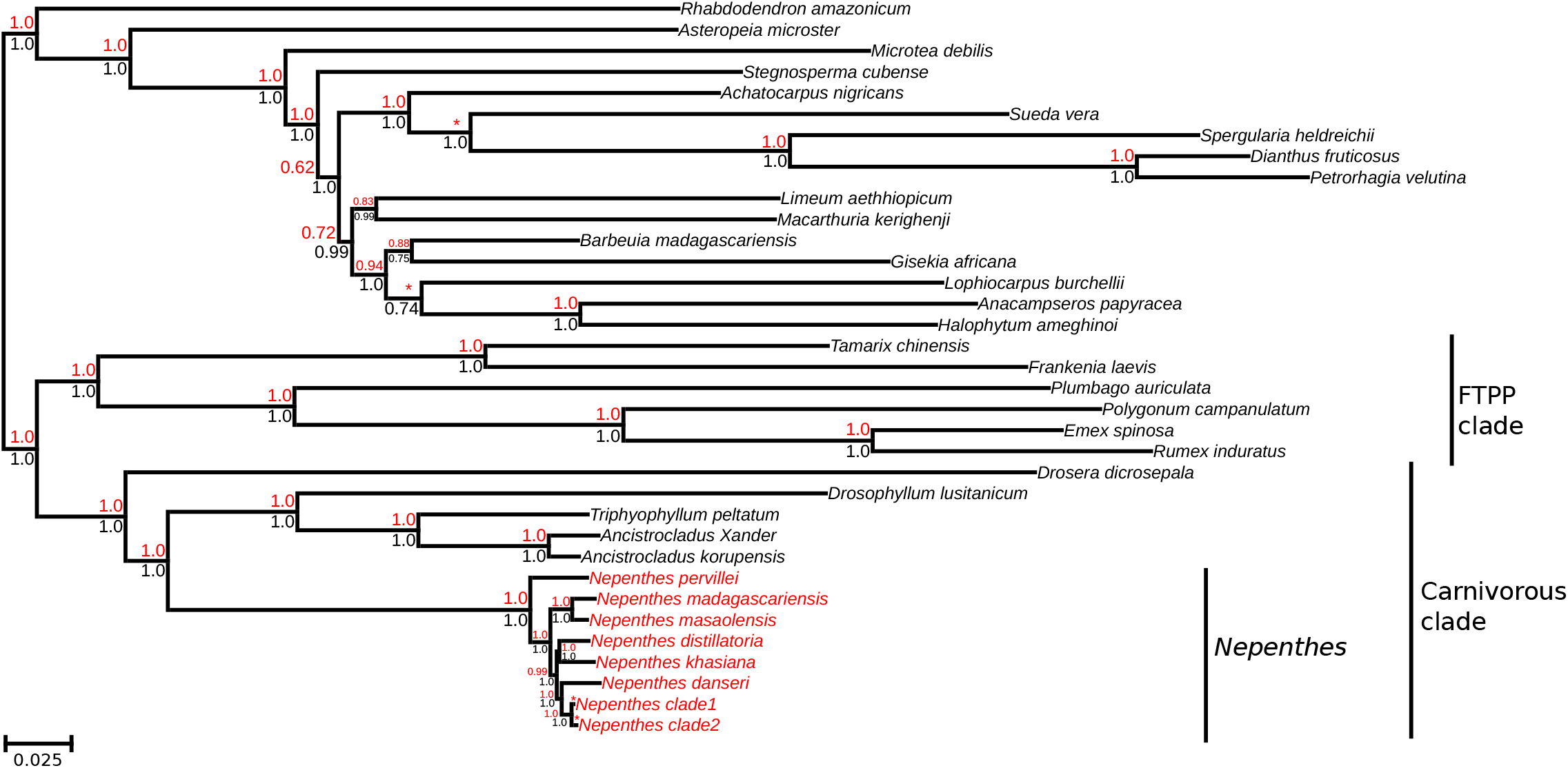
ML 199-Supermatrix tree of family level relationships (major ingroup clades collapsed). Generated with IQTREE with Partitionfinder and Modelfinder options from concatenated CDS loci exceeding 198 taxa. Support values displayed below branches are ultrafast BS values from 1000 replicates. Above branch values (red) are pp values (ASTRAL “-t3” option) from the MQS-CDS- alltaxa ASTRAL analysis. Clades not recovered in the MQS-CDS-alltaxa tree are marked with an asterix. Tree is rooted by core Caryophyllales clade (all out-groups excluding carnivorous clade, Plumbaginaceae and FTPP clade).

### 3.5 *Nepenthes*—major topological findings

Our trees (Figs. 1–4) are largely consistent in the recovery of major clades but there is variation in the positioning of some of these. We find a grade of five western species and *N. danseri* as sister to the rest of the genus which is divided into two main clades. These two clades are consistently recovered except for the position of *N. angustifolia* which is resolved in Clade 1 on the supermatrix tree and in Clade 2 on both MQS trees. Clade 1 contains most of the common, lowland widespread species, traditionally classified in sects *Vulgatae/Nepenthes* and *Urceolatae*, including *N. mirabilis* (and allies), *N. gracilis, N. ampullaria, N. rafflesiana* and *N. bicalcarata*, but also the *N. tomoriana* clade of species from Sulawesi and New Guinea which includes high altitude endemics. Clade 2 contains the wide-ranging sections *Tentaculatae* and *Insignes* as well as geographically defined clades from Sumatra, Indochina, Peninsula Malaysia, Mindanao, Palawan, and Borneo–Wallacea–New Guinea. These contain species mainly from sects. *Montanae, Pyrophytae, Alatae, Villosae* and *Regiae*, depending on the taxonomic scheme. Clade 2 also contains a number of taxa from Borneo which are unstably placed between trees and have lower support values. Our topologies appear to suggest at least seven dispersals across Wallace’s line (once in the *Regiae* clade, *Tentaculatae* clade, *Insignes* clade, stem leading to *N. danseri* and stem leading to the *N. tomoriana* clade, and twice within the *N. mirabilis clade)*.

### 3.6 Western outliers and *Nepenthes danseri*

In both MQS and supermatrix analyses, the Western outlying species form a paraphyletic grade outside the main clade of species (Figs. 1–4). *Nepenthes pervillei* from the Seychelles was recovered as sister to all other *Nepenthes* with maximal support in both MQS and 199-supermatrix analyses (Fig. 1). This species has always been found to be either sister to the rest of the genus or part of a polytomy outside the main clade of species (Alamsyah and Ito, 2013; Meimberg and Heubl, 2006; Mullins, 2000). It has also been recognised as morphologically unique. A distinct seed morphology without filaments led Hooker (1873) to place it in a separate subgenus *Anourosperma*. This trait could plausibly be interpreted as due to selection against wind dispersed seeds on a small oceanic island and such adaptations could evolve quickly and may not represent a plesiomorphy or a long, independent evolutionary history (McPherson, 2009). However, our trees do show quite a long branch between this species and the rest of the genus.

The sister pairs of *N. masaolensis* plus *N. madagascariensis*, and *N. khasiana* plus *N. distillatoria*, and the single species *N. danseri* were successively sister to all remaining species with maximal LPP and BS support in MQS and supermatrix trees (Figs. 1–4). While similar topology has been recovered before, the order of branching of these species has not been decisively established (Alamsyah and Ito, 2013; Meimberg and Heubl, 2006; Mullins, 2000). Our analyses confirm *N. danseri* from Waigeo island off New Guinea as sister to all the Southeast Asian species. Although we have yet to perform biogeographical analyses, the geographical book-ending of the main radiation by these Western and Eastern species is notable. GCF and SCF values are somewhat higher in this part of the tree than for the other internal nodes in the tree but there are still relatively few genes concordant with the species topology. For example, the sister relationship of *N. khasiana* and *N. distillatoria* (branch id 252) has 28% gene concordance but 9% and 15% for the two next most supported topologies, while the SCF score of 39% is only a little higher than equal support for the three alternative quartet topologies (Figs. 2, S1, Table S3).

**Fig. 2.**
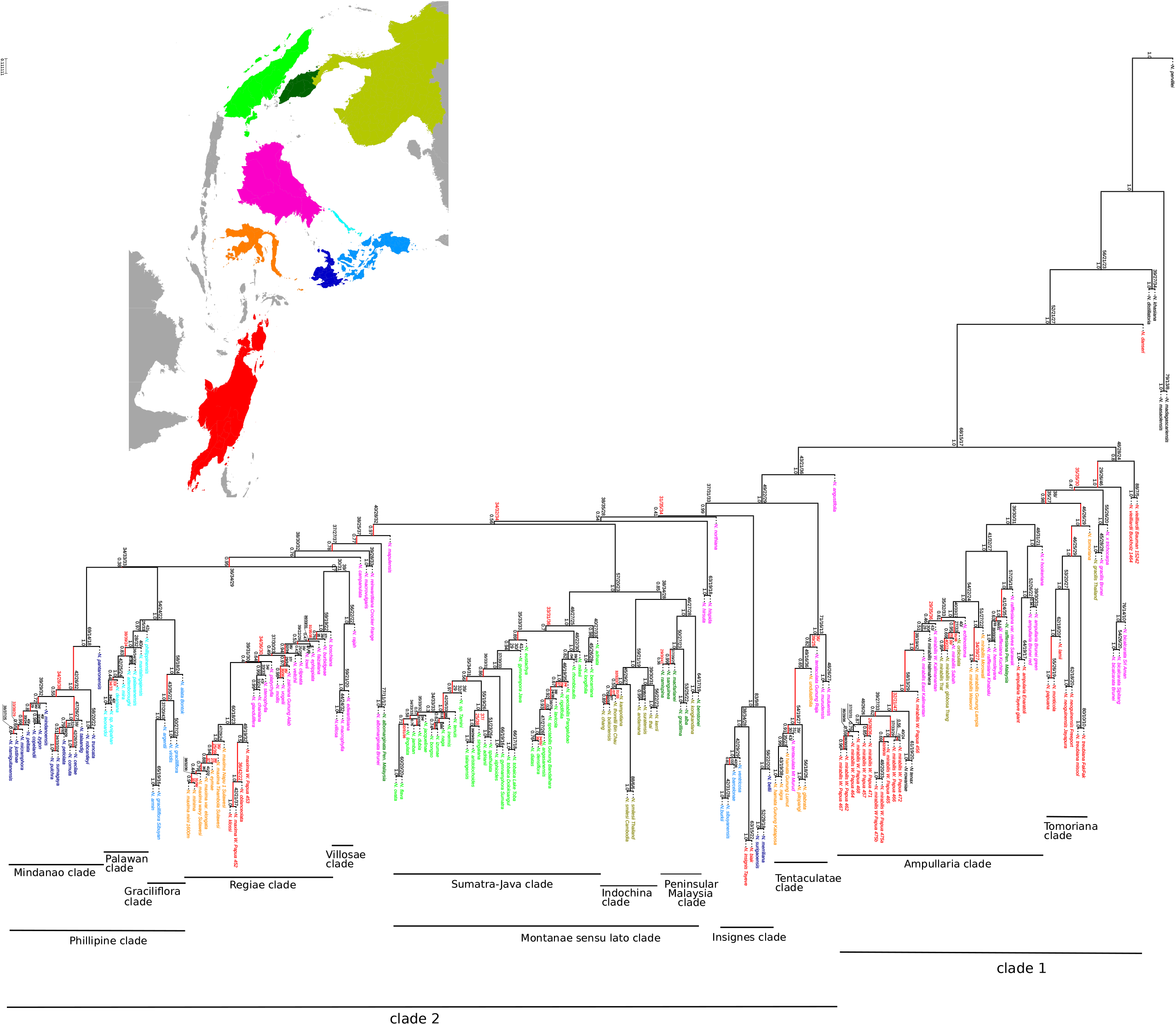
MQS-all-loci tree for ingroup taxa generated with ASTRAL III based on ML partitioned gene trees from all loci. Branch lengths for internal branches are in coalescent units (generations/ effective population size). Dotted line branches indicate arbitrary terminal branch lengths not calculated by ASTRAL. Tree is rooted to *N. pervillei* based on results in Fig. 1. Support values below displayed branches are local posterior probabilities (-t 3 option) described in Sayyari and Mirarab (2016). Above branch values are site concordance and discordance (SCF/SDF1/SDF2) scores generated with IQTREE based on quartet-trees for individual sites. The three above branch scores for a node add up to 100% (before rounding). Branches not recovered in the 177-supermatrix ML analysis (Fig 3.) are shown in red. Coloured tip names correspond to the accompanying map.

### 3.7 *Nepenthes* Clade 1

Our phylogenetic trees consistently recover Clade 1 comprising the same set of species except for *N. angustifolia* which is resolved as sister to Clade 2 in our MQS trees (Figs. 2, 4). This taxon has usually been considered a variant of *N. gracilis* (Danser 1928; Cheek and Jebb, 2001) and therefore it seems likely that the MQS topology is erroneous. However, ILS or introgression could be responsible or *N. angustifolia* may indeed be a separate but closely related species to *N. gracilis*. Our sample was taken from cultivated material of uncertain origin and therefore further sampling from wild populations would be necessary to clarify the relationships of this taxon.

In all trees, Clade 1 contains several common, lowland and mostly widespread taxa placed in sects *Vulgatae* or *Urceolatae (N. mirabilis, N. ampullaria, N. rafflesiana, N. gracilis* and *N. bicalcarata)* as well as an *N. tomoriana* clade of Wallacean and New Guinean endemics. It is notable that, like *N. danseri* and the Western outliers (except *N. khasiana)*, many of these species have paniculate inflorescences with 3 or more flowers per partial peduncle, rather than 1 or 2 as in most of the genus. *Nepenthes vieillardii* is inconsistently placed as sister to the rest of Clade 1 or within the *N. tomoriana* clade. The various widespread taxa were also resolved together in the nuclear tree of Mullins’s (2000), although conflicting plastid data was interpreted as indicating deep introgression of these taxa. While we cannot rule out plastid capture, we find no more evidence of discordant nuclear signal for these taxa than for the rest of the genus. While of interest in relation to dispersal and pollination patterns in a dioecious genus (Petit and Excoffier, 2009), the plastid is usually considered to represent only a single, non-recombining locus and therefore its significance compared to hundreds of nuclear loci is debatable. The two known natural hybrid taxa we sampled, *N. × hookeriana (N. rafflesiana × N. ampullaria)* and *N × trichocrapa (N. gracilis × N. ampullaria)*, are resolved inconsistently with either of their parent taxa on our trees. The multiple samples of *N. mirabilis* in this clade are discussed further below.

The *N. tomoriana* clade is well supported and, with the exception of *N. vieillardii* mostly consistent between our trees. The order of branching within this clade suggests an eastward radiation since *N. tomoriana* is found in the Tomori Bay area on the eastern coast of Sulawesi, while *N. treubiana* (sampled here from both the island of Misool and the western coast of Indonesian Papua) is sister to the other species which extend further inland and to higher altitudes. Wallace’s line was presumably crossed by ancestors on the stem to this clade. This *N. tomoriana–New* Guinea clade was not recovered by Mullins (2000) or Alamsyah and Ito (2013) but was resolved with moderate support on the plastid *trnK* phylogeny of Meimberg and Heubl (2006), with the same order of internal branching. This consistency with our nuclear genomic sampling and our inclusion of the more Western, Misool population of *N. treubiana* provides good evidence for this topology and for potential biogeographical interpretation. The recovery of the racemose species, *N. papuana*, in this clade agrees more with the plastid tree rather than the nuclear tree of Mullins (2000) and also conflicts with the taxonomic placement of this species in sect. *Urceolatae* by Clarke et al. (2018b). Further paniculate species from this region, *N. paniculata, N. weda* and *N. halmahera*, were not sampled here. While we would expect the former to belong in this clade based on biogeography and morphology, the latter two species appear more closely related to *N. danseri* (Cheek, 2015).

Our samples of *N. vieillardii* and *N. bicalcarata* and *N. gracilis* are recovered in that order as successive sister clades to the rest of clade 1 on our MQS analysis (Fig. 2, PP: 1, 0.47 and 1 respectively). The super-loci MQS species tree is similar but the position of the clades containing samples of *N. bicalcarata* and *N. gracilis* are reversed. However, on our supermatrix tree (Fig. 3), *N. vieillardii* is recovered with maximal support within the *N. tomoriana* clade as sister to the New Guinea endemics. Although it is hard to be confident in either topology, we suggest the supermatrix topology is more likely in this instance. *Nepenthes vieillardii* is endemic to New Caledonia and the sole *Nepenthes* species found there. Therefore, the supermatrix topology, which resolves it as sister to the endemic New Guinea taxa (Fig. 2), is biogeographically plausible, although not suggested by previous molecular studies (Alamsyah and Ito, 2013; Meimberg and Heubl, 2006; Mullins, 2000). Furthermore, *N. lamii* and *N. monticola* from the New Guinea clade were previously considered synonymous with this species by Danser (1928). Our two *N. vieillardii* samples were taken from herbarium specimens and sequence recovery was poor. While missing data is likely to have increased the general problem of limited signal in individual gene trees, the accumulated signal in the supermatrix alignment may be adequate to resolve the samples correctly. It is noticeable that several taxa with poor recovery have very long branches on the supermatrix tree including *N. vieillardii*, a problem known to occur with missing data (Darriba et al., 2016). This could result in long-branch attraction and misplacement of taxa in both the supermatrix tree and in individual gene trees and consequently the MQS species tree. We tried to limit the amount of type 2 missing data in our supermatrix alignment by removing loci with fewer than 177 (ca. 90%) of 198 samples represented but this did not completely eliminate the long-branch problem. However, other taxa with long branches apparently due to missing data (e.g. *N. pantorensis, N. rafflesiana var*. Nivea, *N. spectabilis* Pangulabao and *N. undulatifolia)* do appear to be placed in appropriate clades on the supermatrix tree, suggesting the placement of *N. vieillardii* in the supermatrix tree may be accurate.

**Fig. 3.**
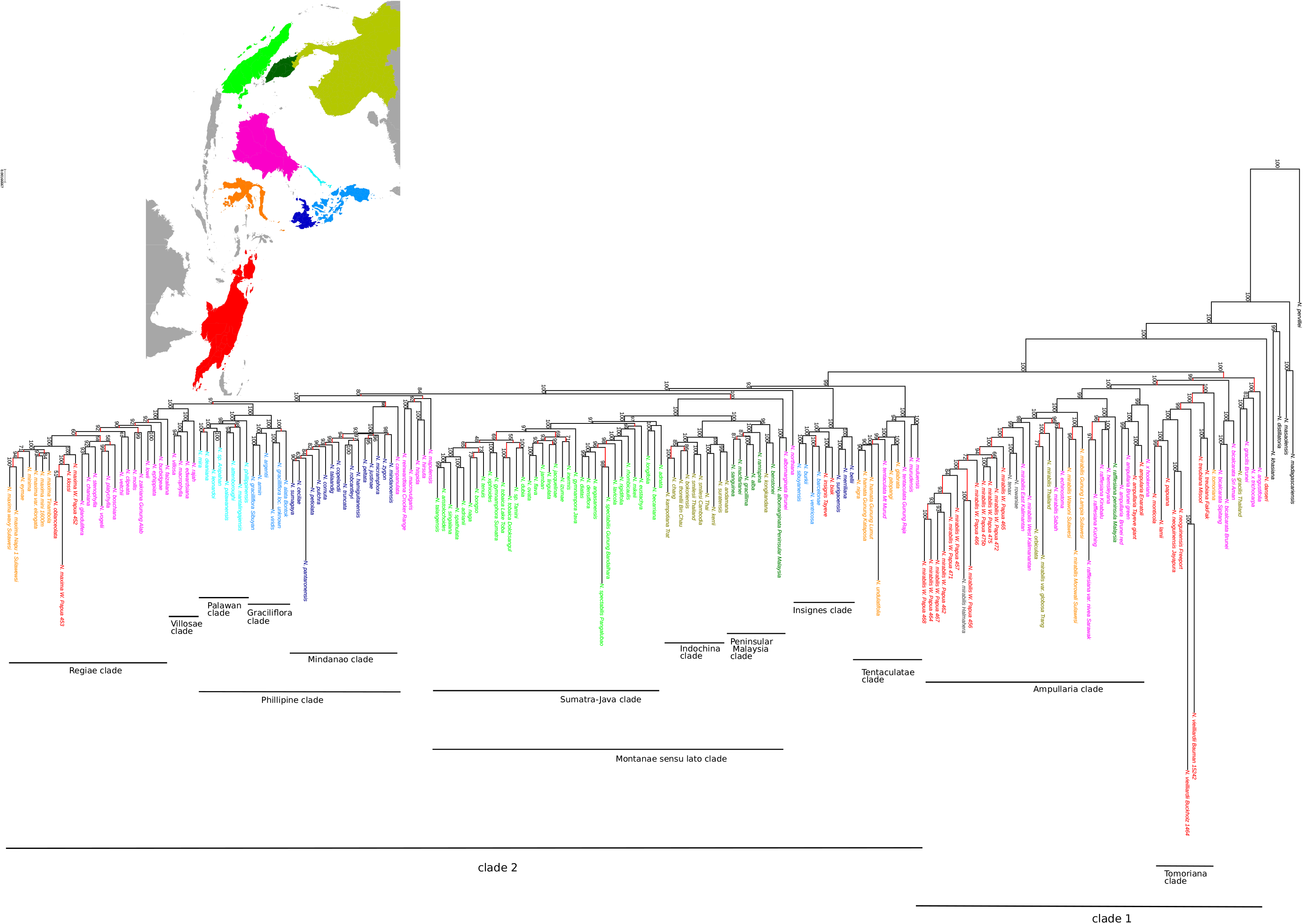
177-Supermatrix ML tree tree for ingroup taxa, generated with IQTREE partitioned analysis with Partitionfinder and Modelfinder option from 177-Supermatrix alignment of concatenated loci. Tree is rooted to *N. pervillei* based on results in Fig. 1. Support values are ultrafast bootstrap scores from 1000 replicates. Branches not recovered in the MQS-all-loci analysis (Fig. 2) are shown in red. Coloured tip names correspond to the accompanying map.

Concordance data (for the clade containing *N. vieillardii N. neoguineensis* and others (branch id 289), shows more genes supporting an alternative topology to the 177-supermatrix tree (GCF 0.45%, GDF2 = 4.93%), while site support (SCF 51.64%) is relatively good (Fig 3, S2, Table S4). This may explain why the MQS-all-loci tree (Fig. 2) shows a different topology here since this tree is based on gene trees but the supermatrix tree is not. Furthermore, the SCF data (Table S3, Fig. S1) for the MQS-all-loci tree (Fig. 2) shows the branch leading to the sister clade to *N. vieillardii* (branch id 247) has a SDF2 score (45.5%) considerably higher than the SCF score (28.7%), indicating more support for an alternative topology. While it may not be a surprise to see site data supporting a supermatrix tree rather than the gene tree based MQS tree, it is worth noting that, unlike the 177-supermatrix tree itself, the SCF scores should not be vulnerable to ILS because they are derived from quartet-trees which do not have an anomaly zone (Minh et al., 2018). Therefore, they provide additional evidence for the supermatrix topology.

### 3.8 Clade 2

Within Clade 2, major clades corresponding often to geographic areas are resolved consistently between trees, although the backbone topology is unstable in places and several Bornean taxa are inconsistently or uncertainly placed around these clades. Several of these taxa have been problematic in previous studies and taxonomic accounts, namely *N. macrovulgaris*, *N. campanulata, N. northiana, N. mapuluensis, N. reinwardtiana, N. albomarginata, N. hirsuta* and *N. hispida* (Alamsyah and Ito, 2013; Cheek and Jebb, 2001; Clarke et al., 2018a; Meimberg and Heubl, 2006; Mullins, 2000).

#### 3.8.1 *Tentaculatae* and *Insignes* clades

On all our trees (Figs. 2–4), two strongly supported clades representing sects. *Tentaculatae* and *Insignes* are resolved as successively sister to the rest of Clade 2 with maximal support. Both the *Tentaculatae* and *Insignes* clades were described as sections based on morphology (Cheek et al., 2018; Cheek and Jebb, 2016a; Danser, 1928). It is notable that these sections both range across Wallace’s line. They were supported by the phylogenies of Meimberg and Heubl (2006) and, less clearly, that of Mullins (2000) and Alamsyah and Ito (2013). We included in our study the previously unsampled species *N. biak, N. surigaoensis* and *N. barcelonae* from sect. *Insignes*, and *N. nigra, N. pitopangii* and *N. undulatifolia* from sect. *Tentaculatae*, confirming the previous morphologically-based sectional placement of these species (Cheek et al., 2018, 2015; Cheek and Jebb, 2016a; Nerz et al., 2011b, 2011a). The affinities of *N. undulatifolia* were considered unclear when the species was described (Nerz et al., 2011b) but was included in sect. *Tentaculatae* by Cheek and Jebb (2016a). Unfortunately we were unable to recover DNA from *N. maryae*, also of sect. *Tentaculatae*. We do not find any indication of a close relationship of sect. *Insignes* with *N. campanulata* as suggested by Clarke et al. (2018b) despite the variable placement of the latter species in our trees. *Nepenthes northiana* from Borneo was included with sect. *Insignes* by Danser but this position was considered doubtful (Cheek et al. 2018). Here (Figs. 2–3) it is resolved alone as sister to the remaining taxa of Clade 2 after the *Tentaculatae* and *Insignes* clades, although it is weakly supported in the MQS-all-loci tree. In the MQS-super-loci tree (Fig. 4) a clade of *N. northiana, N. hispida* and *N. hirsuta* is resolved in the same position.

**Fig. 4.**
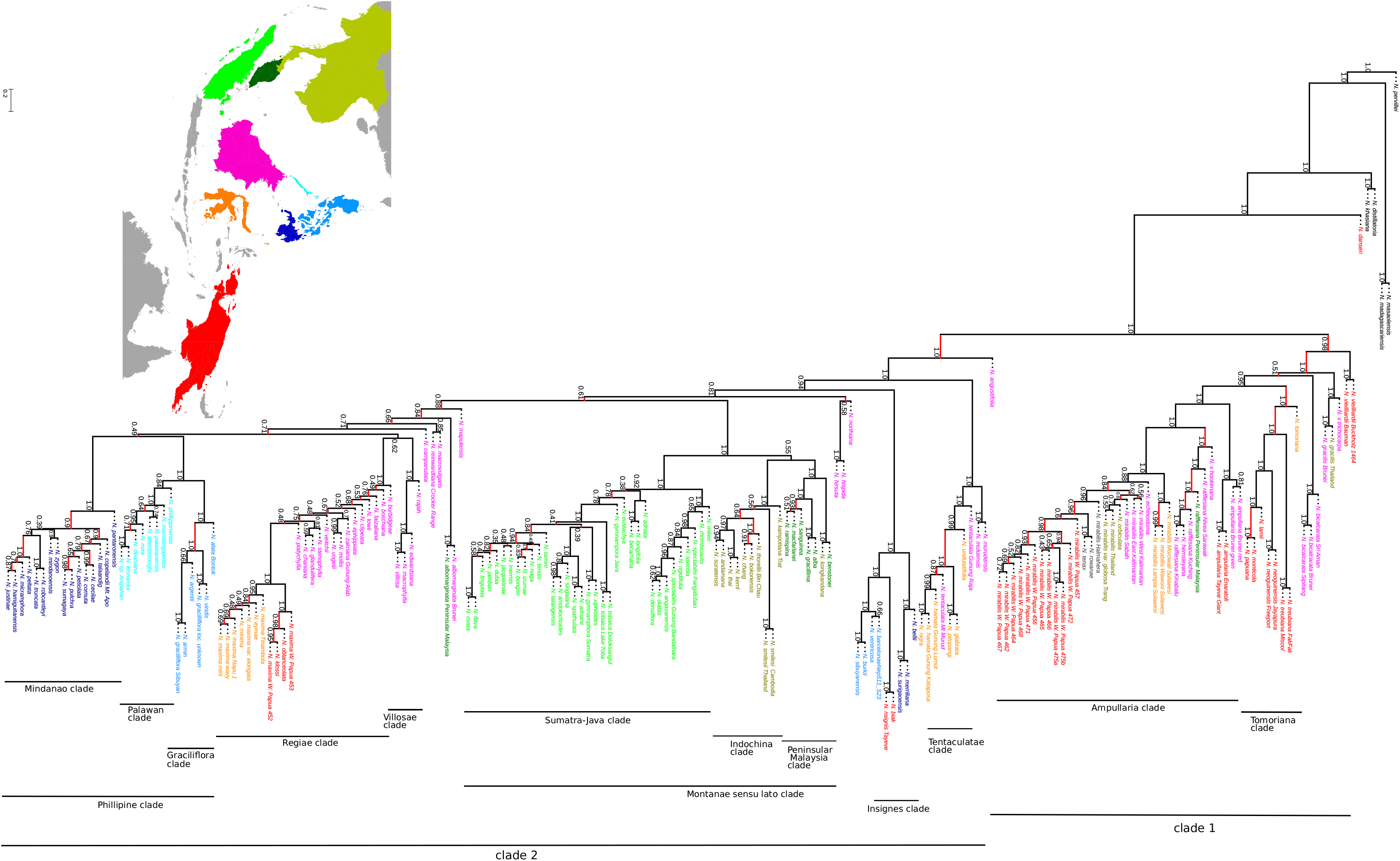
MQS-super-loci species tree for ingroup taxa generated with ASTRAL III based on individual ML partitioned gene-trees for 318 combined CDS-intron super-loci generated with IQTREE using the Partitionfinder and Modelfinder options. Tree is rooted to *N. pervillei* based on results in Fig. 1. Support values are posterior probabilty scores (ASTRAL “-t 3”) option. Branches not recovered in the 177-supermatrix ML analysis (Fig 3.) are shown in red. Coloured tip names correspond to the accompanying map.

#### 3.8.2 *Montanae sensu. lato* clade (Sumatra, Peninsular Malaysia and Indochina)

A large clade of Sumatran (and occasionally Javan) taxa is consistently resolved by our analyses, referred to here as sect. *Montanae sensu lato* (Figs. 2–4). Separate Peninsular Malaysia and Indochina clades are also consistently resolved as sister to each other and in turn to the Sumatra clade. Support for these clades is very good except for the sister pairing of the Indochinese and Peninsular Malaysian species on the super-loci tree (PP 0.55). Gene and site concordance, however, are low (Figs. 2, S1–2, Tables S3-4). While many of these species have been grouped together before on morphological grounds, the overall geographical sub-structure of this large clade is notable and was not previously as well defined as it is here. Danser (1928) and Cheek and Jebb (2009) grouped the mainly montane, forest, Sumatran species together with the morphologically similar Malaysian species in sect. *Montanae*. Mullins (2000) failed to recover the Sumatran Clade in either of his main trees although in his combined nuclear–plastid reduced tree with successive weighting the Sumatran clade and a separate Indochina–Peninsular Malaysia clade are recovered, similarly to here. Meimberg and Heubl (2006) also recovered a Sumatran clade in their plastid analysis. Alamsyah and Ito’s (2013) ITS topology is close to ours, recovering a Sumatran clade as sister to a clade containing mainly Indochinese–Peninsular–Malaysian taxa, but the presence of the wider-ranging (Borneo and Sumatra) *N. reinwardtiana* and Bornean *N. macrovulgaris* in the latter clade is not replicated here. Informed by these studies, Clarke et al. (2018b) restricted sect. *Montanae* to Sumatran taxa while adding all Indochinese and Peninsular Malaysian species to sect. *Pyrophytae*, although most are not known as pyrophytes.

The recovery in all our trees of *N. kongkandana* from Southern Thailand with the Peninsular Malaysian species is a minor exception to the geographical structure of these clades. The Kangar–Pattani Line, close to the Thai–Malaysia border is often considered the phyto-geographical boundary between Indochina and Sundaland (Whitmore, 1998), and *N. kongkandana* ranges from roughly this boundary north to the Khao Sok National Park 400km away (McPherson, 2009). This boundary is based on a perceived climatological boundary between ever-wet and seasonal rainfall patterns but Woodruff (2010) suggests the narrow isthmus of Kra, a more popular boundary for zoogeographers, may in fact better correspond with this climatological and phytogeographical transition. This would put *N. kongkandana* in the (Malaysian) Sundaland zone. However, this species does apparently experience seasonal drought and shares the pyrophytic adaptations of the Indochinese species (McPherson, 2009) so it would appear that these two clades contain a mixture of traits regardless of where the phytogeographical boundary is drawn. Similarly, *N. thai* and *N. bokorensis*, which were thought to be less seasonally adapted and treated by Cheek and Jebb (Cheek and Jebb, 2016b, 2009) as part of the Peninsular Malaysian *Montanae*, are resolved here with the pyrophytic Indochinese taxa. The latter of these does, in fact, occur in seasonal environments although it seems to be unknown whether it possesses pyrophytic adaptations (Mey, 2009). Since other unrelated taxa such as *N. minima* (Cheek and Jebb, 2016c) also appear to possess pyrophytic traits, these may be evolutionarily labile.

Based presumably on previous molecular studies, Clarke et al. (2018b) placed in sect. *Pyrophytae* several species that occur in Peninsular Malaysia and/or Sumatra but which are not endemic to these regions, namely *N. reinwardtiana, N. albomarginata* and *N. rafflesiana*, along with *N. hemsleyana* (endemic to Borneo but related to *N. rafflesiana)*. Our sampling included representatives of some of these taxa from this region but does not support this placement, instead resolving these taxa in Clade 1, except for *N. albomarginata* which is resolved as sister to the entire *Montanae sensu lato* clade on Figs. 3 and 4. It is possible, given Mullins’s (2000) results and the limited clarity of Meimberg and Heubl’s (2006) analyses, that plastid and nuclear nuclear signals may be strongly conflicting with regard to these species. However, we note that for other taxa *(N. maxima, N. neoguineensis, N. mapuluensis)* with conflicting positions between Mullins’s (2000) trees, the plastid phylogeny of Meimberg and Heubl (2006) agrees more with our nuclear one. Support is also fairly low in Mullins’s (2000) trees. We suggest apparent conflicts between plastid and nuclear signal in Mullins’s (2000) study may be based partly on the weakness of signal from single loci and may not be more significant than that between any of the hundreds of loci sequenced here. Full plastome sequencing would help clarify this issue.

#### 3.8.3 Philippine endemic clades and Sect. *Regiae*

Sister to the *Montanae sensu lato* Clade on our trees are three main clades, the combined *Regiae* plus *Villosae* clade, the Mindanao clade and the Palawan plus *N. graciliflora* clade. The relationship of these clades is resolved differently on our trees (Figs. 2–4). Both our MQS trees (Figs. 2, 4) recover all the Philippine taxa (except for the *Insignes* taxa discussed above) as a clade. Within this, the Palawan Clade and the *N. graciliflora* Clade together are sister to the well supported Mindanao Clade. This large Philippine Clade is sister in turn to a clade consisting of the small *Villosae* Clade, which is then sister to the large *Regiae* Clade, with *N. maxima* and its allies forming a sub-clade nested within the other Bornean *Regiae* taxa. However, the 177-supermatrix tree (Fig. 3) instead resolves the *Regiae* Clade as sister to the combined Palawan and *N. graciliflora* clades, with this combined clade in turn sister to the Mindanao Clade and *N. campanulata*.

Previous studies have not sampled the *non-Insignes* Philippine taxa in great depth. Mullins (2000) does resolve separate Mindanaoan and Palawan–Sibuyan clades on both trees but with few samples and inconsistent placement. In the absence of clarity from phylogenetic studies, classification of the Philippine taxa has been controversial (Clarke et al. 2018b). Cheek and Jebb (2015) have placed taxa in three main groups in addition to sect. *Insignes:* sects. *Villosae, Micramphorae* and *Alatae*. Their *Villosae* contained, in addition to *N. rajah* and the core group of species with dramatically ridged peristomes from northern Borneo, most of the Palawan taxa and *N. argentii* from Sibuyan. Clarke et al. (2018b), following Robinson et al. (2008), instead group all Palawan species with the *Regiae*, along with the large-pitchered Mindanaoan species *N. peltata*. Here, we consistently resolve *N. rajah* with the three Bornean species of the *Villosae sensu stricto*, but do not recover the Palawan taxa within this clade. Instead, the *Villosae* and *Regiae* are resolved as sister clades on all our trees although with little gene concordance and poor support on the MQS species trees (Figs. 2, 4).

Sect. *Micramphorae* was established by Cheek and Jebb (2015) to accommodate *N. micramphora, N. cid* and *N. abgracilis*. Only the former is sampled here and is recovered with other Mindanaoan taxa. We predict the other taxa of this section would also be resolved within this well-supported and geographically homogeneous Mindanao clade. However, Cheek and Jebb (2015, 2014, 2013b), placed most other Mindanaoan species in their large sect. *Alatae*, along with *N. armin* from Sibuyan, *N. alata* and *N. ultra* from Luzon, two toponymous species from the islands of Leyte and Negros (not sampled here) and the widespread *N. graciliflora*. In contrast, Clarke et al. (2018b) place most of the *Alatae* with Bornean species in sect. *Villosae* and sect. *Regiae*. Although our trees group the Bornean and Philippine clades together, and the proximity of Palawan in particular makes a connection between Borneo and the Philippines plausible (Cheek, 2011), in all our trees both the Palawan and Mindanao taxa form separate, strongly-supported geographical clades expanding those recovered by Mullins (2000). The sister relationship between the Palawan clade and the mixed Philippine *N. graciliflora* clade is also well-supported and consistent. However, this finding complicates taxonomic classification by separating *N. alata*, the type species of sect *Alatae*, from most of the taxa formerly in this section. Geographical distribution again appears to be a better guide to relationships than morphology, although several Philippine taxa remain unsampled. Unfortunately, the backbone topology connecting these clades is uncertain because of inconsistency between the supermatrix and MQS trees and low support. Although our MQS-super-loci tree (Fig. 4) recovers the same topology as the MQS-all-loci species tree (Fig. 2), concordance factors show zero genetrees recovering either this plausible pan-Philippine clade or the larger Regiae+Philippine clade.

The clade of *N. maxima* and allies from Sulawesi and New Guinea is nested within the Bornean *Regiae* clade in all our trees and represents a single crossing of Wallace’s line. Several of the Bornean species have previously been considered close relatives of *N. maxima* and to form a *N. maxima* group, so this topology is expected (Jebb and Cheek, 1997; Robinson et al., 2011). The New Guinean sub-clade of the *Regiae* is further discussed in section 3.11 below.

### 3.10 Other unstable taxa

A number of other Bornean taxa are inconsistently placed within Clade 2: *N. campanulata, N. reinwardtiana, N. macrovulgaris, N. albomarginata, N. mapuluensis* and the sister clade of *N. hirsuta* and *N. hispida*. Several of these taxa are known to occur on ultramafic or limestone substrates (Cheek and Jebb, 2001). Our trees do agree about a sister relationship between *N. macrovulgaris* and *N. reinwardtiana* and a close relationship between these species was also found in the plastid trees of Meimberg et al. (2000) and Mullins (2000) and the ITS tree of Alamsyah and Ito (2013). *Nepenthes hispida* and *N. hirsuta* are sister to this pair on our supermatrix tree (Fig. 3) and are usually considered morphologically close to *N. macrovulgaris* (Cheek and Jebb, 2001; McPherson, 2009) but our MQS trees (Figs. 2,4) do not recover this relationship. Our trees also agree to some extent in placing *N. albomarginata* close to the combined *Montanae sensu lato* Clade. *Nepenthes mapuluensis* is morphologically similar to *N. northiana* and both occur on limestone. However, these taxa are not shown to be closely related to each other by any of our trees, or by any previous molecular studies.

### 3.11 Taxa with multiple samples

Where possible, we tried to sample multiple individuals of geographically wide-ranging taxa, an approach recommended for detecting incipient speciation or ancestral polymorphism manifested as non-monophyletic species clades (Pennington and Lavin, 2016). Given the extent of infra-specific morphological variation and inter-specific hybridisation within the genus, it was hypothesised that our same-species samples might not always form clades. This is indeed the case, but interpretation is difficult due to low gene and site concordance and noise. Multiple samples of the widespread and variable *N. maxima* are resolved according to their origin but the species as a whole is rendered paraphyletic by *N. eymae, N. klossii, N. minima*, and *N. oblanceolata*. Although all are considered related to *N. maxima*, the first two species in particular have very distinct pitchers, as well as differences in ecology and altitudinal distribution (McPherson, 2009). Few authorities would dispute their species status. Within the *Tentaculatae* clade, although we were not able to sample *N. tentaculata* from Sulawesi where it is also found, multiple samples each of *N. tentaculata* and *N. hamata* were not monophyletic on our trees. Poor recovery for *N. undulatifolia* may contribute to inaccurate topology in this part of the tree. However, *N. tentaculata* is a very common species and therefore non-monophyly could be caused by the long coalescence time due to a large effective population size (Pennington and Lavin, 2016). Being strictly dioecious and therefore obligate out-crossers, *Nepenthes* also have a larger effective population size than selfing species.

As the widest-ranging and likely most abundant species, we sampled *N. mirabilis* more than any other species. Our clade of *N. mirabilis* samples also includes several taxa recently segregated as separate species *(N. orbiculata, N. tenax, N. rowaniae)* as well as other notable variants sometimes considered species *(N. echinostoma* and *N. mirabilis var. globosa)*. Recognising these as separate species renders *N. mirabilis* paraphyletic. However, this is probably not an unusual taxonomic problem and some of the specific issues relating to *N. mirabilis* are discussed by Clarke et al. (2009). Despite this obvious paraphyly, when we consider the very large range, lack of obvious long-range dispersal mechanisms and propensity for hybridisation of *N. mirabilis* (McPherson, 2009), it is notable that these related species do form a clade. Further paraphyly of some of our subclades within *N. mirabilis* are probably partly due to error caused by noise or lack of resolution and support in the individual gene trees. For instance, the placement by our MQS-all-loci tree (Fig. 2) of the two new Australian taxa, *N. rowaniae* and *N. tenax*, within a clade of New Guinea individuals sampled from a very small population should be treated with scepticism. Similarly, the recovery of a sample of *N. mirabilis* from Halmahera within this clade on our 177-supermatrix tree (Fig. 3) is probably related to the poor recovery and long branch for this sample.

For *N. ampullaria, N. gracilis* (with the exception of *N. angustifolia* discussed above*), N. bicalcarata, N. treubiana, N. neoguineensis, N. smilesii, N. vieillardii* and *N. tobaica*, multiple samples were resolved as clades. *Nepenthes rafflesiana* was rendered paraphyletic by *N. hemsleyana*. Although poor recovery might be responsible for this finding, it is also possible that closely located populations of *N. hemsleyana* and *N. rafflesiana* are more closely related than samples of the latter species from across its range. The two recently separated species *N. insignis* and *N. biak* were also grouped together, as were the morphologically very similar *N. hispida* and *N. hirsuta*. The two *Nepenthes graciliflora* samples were part of the same small clade but not sisters, although SCF scores suggest a possible alternative topology within this clade. *N. graciliflora* is known to be a wide-ranging and varied taxon (Cheek and Jebb, 2013c) so non-monophyly of samples would not be surprising. The Javan and Sumatran samples of *N. gymnamphora* were similarly close but not sisters, as were the two samples of *N. hamata* which formed a clade with *N. nigra*.

### 3.11 Gene conflict

Important differences between our trees have been noted above and make the backbone topology unclear in places, mainly regarding relationships in Clade 2 between the *Regiae, Montanae sensu lato* and Philippine clades and the unstable Bornean taxa. These inconsistencies hint at extensive conflict that is hidden by the high or maximal BS and LPP support values in our trees. It is widely accepted that bootstraps in particular can mask incorrect topology and gene tree conflict (Xi et al., 2014). The GCF scores for our trees (Tables S3–4, Figs. S1–2) typically indicate very low gene concordance for most branches. SCF values (Figs. 2, S1–2, Tables S3–4) are somewhat higher but it should be noted that because these are quartet-based, only a score above 33% is indicative of the given topology being more prevalent than the alternatives (Minh et al., 2018). However, for both GCF and SCF values, there are relatively few cases of an alternative topology being strongly more favoured, which suggests generally weak phylogenetic signal.

Quartet methods such as ASTRAL, which is used here, have been proven to be statistically consistent under the multi-species coalescent as long as the input gene trees are true, and can potentially produce the correct topology when supermatrix approaches are misled by ILS (Mirarab et al., 2014). However, these methods have been criticised because in practise they can be undermined by poorly resolved, inaccurate gene trees (Gatesy and Springer, 2014). Our MQS-all-loci gene trees had an overall mean BS value of 57 across all nodes (Table S2). After collapsing nodes with bootstrap support lower than 10 prior to generating our MQS species tree, the average bootstrap support rose to 68. ASTRAL does not take support values into account when estimating the species tree and therefore poorly resolved gene trees could give a misleading impression of gene discordance, as well as an inaccurate species tree. SCF scores are not dependent on genetrees and for some key nodes, our SCF scores do provide greater confidence. For, example, in the MQS-all-loci analysis (Fig. 2) the Sumatran clade (branch id 360) is supported by just 1.07% (i.e. 7) of loci but the SCF is more encouraging (47.6% versus 27.2% and 25.2%). This helps explains why the 177-supermatrix tree (Fig. 4) is able to clearly resolve this clade. Furthermore, the SCF scores provide further confidence because, being based on quartets, they are not vulnerable to ILS in the same way that the supermatrix tree could be.

In general, there are few instances were SCF scores strongly contradict the locus-based topology of the MQS-all-loci tree (Fig. 2), indicating that while the signal from individual loci is weak it does not generally distort the signal from individual sites. One exception is at internode 232 which places *N. rowaniae* and *N. tenax* within the clade of Papuan *N. mirabilis* samples on our MQS species tree (Figs. 2, S1;Table S3). Here there is a score of 31.93% for the MQS-all-loci topology (Fig. 2) versus 47.35% for an alternative topology, suggesting the alternative topology of the 177-supermatrix tree (Fig. 3) may be more likely. Interestingly, our MQS-super-loci tree (Fig. 4) agrees with the 177-supermatrix topology here. This may be because the super-loci tree is produced from loci with greater signal (hence a higher average bootstrap value of 78) and therefore in cases where the CDS loci are unreliable, it is more likely to replicate the topology of the supermatrix tree. Despite this, the topology for this tree is for the most part the same as the MQS species tree (Fig. 2).

SCF values, however, are also generally quite low suggesting considerable noise or conflict in the signal not just between loci but between individual sites. Our 177-supermatrix tree (Fig. 3) tends to show long terminal branches throughout the tree, suggesting a large number of variable sites are not parsimony-informative. This applies also to our population samples of *N. mirabilis* from West Papua which were taken from sites very close together and showed no obvious morphological variation. These samples do form a clade on our supermatrix tree but have terminal branch lengths indicating a large amount of variation, comparable to that between species. In fact, our technical replicate samples of *N. mirabilis* 475 also show non-negligible branch lengths. As mentioned, further analysis of these samples revealed a considerable number of CDS loci showing variation between these replicates. Future work should address this problem of noise in order to better utilise the Angiosperms-353 probe set at the species and population level. We suggest that a species-specific custom target-file may help avoid non-homologous sequences being added to alignments. An automated method for checking and removing spurious, non-aligned sections of individual sequences from alignments without removing the columns from other sequences might also be beneficial if an appropriate threshold could be found.

### 3.12 Consequences for infrageneric classification

Our findings suggest a number of modifications to the sections suggested by Danser (1928), Cheek and Jebb (2016b, 2015, 2009) and Clarke et al. (2018b). Most of the common, lowland, widespread species are consistently resolved together in Clade 1, suggesting that either sect. *Urceolatae* or sect. *Nepenthes (Vulgatae)* might be expanded to correspond with this clade. *N. rafflesiana* and *N. hemsleyana*, which were grouped in sect. *Pyrophytae* by Clarke et al. (2018b), should also be included here according to our data, which agrees in this respect with the 5S-NTS nuclear tree of Mullins (2000). The *N. tomoriana* clade containing mainly species from New Guinea could become a subsection within this section, although it is unclear whether *N. vieillardii* belongs here. Contrary to Clarke et al. (2018b), our results suggest *N. papuana* should be included in this section and we would also expect the paniculate species *N. paniculata* to belong here. Our data also supports an exclusively Sumatran–Javan sect. *Montanae*. However, considering the high gene and site conflict, it may be wiser to retain all Peninsular Malaysian and Indochinese species within the *Montanae sensu lato* and treat each of the Sumatra–Java, Indochina *(Pyrophytae)* and Peninsular Malaysia clades as subsections.

The Mindanaoan clade would make an obvious section. Unfortunately, most of these species have been attributed to sect. *Alatae* in the past and the type species of this section, *N. alata*, is clearly not part of this group according to our data. The well supported clade containing all the Palawan species may merit sectional status, with or without the rump of sect. *Alatae* (our *N. graciliflora* Clade) which forms its sister clade. The geographically heterogeneous, morphologically-defined sections *Tentaculatae* and *Insignes* are also strongly confirmed, while our data suggest sect. *Regiae* should be reduced to a much smaller set of taxa than that suggested by Clarke et al. (2018b). Sect. *Villosae* is supported in the narrow sense of the three morphologically and geographically close Bornean species plus the less morphologically obvious *N. rajah*. It is unclear whether this group is best considered a subsection of the *Regiae* or a separate section*. Nepenthes maxima* and its New Guinean and Sulawesian allies might be considered a subsection within the *Regiae* although it is unclear from our results which (if any) of the Bornean species resembling *N. maxima* should be included. The remaining Bornean taxa *N. campanulata, N. macrovulgaris, N. mapuluensis, N. hirsuta, N. hispida, N. albomarginata* and *N. reinwardtiana* cannot yet confidently be placed within any of these sections although they are clearly resolved in Clade 2.

## 4. Conclusions

Our use of phylogenomic sequencing resolves most species into well supported clades and reveals new topological detail about the evolution of lineages. Gene conflict is extensive but for the most part does not prevent a confident assessment of relationships, as indicated by the broad agreement between trees produced by different methods and from different datasets. We confirm previous findings regarding the early diverging Western taxa and are able to confirm *N. pervillei* from the Seychelles as sister to the rest of the genus and *N. danseri* from Waigeo island as sister to the remaining species. We find that several clades are strongly geographically delimited, particularly species from Sumatra, Mindanao and Palawan. Taxa from Borneo are separated into multiple clades, suggesting multiple migrations into or out of Borneo, but a number of Bornean taxa are not clearly resolved, posing an obstacle to further interpretation. Overall, our findings also suggest at least seven dispersals across Wallace’s line. Our sampling of multiple individuals of several widespread taxa helps add confidence that these are correctly considered single species, although the extent of gene flow between different populations is unknown. The segregation of several taxa from *N. mirabilis* and *N. maxima* result in these taxa being non-monophyletic, raising questions about the recognition of paraphyletic taxa. Whilst some concerns remain over noise and resolution of individual gene trees, our results show the potential for using the Angiosperms-353 probe set for species level systematics.

## Acknowledgements

This work was funded by the Natural Environment Research Council (NERC) Science and Solutions for a Changing Planet Doctoral Training Partnership. Further funding from the Emily Holmes Foundation is gratefully acknowledged. JR was funded an Individual Research Fellowship from NERC (NE/L011611/1). Through TB and JR this study is a contribution to Imperial College’s Grand Challenges in Ecosystems and the Environment initiative.

We would like to thank Chester Zoo, in particular Paul Leach, Richard Hewitt and Philip Esseen, for allowing extensive sampling from the National Nepenthes Collection. We are also grateful to RBG Kew for samples from their living collection and to Rebecca Hilgenhoff, Jess Snowball, Tom Pickering and Lara Jewitt for their assistance. We also thank Andreas Wistuba for kindly providing samples from his nursery, Mr Yanto for samples from his private collection in Java, the Herbarium at Kew for allowing sampling of herbarium specimens and Xander van der Burgt for additional samples.

Fieldwork in West Papua was carried out under RISTEK permit number 1161/SIP/FRP/E5/ Dit.KI/VI/2017. We gratefully acknowledge the assistance of Freddy Pattiselano and his colleagues at the Biodiversity Research Center, UNIPA, Manokwari, Jimmy Wanma at UNIPA and Muhammad Mansur and Lina Juswara at the Indonesian Institute of Sciences, Jakarata-Bogor for assistance collecting and processing samples from West Papua. Charlie Heatubun (Balitbangda Provinsi Papua Barat) is especially thanked for inspiring our field visit to Indonesia.

From RBG Kew, we additionally thank Laszlo Csiba, Grace Brewer and Niroshini Epitawalage for laboratory assistance and Jan Kim for bioinformatics advice. We acknowledge the use of the high performance cluster and support of the HPC team at Imperial College, London. Finally, we are grateful to the PAFTOL team at RBG Kew, in particular Lisa Pokorny, Steven Dodsworth and Olivier Maurin, and the 1KP project for providing sequenced samples and for early use of the Angiosperms-353 gene probe set.

