## Appendix 1 for "A phylogenomic analysis of *Nepenthes* (Nepenthaceae)"

| **Species** | **Authority** | **Year** | **Biogeographical area** | **Danser (1928)** | **Cheek & Jebb** | **Clarke et al., (2018)** |
| --- | --- | --- | --- | --- | --- | --- |
| *Nepenthes abalata* | Jebb & Cheek | 2013 | Phillipines |  | Pyrophytae? | Regiae |
| *Nepenthes abgracilis* | Jebb & Cheek | 2013 | Phillipines |  | Micramphora | Villosae |
| *Nepenthes adnata* | Tamin & M.Hotta ex Schlauer | 1994 | Sumatra |  | Tentaculatae | Montanae |
| *Nepenthes aenigma* | Nuytemans, W.Suarez & Calaramo | 2016 | Phillipines |  | Insignes | Insignes |
| *Nepenthes alata* | Blanco | 1837 | Phillipines | Vulgatae | Alatae | Regiae |
| *Nepenthes alba* | Ridl. | 1924 | Peninsular Malaysia |  | Montanae | Pyrophytae |
| *Nepenthes albomarginata* | T.Lobb ex Lindl. | 1849 | widespread | Vulgatae |  | Pyrophytae |
| *Nepenthes alzapan* | Jebb & Cheek | 2013 | Phillipines |  | Insignes | Insignes |
| *Nepenthes ampullaria* | Jack | 1835 | widespread | Urceolate |  | Urceolatae |
| *Nepenthes andamana* | M.Catal. | 2010 | Indochina |  | Pyrophatae | Pyrophytae |
| *Nepenthes angasanensis* | Maulder, D.Schub., B.R.Salmon & B.Quinn | 1999 | Sumatra |  | Montanae | Montanae |
| *Nepenthes appendiculata* | Chi.C.Lee, Bourke, Rembold, W.Taylor & S.T.Yeo | 2011 | Borneo |  |  | Regiae |
| *Nepenthes argentii* | Jebb & Cheek | 1997 | Phillipines |  | Villosae | Regiae |
| *Nepenthes aristolochioides* | Jebb & Cheek | 1997 | Sumatra |  | Montanae | Montanae |
| *Nepenthes armin* | Jebb & Cheek | 2014 | Phillipines |  | Alatae | Regiae |
| *Nepenthes attenboroughii* | A.S.Rob., S.McPherson & V.B.Heinrich | 2009 | Phillipines |  | Villosae | Regiae |
| *Nepenthes barcelonae* | Tandang & Cheek | 2015 | Phillipines |  | Insignes | Insignes |
| *Nepenthes beccariana* | Macfarl. | 1908 | Sumatra |  | Montanae |  |
| *Nepenthes bellii* | K.Kondo | 1969 | Phillipines |  | Insignes | Insignes |
| *Nepenthes benstonei* | C.Clarke | 1999 | Peninsular Malaysia |  | Montanae | Pyrophytae |
| Nepenthes biak | Cheek & Jebb | 2018 | New Guinea |  | Insignes |  |
| *Nepenthes bicalcarata* | Hook.f. | 1873 | Borneo | Urceolate |  | Urceolatae |
| *Nepenthes bokorensis* | Mey | 2009 | Indochina |  | Montanae | Pyrophytae |
| *Nepenthes bongso* | Korth. | 1839 | Sumatra | Montanae | Montanae | Montanae |
| *Nepenthes boschiana* | Korth. | 1839 | Borneo | Regiae | Regiae | Regiae |
| *Nepenthes burbidgeae* | Hook.f. ex Burb. | 1882 | Borneo | Regiae | Regiae | Regiae |
| *Nepenthes burkei* | Hort.Veitch ex Mast. | 1889 | Phillipines | Insignes | Insignes | Insignes |
| *Nepenthes campanulata* | Sh.Kurata | 1973 | Borneo |  |  | Insignes |
| *Nepenthes carunculata* | Danser | 1928 | Sumatra | Montanae | Montanae |  |
| *Nepenthes ceciliae* | Gronem., Coritico, Micheler, Marwinski, Acil & V.B.Amoroso | 2011 | Phillipines |  | Alatae | Villosae |
| *Nepenthes chang* | M.Catal. | 2010 | Indochina |  | Pyrophytae | Pyrophytae |
| *Nepenthes chaniana* | C.Clarke, Chi.C.Lee & S.McPherson | 2006 | Borneo |  | Regiae | Regiae |
| *Nepenthes cid* | Jebb & Cheek | 2013 | Phillipines |  | Micramphora | Regiae |
| *Nepenthes clipeata* | Danser | 1928 | Borneo | Regiae | Regiae | Regiae |
| *Nepenthes copelandii* | Merr. ex Macfarl. | 1908 | Phillipines |  | Alatae | Villosae |
| *Nepenthes cornuta* | Marwinski, Coritico, Wistuba, Micheler, Gronem., Gieray & V.B.Amoroso | 2014 | Phillipines |  | Alatae | Villosae |
| Nepenthes dactylifera | A.S.Rob., Golos, S.McPherson & Barer | 2019 | Borneo |  |  |  |
| *Nepenthes danseri* | Jebb & Cheek | 1997 | Waigeo |  | Danseri group | Nepenthes |
| *Nepenthes deaniana* | Macfarl. | 1908 | Phillipines | Nobiles | Villosae | Regiae |
| *Nepenthes densiflora* | Danser | 1940 | Sumatra |  | Montanae | Montanae |
| *Nepenthes diatas* | Jebb & Cheek | 1997 | Sumatra |  | Montanae | Montanae |
| *Nepenthes distillatoria* | L. | 1753 | Sri Lanka | Vulgatae |  | Nepenthes |
| *Nepenthes dubia* | Danser | 1928 | Sumatra | Montanae | Montanae | Montanae |
| *Nepenthes echinostoma* | Hook.f. | 1873 | Borneo | synononymn mirabilis | synononymn mirabilis | synononymn mirabilis |
| *Nepenthes edwardsiana* | H.Low ex Hook.f. | 1859 | Borneo |  | Villosae | Villosae |
| *Nepenthes ephippiata* | Danser | 1928 | Borneo | Regiae | Regiae | Regiae |
| *Nepenthes epiphytica* | A.S.Rob., Nerz & Wistuba | 2011 | Borneo |  | Regiae | Regiae |
| *Nepenthes eustachya* | Miq. | 1858 | Sumatra |  | Montanae | Montanae |
| *Nepenthes extincta* | Jebb & Cheek | 2013 | Phillipines |  | Alatae |  |
| *Nepenthes eymae* | Sh.Kurata | 1984 | Wallacea |  | Regiae | Regiae |
| *Nepenthes faizaliana* | J.H.Adam & Wilcock | 1991 | Borneo |  | Regiae | Regiae |
| *Nepenthes fallax* | Beck | 1895 | Borneo |  | Regiae | Regiae |
| *Nepenthes flava* | Wistuba, Nerz & A.Fleischm. | 2007 | Sumatra |  | Montanae | Montanae |
| *Nepenthes fusca* | Danser | 1928 | Borneo | Regiae | Regiae | Regiae |
| *Nepenthes gantungensis* | S.McPherson, Cervancia, Chi.C.Lee, Jaunzems, Mey & A.S.Rob. | 2010 | Phillipines |  | Villosae | Regiae |
| *Nepenthes glabrata* | J.R.Turnbull & A.T.Middleton | 1984 | Wallacea |  | Tentaculatae | Tentaculatae |
| *Nepenthes glandulifera* | Chi.C.Lee | 2004 | Borneo |  | Regiae | Regiae |
| *Nepenthes graciliflora* | Elmer | 1912 | Phillipines |  | Alatae | Regiae |
| *Nepenthes gracilis* | Korth. | 1839 | widespread | Vulgatae |  | Urceolatae |
| *Nepenthes gracillima* | Ridl. | 1908 | Peninsular Malaysia | Montanae | Montanae | Pyrophytae |
| *Nepenthes gymnamphora* | Reinw. ex Nees | 1824 | Java (Sumatra) | Montanae | Montanae | Montanae |
| *Nepenthes halmahera* | Cheek | 2015 | Wallacea |  | Danseri group | Nepenthes |
| *Nepenthes hamata* | J.R.Turnbull & A.T.Middleton | 1984 | Wallacea |  | Tentaculatae | Tentaculatae |
| *Nepenthes hamiguitanensis* | Gronem., Wistuba, V.B.Heinrich, S.McPherson, Mey & V.B.Amoroso | 2010 | Phillipines |  | Alatae | Villosae |
| *Nepenthes hemsleyana* | Macfarl. | 1908 | Borneo |  |  | Pyrophytae |
| *Nepenthes hirsuta* | Hook.f. | 1873 | Borneo | Nobiles | Nobiles |  |
| *Nepenthes hispida* | Beck | 1895 | Borneo |  | Nobiles |  |
| *Nepenthes holdenii* | Mey | 2010 | Indochina |  | Pyrophatae | Pyrophytae |
| *Nepenthes hurrelliana(mollis)* | Cheek & A.L.Lamb | 2003 | Borneo |  | Regiae | Regiae |
| *Nepenthes inermis* | Danser | 1928 | Sumatra | Montanae | Montanae | Montanae |
| *Nepenthes insignis* | Danser | 1928 | New Guinea | Insignes | Insignes | Insignes |
| *Nepenthes izumiae* | Troy Davis, C.Clarke & Tamin | 2003 | Sumatra |  | Montanae | Montanae |
| *Nepenthes jacquelineae* | C.Clarke, Troy Davis & Tamin | 2001 | Sumatra |  | Montanae | Montanae |
| *Nepenthes jamban* | Chi.C.Lee, Hernawati & Akhriadi | 2006 | Sumatra |  | Montanae | Montanae |
| *Nepenthes justinae* | Gronem., Wistuba, Mey & V.B.Amoroso | 2016 | Phillipines |  | Alatae | Villosae |
| *Nepenthes kampotiana* | Lecomte | 1909 | Indochina | Vulgatae | Pyrophatae | Pyrophytae |
| *Nepenthes kerrii* | M.Catal. & Kruetr. | 2010 | Indochina |  | Pyrophatae | Pyrophytae |
| *Nepenthes khasiana* | Hook.f. | 1873 | Northern India | Vulgatae |  | Nepenthes |
| *Nepenthes kitanglad* | Jebb & Cheek | 2013 | Phillipines |  | Alatae | Regiae |
| *Nepenthes klossii* | Ridl. | 1916 | New Guinea | Regiae | Regiae | Regiae |
| *Nepenthes kongkandana* | M.Catal. & Kruetr. | 2015 | Indochina |  |  | Pyrophytae |
| *Nepenthes krabiensis* | Nuanlaong, Onsanit, Chu- | 2016 | Thailand |  |  | Pyrophytae |
| *Nepenthes lamii* | Jebb & Cheek | 1997 | New Guinea |  |  | Nepenthes |
| *Nepenthes lavicola* | Wistuba & Rischer | 1996 | Sumatra |  | Montanae | Montanae |
| *Nepenthes leonardoi* | S.McPherson, Bourke, Cervancia, Jaunzems & A.S.Rob. | 2011 | Phillipines |  | Villosae | Regiae |
| *Nepenthes leyte* | Jebb & Cheek | 2013 | Phillipines |  | Alatae | Regiae |
| *Nepenthes lingulata* | Chi.C.Lee, Hernawati & Akhriadi | 2006 | Sumatra |  | Montanae | Montanae |
| *Nepenthes longifolia* | Nerz & Wistuba | 1994 | Sumatra |  | Montanae | Montanae |
| *Nepenthes lowii* | Hook.f. | 1859 | Borneo | Regiae | Regiae | Regiae |
| *Nepenthes macfarlanei* | Hemsl. | 1905 | Peninsular Malaysia | Montanae | Montanae | Pyrophytae |
| *Nepenthes macrophylla* | (Marabini) Jebb & Cheek | 1997 | Borneo |  | Villosae | Villosae |
| *Nepenthes macrovulgaris* | J.R.Turnbull & A.T.Middleton | 1988 | Borneo |  | Nobiles |  |
| *Nepenthes madagascariensis* | Poir. | 1797 | Madagascar | Vulgatae |  | Nepenthes |
| *Nepenthes mantalingajanensis* | Nerz & Wistuba | 2007 | Phillipines |  | Villosae | Regiae |
| *Nepenthes mapuluensis* | J.H.Adam & Wilcock | 1990 | Borneo |  |  |  |
| *Nepenthes maryae* | Jebb & Cheek | 2016 | Wallacea |  | Tentaculatae | Tentaculatae |
| *Nepenthes masoalensis* | Schmid-Hollinger | 1977 | Madagascar |  |  | Nepenthes |
| *Nepenthes maxima* | Reinw. ex Nees | 1824 | Wallacea, New Guinea | Regiae | Regiae | Regiae |
| *Nepenthes merrilliana* | Macfarl. | 1911 | Phillipines | Insignes | Insignes | Insignes |
| *Nepenthes micramphora* | V.B.Heinrich, S.McPherson, Gronem. & V.B.Amoroso | 2009 | Phillipines |  | Micramphora | Villosae |
| *Nepenthes mikei* | B.R.Salmon & Maulder | 1995 | Sumatra |  | Montanae | Montanae |
| *Nepenthes mindanaoensis* | Sh.Kurata | 2001 | Phillipines |  | Alatae | Villosae |
| *Nepenthes minima* | Cheek & Jebb | 2016 | Wallacea |  | Regiae |  |
| *Nepenthes mira* | Jebb & Cheek | 1998 | Phillipines |  | Villosae | Regiae |
| *Nepenthes mirabilis* | (Lour.) Rafarin | 1869 | widespread | Vulgatae |  | Urceolatae |
| *Nepenthes mollis* | Danser | 1928 | Borneo | Regiae |  | Regiae |
| *Nepenthes monticola* | A.S.Rob., Wistuba, Nerz, M.Mansur & S.McPherson | 2011 | New Guinea |  |  | Nepenthes |
| *Nepenthes muluensis* | M.Hotta | 1966 | Borneo |  | Tentaculatae | Tentaculatae |
| *Nepenthes murudensis* | Culham ex Jebb & Cheek | 1997 | Borneo |  | Tentaculatae | Tentaculatae |
| *Nepenthes naga* | Akhriadi, Hernawati, Primaldhi & M.Hambali | 2009 | Sumatra |  | Montanae | Montanae |
| Nepenthes naquiyudddinii | Adam & Wilcock | 2006 | Borneo |  |  |  |
| *Nepenthes negros* | Jebb & Cheek | 2013 | Phillipines |  | Alatae | Regiae |
| *Nepenthes neoguineensis* | Macfarl. | 1911 | New Guinea | Vulgatae |  | Nepenthes |
| *Nepenthes nigra* | Nerz, Wistuba, Chi.C.Lee, Bourke, U.Zimm. & S.McPherson | 2011 | Wallacea |  | Tentaculatae | Tentaculatae |
| *Nepenthes northiana* | Hook.f. | 1881 | Borneo | Insignes | Insignes? |  |
| Nepenthes orbiculata | M.Catalana & Trong Kruetreepradtit | 2018 | Thailand |  |  |  |
| *Nepenthes ovata* | Nerz & Wistuba | 1994 | Sumatra |  | Montanae | Montanae |
| *Nepenthes palawanensis* | S.McPherson, Cervancia, Chi.C.Lee, Jaunzems, Mey & A.S.Rob. | 2010 | Phillipines |  | Villosae | Regiae |
| *Nepenthes paniculata* | Danser | 1928 | New Guinea |  |  | Nepenthes |
| *Nepenthes pantaronensis* | Gieray, Gronem., Wistuba, Marwinski, Micheler, Coritico & V.B.Amoroso | 2014 | Phillipines |  | Alatae | Regiae |
| *Nepenthes papuana* | Danser | 1928 | New Guinea |  |  | Urceolatae |
| *Nepenthes parvula* | G.W. Wilson & S.Ventner | 2016 | Australia |  |  |  |
| *Nepenthes pectinata* | Danser | 1928 | Sumatra |  | Montanae |  |
| *Nepenthes peltata* | Sh.Kurata | 2008 | Phillipines |  | Villosae | Regiae |
| *Nepenthes pervillei* | Blume | 1852 | Seychelles | Vulgatae |  | Nepenthes |
| *Nepenthes petiolata* | Danser | 1928 | Phillipines | Insignes | Alatae | Villosae |
| *Nepenthes philippinensis* | Macfarl. | 1908 | Phillipines | Vulgatae | Nobiles | Regiae |
| *Nepenthes pilosa* | Danser | 1928 | Borneo | Regiae | Regiae | Regiae |
| *Nepenthes pitopangii* | Chi.C.Lee, S.McPherson, Bourke & M.Mansur | 2009 | Wallacea |  | Tentaculatae | Tentaculatae |
| *Nepenthes platychila* | Chi.C.Lee | 2002 | Borneo |  | Regiae | Regiae |
| *Nepenthes pulchra* | Gronem., S.McPherson, Coritico, Micheler, Marwinski & V.B.Amoroso | 2011 | Phillipines |  | Alatae | Villosae |
| *Nepenthes rafflesiana* | Jack | 1835 | Borneo | Insignes |  | Pyrophytae |
| *Nepenthes rajah* | Hook.f. | 1859 | Borneo | Regiae | Villosae | Villosae |
| *Nepenthes ramispina* | Ridl. | 1909 | Peninsular Malaysia |  | Montanae | Pyrophytae |
| *Nepenthes ramos* | Jebb & Cheek | 2013 | Phillipines |  | Alatae | Regiae |
| *Nepenthes reinwardtiana* | Miq. | 1852 | Borneo, Sumatra | Vulgatae |  | Pyrophytae |
| *Nepenthes rhombicaulis* | Sh.Kurata | 1973 | Sumatra |  | Montanae | Montanae |
| *Nepenthes rigidifolia* | Akhriadi, Hernawati & Tamin | 2004 | Sumatra |  | Montanae | Montanae |
| *Nepenthes robcantleyi* | Cheek | 2011 | Phillipines |  | Alatae | Villosae |
| *Nepenthes rosea* | M.Catal. & Kruetr. | 2014 | Indochina |  | Pyrophatae | Pyrophytae |
| *Nepenthes rowaniae* | F.M.Bailey | 1897 | Australia |  |  |  |
| *Nepenthes samar* | Jebb & Cheek | 2013 | Phillipines |  | Insignes | Insignes |
| *Nepenthes sanguinea* | Lindl. | 1849 | Indochina | Montanae | Montanae | Pyrophytae |
| *Nepenthes saranganiensis* | Sh.Kurata | 2003 | Phillipines |  | Alatae | Regiae |
| *Nepenthes sibuyanensis* | Nerz | 1998 | Phillipines |  | Insignes | Insignes |
| *Nepenthes singalana* | Becc. | 1886 | Sumatra | Montanae | Montanae | Montanae |
| *Nepenthes smilesii* | Hemsl. | 1895 | Indochina |  | Pyrophatae | Pyrophytae |
| *Nepenthes spathulata* | Danser | 1935 | Sumatra, Java |  | Montanae | Montanae |
| *Nepenthes spectabilis* | Danser | 1928 | Sumatra | Nobiles | Montanae | Montanae |
| *Nepenthes stenophylla* | Mast. | 1890 | Borneo | Regiae | Regiae | Regiae |
| *Nepenthes sumagaya* | Cheek | 2014 | Phillipines |  | Alatae | Villosae |
| *Nepenthes sumatrana* | (Miq.) Beck | 1895 | Sumatra |  | Montanae | Montanae |
| *Nepenthes suratensis* | M.Catal. | 2010 | Indochina |  | Pyrophatae | Pyrophytae |
| *Nepenthes surigaoensis* | Elmer | 1915 | Phillipines |  | Insignes | Insignes |
| *Nepenthes talaandig* | Gronem., Coritico, Wistuba, Micheler, Marwinski, Gieray & V.B.Amoroso | 2014 | Phillipines |  | Alatae | Villosae |
| *Nepenthes talangensis* | Nerz & Wistuba | 1994 | Sumatra |  | Montanae | Montanae |
| *Nepenthes tboli* | Jebb & Cheek | 2014 | Phillipines |  | Alatae | Regiae |
| *Nepenthes tenax* | C.Clarke & R.Kruger | 2006 | Australia |  |  |  |
| *Nepenthes tentaculata* | Hook.f. | 1873 | Borneo, Wallacea | Vulgatae | Tentaculatae | Tentaculatae |
| *Nepenthes tenuis* | Nerz & Wistuba | 1994 | Sumatra |  | Montanae | Montanae |
| *Nepenthes thai* | Cheek | 2009 | Indochina |  | Montanae | Pyrophytae |
| *Nepenthes thorelii* | Lecomte | 1909 | Indochina | Vulgatae | Pyrophatae | Pyrophytae |
| *Nepenthes tobaica* | Danser | 1928 | Sumatra | Vulgatae | Montanae | Montanae |
| *Nepenthes tomoriana* | Danser | 1928 | Wallacea | Vulgatae |  | Nepenthes |
| *Nepenthes treubiana* | Warb. | 1891 | New Guinea | Regiae? Insignes |  | Nepenthes |
| *Nepenthes truncata* | Macfarl. | 1911 | Phillipines | Regiae | Alatae | Villosae |
| *Nepenthes ultra* | Jebb & Cheek | 2013 | Phillipines |  | Alatae | Regiae |
| *Nepenthes undulatifolia* | Nerz, Wistuba, U.Zimm., Chi.C.Lee, Pirade & Pitopang | 2011 | Wallacea |  | Tentaculatae | Tentaculatae |
| *Nepenthes veitchii* | Hook.f. | 1859 | Borneo | Regiae | Regiae | Regiae |
| *Nepenthes ventricosa* | Blanco | 1837 | Phillipines | Insignes | Insignes | Insignes |
| *Nepenthes vieillardii* | Hook.f. | 1873 | New Caledonia | Vulgatae |  | Nepenthes |
| *Nepenthes villosa* | Hook.f. | 1852 | Borneo | Insignes | Villosae | Villosae |
| *Nepenthes viridis* | Micheler, Gronem., Wistuba, Marwinski, W.Suarez & V.B.Amoroso | 2013 | Phillipines |  |  | Regiae |
| *Nepenthes vogelii* | Schuit. & de Vogel | 2002 | Borneo |  | Regiae | Regiae |
| *Nepenthes weda* | Cheek | 2015 | Wallacea |  | Danseri group | Nepenthes |
| *Nepenthes xiphioides* | B. Salman & Maulder | 1995 | Sumatra |  |  |  |
| *Nepenthes zakriana* | Adam & Wilcock | 2006 | Borneo |  |  |  |
| *Nepenthes zygon* | Jebb & Cheek | 2014 | Phillipines |  | Alatae | Villosae |

Appendx 1. Extant species with authorities, biogeographical regions and sectional grouping by three authorities.
