## Appendix 2 for "A phylogenomic analysis of *Nepenthes* (Nepenthaceae)"

| **Sample name** | **Tissue source** | **Accession** | **Source of accession** | **Original wild origin** | **Herbarium voucher** |
| --- | --- | --- | --- | --- | --- |
| *N. adnata* | Chester Zoo | 2012.0070 | Andreas Wistuba | Kelog Sembilan, Sumatra | Murphy 296 (K) |
| *N. adrianii* | Chester Zoo | 2012.0072 | Andreas Wistuba | Gunung Salak | Cheek 17867 (K) |
| *N. alata Luzon* | Chester Zoo | 2012.0076 | Exotica Plants | Bontoc province, Luzon, Philippines | Murphy 306 (K) |
| *N. alba* | M. Golos |  | Kamil Pasek | Gunung Tahan, Peninsular Malaysia |  |
| *N. albomarginata Brunei* | Kew living Collection | *2005-1760* | Borneo Exotics | Labi Road, Brunei, Borneo | Murphy 303 (K) |
| *N. albomarginata Pen. Malaysia* | Chester Zoo | 2012.083 |  | Gunung Jerai, N.W. Pen. Malaysia |  |
| *N. ampularia Enaratoli-Nabiri* | Chester Zoo | 2012.0112 |  | Enaratoli-Nabiri, Papua, Indonesia |  |
| *N. ampullaria Brunei green* | Kew living Collection | *2010-1266* | Borneo Exotics | Labi Road, Brunei |  |
| *N. ampullaria Brunei red* | Kew living Collection | *2010-1257* | Borneo Exotics | Labi Road, Brunei |  |
| *N. ampullaria Tayeve Giant* | Chester Zoo | 2012.0089 | Andreas Wistuba | Taiyeve, Mamberamo Regency, Papua |  |
| *N. andamana* | Chester Zoo | 2012.0123 | Marcello Cattelano | Takuapa, Phang-nga, Thailand | Murphy 365 (K) |
| *N. angasanensis* | Chester Zoo | 2012.01250 | Christian Klein | Sumatra |  |
| *N. angustifolia* | Kew living Collection | 2003-2746 | VOAR | Borneo |  |
| *N. argentii* | Andreas Wistuba |  |  | Mount Guiting-Guiting, Sibuyan |  |
| *N. aristolochoides* | Chester Zoo | 2012.01280 |  | Jambi, Sumatra |  |
| *N. armin* | Kew living Collection | 2009-830 | Borneo Exotics | Sibuyan, Philippines | Murphy 300 (K) |
| *N. attenboroughi* | M. Golos |  | Andy Smith | Palawan, Philippines |  |
| *N. barcelonae* | Andreas Wistuba |  |  | Luzon, Philippines |  |
| *N. becarriana* | Chester Zoo | 2012.0130 | Francois Mey | Sibolga, North Sumatra | Cheek 17894 (K) |
| *N. bellii* | Chester Zoo | 2012.0133 |  | Mindanao | Cheek 17884 (K) |
| *N. benstonei* | Chester Zoo | NBN-02-S | Kamil Pasek | Bukit Bakar, Kelantan, Pen. Malaysia |  |
| *N. biak* | Kew living Collection | 2003-2747 | VOAR | Biak, Papua, Indonesia | Murphy 313 (K) |
| *N. bicalcarata Brunei* | Kew living Collection | 2005-1752 | Borneo Exotics | Labi Road, Brunei | Murphy 325 (K) |
| *N. bicalcarata Sipitang* | Chester Zoo | 2011.0522 | Andreas Wistuba | Sipitang, Sabah |  |
| *N. bicalcarata Sri Aman* | Chester Zoo | Sri Aman | Malesiana Tropicals | Sri Aman, Sarawak |  |
| *N. bokoriensis* | Kew living Collection | 2009-1736 | DECLA | Bokor Falls, Cambodia | Murphy 308 (K) |
| *N. bongso* | Chester Zoo | 2011.0505 | Simon Lumb | Gunung Merapi, Sumatra | Cheek 17854 (K) |
| *N. boschiana* | Kew living Collection | 2009-826 | Borneo Exotics | Gunung Besar, South Kalimantan |  |
| *N. burbidgeae* | Chester Zoo | 2012.0149 | Andy Smith | Mt. Kinabalu, Borneo | Cheek 17827 (K) |
| *N. burki* | Chester Zoo | 2012.01500 | Borneo Exotics | Mt. Halcon, Mindoro, Philippines | Cheek 17851 (K) |
| *N. campanulata* | Chester Zoo | 2012.0152 |  | Gunung mulu, Sarawak |  |
| *N. cecilae* | Chester Zoo | 2012.0156 |  | Mt Kiamo, Mindano, Philippines | Murphy 212 (K) |
| *N. chang* | Chester Zoo | MC15 MC | Marcello Catellano | Ko Chang Island | Cheek 17897 (K) |
| *N. chaniana* | Chester Zoo | 2012.0158 | Borneo Exotics | Gunung Batu Lawi, Sarawak, BE | Cheek 17869 (K) |
| *N. clipeata* | Chester Zoo | 2012.0162 | Andreas Wistuba | Gunung Kelam, West Kalimantan | Cheek 17887 (K) |
| *N. copelandii* | M. Golos | BE-3046 | Borneo Exotics | Mt Apo, Mindanao, Philippines |  |
| *N. cornuta* | Chester Zoo | 2016.0046 |  | Mindanao, Philippines |  |
| *N. danseri* | Kew living Collection | 2004-2437 | Borneo Exotics | Waigeo, Raja Amput Regency, W. Papua | Murphy 309 (K) |
| *N. deaniana* | Chester Zoo | 2012.01670 |  | Mt Pulgar/Thumb’s Peak, Palawan |  |
| *N. densiflora* | Chester Zoo | 2012.01690 |  | Aceh, Sumatra, Indonesia | Murphy 214 (K) |
| *N. diatas* | Chester Zoo | 2012.017 | Borneo Exotics | Gunung Bandalhara, Aceh, Sumatra | Cheek 17888 (K) |
| *N. distillatoria* | Chester Zoo |  | Borneo Exotics | Sri Lanka | Murphy 366 (K) |
| *N. dubia* | Chester Zoo | 2012.0173 |  | Gunung Talakamau, W. Sumatra |  |
| *N. echinostoma* | Chester Zoo | 2013.0274 | Borneo Exotics | Lambi Hills, Sarawak |  |
| *N. edwardsiana* | M. Golos |  | Jasper Knight | Mt Tambuyukon, Sabah, Borneo |  |
| *N. ephippiata* | Chester Zoo | 2012.174 | Borneo Exotics | Gunung Raya, W. Kalimantan, Indonesia | Cheek 17880 (K) |
| *N. eustachya* | Chester Zoo | 2012.0175 | Andreas Wistuba | Kelong Semblian, Sumatra |  |
| *N. eymae* | Kew living Collection | *1983-4248* | John Turnbull | Sul Teng, Sulawesi |  |
| *N. faizaliana* | Chester Zoo | 2012.177 |  | Gunung Mulu, Sarawak | Photo voucher |
| *N. flava* | Chester Zoo | 2012 0179 | Andreas Wistuba | Sumatra | Cheek 17845 (K) |
| *N. glabrata* | Chester Zoo | 2012.0509 | Borneo Exotics | Highlands, Central Sulawesi | Cheek 17832 (K) |
| *N. glandulifera* | Chester Zoo | 2012.0187 | Exotica Plants | Hose Mountains, Sarawak, Malaysia | Cheek 17862 (K) |
| *N. gracilis Brunei* | Kew living Collection | 2005-1946 | Borneo Exotics | Labi Road, Brunei | Murphy 310 (K) |
| *N. gracilis Thailand* | Chester Zoo | 2012.0194 | Marcello Catellano | Kuche, Tak Bai, Narathiwat, S. Thailand |  |
| *N. gracilliflora loc. unknown* | Kew living Collection | 2003-2745 | VOAR |  | Murphy 305 (K) |
| *N. gracilliflora Sibuyan* | Chester Zoo | 2012.0074 | Borneo Exotics | Foothills Mt Guiting Guiting, Sibuyan | Murphy 362 (K) |
| *N. gracillima* | Chester Zoo | 2012.0196 | Christian Klein | Tahan Mnts., Pahang, Pen. Malaysia |  |
| *N. gymnamphora Java* | Chester Zoo | 2012.0197 | PG | Wonosobo, Dieng Mts., C. Java | Cheek 17864 (K) |
| *N. gymnamphora Sumatra* | Chester Zoo | 2012.0196 | Borneo Exotics | Mt Talakmau, Ophir District, Sumatra | Cheek 17834 (K) |
| *N. hamata Gunung Katoposa* | M. Golos |  | Andreas Wistuba | Gunung Katopasa, Sulawesi |  |
| *N. hamata Gunung Lumut* | M. Golos |  | Andreas Wistuba | Gunung Lumut, Sulawesi |  |
| *N. hamiguitanensis* | Andreas Wistuba |  |  | Mt Hamiguitan, Mindanao |  |
| *N. hemsleyana* | Kew living Collection | 2005-1761 | Borneo Exotics | Labi Road, Brunei | Murphy 311 (K) |
| *N. hirsuta* | Kew living Collection | 2004-2461 | Borneo Exotics | Gunung Serapai, Sarawak |  |
| *N. hispida* | Kew living Collection | 1990-931 | Borneo Exotics | Brunei | Murphy 314 (K) |
| *N. inermis* | Chester Zoo | 2012.0207 | Andreas Wistuba | Gunung Gadut, Padang, W. Sumatra | Cheek 17841 (K) |
| *N. insignis Tayeve* | Andreas Wistuba |  |  | Taiyeve, Mamberamo Regency, Papua |  |
| *N. izumae* | Chester Zoo | 2012.0213 |  | Bukit barisian, West Sumatra |  |
| *N. jacquelinae* | Chester Zoo | 2012.0216 | Borneo Exotics | Gunung Gadung, W Sumatra | Cheek 17835 (K) |
| *N. jamban* | Chester Zoo | 2011.0514 | Borneo Exotics | Bukit Barisan | Cheek 17870 (K) |
| *N. justinae* | M. Golos |  | Andreas Wistuba | Mt. Hamiguitan, Mindanao, Philippines |  |
| *N. kampotiana* | Chester Zoo | 2012.0219 | Marcello Catellano | Trat Province, Thailand |  |
| *N. kerrii* | Chester Zoo | 2012.0229 |  | Satun Province, Thailand | Murphy 361 (K) |
| *N. khasiana* | Chester Zoo |  | Andreas Wistuba | India | Cheek 17896 (K) |
| *N. klossi* | Mr Yanto, Jakarta |  |  | Papua |  |
| *N. kongkandana* | Chester Zoo | 2012.231 |  | Chana, Hat Yali, Thailand | Cheek 17875 (K) |
| *N. lamii* | Andreas Wistuba |  |  | Mount Dorman, Puncak Regency, Papua |  |
| *N. lavicola* | Chester Zoo | 2012.0233 | Andreas Wistuba | Gunung Geureadong, Sumatra |  |
| *N. leonardoi* | Chester Zoo |  |  | Palawan, Philippines |  |
| *N. lingulata* | Chester Zoo | 2012.0235 |  | Bukit Barisan, Sumatra | Cheek 17865 (K) |
| *N. longifolia* | Chester Zoo | 2012.0236 |  | Kelong Semblian, Sumatra | Murphy 363 (K) |
| *N. lowii* | Chester Zoo | 2011.0531 | GH | Gunung Mulu, Sarawak, Malaysia |  |
| *N. macfarlanei* | Kew living Collection | 2004-2390 | Borneo Exotics | Cameron Highlands, Pen. Malaysia |  |
| *N. macrophylla* | M. Golos |  | Thomas Carow | Mt. Trus Madi, Sabah, Borneo |  |
| *N. macrovulgaris* | Andreas Wistuba |  |  | Gunung Silam, Sabah, Malaysia |  |
| *N. madagascariensis* | Kew living Collection | 1985-2632 | Laurence Dorr | Fort Dauphin area, Madagascar |  |
| *N. mantalingajensis* | Andreas Wistuba |  |  | Mount Mantalingajan, Palawan |  |
| *N. mapuluensis* | Chester Zoo | 2012.0247 | Andreas Wistuba | Sankulirang Range, East Kalimantan |  |
| *N. masaolensis* | Chester Zoo | *NMO-1-S* | Andreas Wistuba | Mt Ambato, Masoala Pen., Madagascar | Cheek 17895 (K) |
| *N. maxima mini 1600m* | Chester Zoo | 2012.0258 |  | Maluku, Sulawesi, 1600m | Cheek 17842 (K) |
| *N. maxima Napu 1 Sulawesi* | Chester Zoo | NMX-09-S | Borneo Exotics | Napu valley, 350m, Sulawesi |  |
| *N. maxima Pap452* | Field |  |  | Anggi, Manokwari, W. Papua | Murphy 1003 (K) |
| *N. maxima Pap453* | Field |  |  | Sopnyai, Hingk, Manokwari, W.Papua | Murphy 1015 (K) |
| *N. maxima Tinambola Sulawesi* | Chester Zoo | 2012.0250 | Borneo Exotics | Mt.Tinambola road, Sulawesi | Cheek 17891 (K) |
| *N. maxima var. elongata* | Chester Zoo | 2012.0253 |  | Central Sulawesi | Cheek 17879 (K) |
| *N. maxima wavy Sulawesi* | Chester Zoo | 2012.0254 | Borneo Exotics | Mt. Sessian, Sulawesi | Murphy 222 (K) |
| *N. merriliana* | Kew living Collection | 2004-1766 | Borneo Exotics | Foothills of Mount Legaspi, Mindanao | Murphy 312 (K) |
| *N. micramphora* | Andreas Wistuba |  |  | Mount Hamiguitan, Mindanao |  |
| *N. mikeii* | Chester Zoo | 2012.0262 | Borneo Exotics | Mt.Bandahara, Aceh, Sumatra | Cheek 17857 (K) |
| *N. mindanoensis* | Andreas Wistuba |  |  | Dinagat |  |
| *N. minima* | Kew living Collection | 1985-3522 | John Turnbull | Lake Poso area, Sulteng., Sulawesi |  |
| *N. mira* | Chester Zoo | 2012.0264 | Andreas Wistuba | Palawan, Philippines | Cheek 17824 (K) |
| *N. mirabilis East Kalimantan* | Chester Zoo | NMB-07-S | ST | South of Samarinda, East Kalimantan |  |
| *N. mirabilis Halmahera* | Herbarium (K) |  |  | Halmahera, Maluku | Merello 3295 (K) |
| *N. mirabilis Lampia Sulawesi* | Chester Zoo | NMB-04-S | Andreas Wistuba | Gunung Lampia, Sulawesi |  |
| *N. mirabilis Morowali* | Field |  |  | Morowali, Central Sulawes | Trethowan 214 (K) |
| *N. mirabilis Pap456* | Field |  |  | Gunung Botak, Manokwari, W. Papua | Murphy 1052 (K) |
| *N. mirabilis Pap457* | Field |  |  | Gunung Botak, Manokwari, W. Papua | Murphy 1054 (K) |
| *N. mirabilis Pap462* | Field |  |  | Gunung Botak, Manokwari, W. Papua | Murphy 1043 (K) |
| *N. mirabilis Pap464* | Field |  |  | Gunung Botak, Manokwari, W. Papua | Murhpy 1048 (K) |
| *N. mirabilis Pap465* | Field |  |  | Gunung Botak, Manokwari, W. Papua | Murphy 1045 (K) |
| *N. mirabilis Pap466* | Field |  |  | Gunung Botak, Manokwari, W. Papua | Murphy 1051 (K) |
| *N. mirabilis Pap467* | Field |  |  | Gunung Botak, Manokwari, W. Papua | Murphy 1044 (K) |
| *N. mirabilis Pap468* | Field |  |  | Gunung Botak, Manokwari, W. Papua | Murphy 1053 (K) |
| *N. mirabilis Pap471* | Field |  |  | Gunung Botak, Manokwari, W. Papua | Murphy 1055 (K) |
| *N. mirabilis Pap472* | Field |  |  | Gunung Botak, Manokwari, W. Papua | Murhpy 1050 (K) |
| *N. mirabilis Pap475a* | Field |  |  | Gunung Botak, Manokwari, W. Papua | Murphy 1059 (K) |
| *N. mirabilis Pap475b* | Field |  |  | Gunung Botak, Manokwari, W. Papua | Murphy 1059 (K) |
| *N. mirabilis Sabah* | Kew living Collection | 1981-5655 | John Lonsdale | Nr Kimarnis, Beaufort Road, Sabah | Murphy 324 (K) |
| *N. mirabilis Thailand* | Chester Zoo | 2012.0269 |  | Trat, Thailand |  |
| *N. mirabilis var. globosa Trang* | Chester Zoo | 2012.0277 | Andreas Wistuba | Trang, Thailand | Murphy 364 (K) |
| *N. mirabilis Wawoni* | Field |  |  | Wawonii, South East Sulawesi | Trethowan 527 (K) |
| *N. mirabilis West Kalimantan* | Kew living Collection | 1983-2176 | John Turnbull | Pasir Panjang, W. Kalimantan | Murphy 304 (K) |
| *N. mollis* | Chester Zoo | 2012.0206 | Christian Klein | Borneo |  |
| *N. monticola* | Herbarium (K) |  |  | Freeport, Mimika Regency, Papua | Willis 147 (K) |
| *N. muluensis* | M. Golos |  | Christian Klein | Gunung Murud, Sarawak, Borneo |  |
| *N. murudensis* | Chester Zoo | 2016.0094 | Borneo Exotics | Gunung Murud, Sarawak, Borneo |  |
| *N. naga* | Chester Zoo | 2012.0281 | Gareth Davies | Bukit Bansan, Sumatra | Cheek 17860 (K) |
| *N. neoguinensis Freeport* | Herbarium (K) |  |  | Freeport, Mimika Regency, Papua | Argent 521 (K) |
| *N. neoguinensis Jayapura* | Chester Zoo | 2012.0283 |  | Jayapura, Papua |  |
| *N. nigra* | Andreas Wistuba |  |  | Gunung Katopasa, Sulawesi |  |
| *N. northiana* | Chester Zoo | 2012.0284 | Borneo Exotics | Near Bau, S. Kuching, Sarawak |  |
| *N. oblanceolata* | Chester Zoo | 2016.0049 |  | Wamena area, Papua, Indonesia |  |
| *N. orbiculata* | Chester Zoo | NGS-03-1S | Christian Klein | Takuapa, Phang-nga, Thailand | Cheek 17877 (K) |
| *N. ovata* | Chester Zoo | 2012.0286 | Andy Smith | Gunung Pangulabao, Sumatra | Cheek 17830 (K) |
| *N. palawanensis* | M. Golos |  | Simon Lumb | Sultan Peak, Palawan, Philippines |  |
| *N. pantaronensis* | Andreas Wistuba |  |  | Mindanao, Philippines |  |
| *N. papuana* | Chester Zoo | 2016.0052 | Andreas Wistuba | Mt. Doorman, Puncak Regency, Papua |  |
| *N. peltata* | Chester Zoo | 2012.0289 |  | Mount Hamiguitan, Mindanao | Cheek 17849 (K) |
| *N. pervillei* | Andreas Wistuba |  |  | Seychelles |  |
| *N. petiolata* | Kew living Collection | 2004-2180 | Borneo Exotics | Mount Hilong-Hilong, Mindanao | Murphy 320 (K) |
| *N. phillippinensis* | Chester Zoo | 2012.0293 | Borneo Exotics | Palawan | Cheek 17872 (K) |
| *N. pitopangi* | Chester Zoo | 2016.0009 | Andreas Wistuba | Wistuba |  |
| *N. platychylla* | Chester Zoo | 2012.0296 | Borneo Exotics | Bukit Batu, Hose Mountains, Sarawak | Cheek 17873 (K) |
| *N. pulchra* | Andreas Wistuba |  |  | Mindanao, Philippines |  |
| *N. rafflesiana Kinabalu* | Kew living Collection | 1998-1487 | PELE | Mt Kinabalu, Sabah |  |
| *N. rafflesiana Kuching* | Chester Zoo | 2012.0306 | Malesiana Tropicals | Kuching, Sarawak |  |
| *N. rafflesiana Pen. Malaysia* | Chester Zoo | 2012.0301 | Borneo Exotics | Jahore Bahru, Peninsular Malaysia |  |
| *N. rafflesiana var. Nivea Sarawak* | Chester Zoo | 2012.0297 | Borneo Exotics | Lundu, Sarawak, Malaysia |  |
| *N. rajah* | Kew living Collection | 1982-144 | Tony Lamb | Gunung Tamboyokan, Kinabalu, Sabah |  |
| *N. ramispina* | Chester Zoo | 2012.0313 | Borneo Exotics | Gunung Ulu Kali, Pen. Malaysia | Cheek 17859 (K) |
| *N. reinwardtiana Crocker Range* | Kew living Collection | 2005-1947 | Borneo Exotics | Tambunan Road, Crocker Range, Sabah | Murphy 319 (K) |
| *N. rhombicaulis* | Chester Zoo | 2012.0320 | Simon Lumb | Gunung Pangulabao, Sumatra | Cheek 17831 (K) |
| *N. rigidifolia* | Chester Zoo | 2016.005 | Simon Lumb | Mt. Sidikalang, Karo Regncy, Sumatra |  |
| *N. robcantleyi* | Kew living Collection | 2012-155 | Borneo Exotics | Mindanao |  |
| *N. rowanae* | Chester Zoo | *NRO-01-S* | Exotica Plants | Cape York Pen., Queensland, Australia | Cheek 17886 (K) |
| *N. sanguinea* | Kew living Collection | 2004-2415 | Borneo Exotics | Cameron Highlands, Pen. Malaysia | Murphy 317 (K) |
| *N. sibuyanensis* | Kew living Collection | 2004-2403 | Borneo Exotics | Mount Guiting-Guiting, Sibuyan Island | Murphy 316 (K) |
| *N. singalana* | Kew living Collection | 2010-1263 | Borneo Exotics | Gunung Belirang, Sumatra |  |
| *N. smilesii Kirirom NP Cambodia* | Chester Zoo | NSM-04-S | Francois Mey | Kiriom, Cambodia |  |
| *N. smilesii Phu Kradeung NP* | Chester Zoo | 2012.0329 |  | Phu Kradung, Loei, Thailand |  |
| *N. sp. Anipahan* | M. Golos |  | Christian Klein | Mt. Anipahan, Palawan, Philippines |  |
| *N. sp. tamini* | M. Golos |  | Andy Smith | Sumatra | Murphy 1089 (K) |
| *N. spathulata* | Chester Zoo | 2011.0499 | Borneo Exotics | Sumatra | Cheek 17829 (K) |
| *N. spectabilis Gunung Bandalhara* | Chester Zoo | 2012.0340 | Borneo Exotics | Gunung Bandahara, Sumatra | Cheek 17881 (K) |
| *N. spectabilis Pangalubao* | Chester Zoo | 2011.0523 | Borneo Exotics | Gunung Pangulabao, N. Sumatra | Cheek 17848 (K) |
| *N. stenophylla* | Chester Zoo | 2012.0344 | Borneo Exotics | Bareo, Sarawak |  |
| *N. sumagaya* | Chester Zoo | 2016.0015 | Andreas Wistuba | Mount Sumagaya, Mindanao |  |
| *N. suratensis* | Chester Zoo | NSS-01-S | GH | Suratthani Province, Thailand | Cheek 17899 (K) |
| *N. surigaoensis* | Chester Zoo | 2017.0772 | Andreas Wistuba | Mt Masay Elliot, Mindanao, Philippines |  |
| *N. talaandig* | Andreas Wistuba |  |  | Mindanao, Philippines |  |
| *N. talangensis* | Chester Zoo | 2012.0350 | Borneo Exotics | Gunung Talang, Sumatra | Cheek 17861 (K) |
| *N. tenax* | Chester Zoo | 2012.0351 |  | Jardine Swamp, Cape York, Australia | Cheek 17866 (K) |
| *N. tentaculata Gunung Murud* | Chester Zoo | 2012.0353 | Borneo Exotics | Gunung Murud, Sarawa, Malaysia | Cheek 17844 (K) |
| *N. tentaculata Gunung Raja* | Chester Zoo | 2012.0357 | Andreas Wistuba | Gunung Raja, South Kalimantan |  |
| *N. tenuis* | Chester Zoo | 2012.0359 | Simon Lumb | Mt.Taram,Tjampo River, W.Sumatra | Cheek 17863 (K) |
| *N. thai* | Chester Zoo | 2012.0360 |  | Kao Aidang, Hala Bala N.P, Thailand | Cheek 17874 (K) |
| *N. thorellii Bin Chau* | Chester Zoo | 2012.0361 | Gareth Davies | Bin Chau Vietnam “Mr Son” |  |
| *N. tobaica Lake Toba* | Kew living Collection | 2004-2432 | Borneo Exotics | Lake Toba, Sumatra | Murphy 321 (K) |
| *N. tobaica Red Doloksanggul* | Chester Zoo | 2012.0364 | Borneo Exotics | Doloksangul | Cheek 17833 (K) |
| *N. tomoriana* | Chester Zoo | 2012.0365 | Borneo Exotics | Gunung Lampia, Tomoroi bay, Sulawesi |  |
| *N. treubiana FakFak* | Chester Zoo | 2012.0366 |  | Fak Fak, Papua | Cheek 17871 (K) |
| *N. treubiana Misool* | Chester Zoo | 2012.0366 |  | Raja Amput Regency, W. Papua |  |
| *N. truncata* | Kew living Collection | 2004-2404 | Borneo Exotics | Gunung Pasian, Mindanao |  |
| *N. undulatifolia* | Andreas Wistuba |  |  | Sulawesi |  |
| *N. veitchii* | Kew living Collection | 2005-1757 | Borneo Exotics | Bario Highlands, Sarawak | Murphy 323 (K) |
| *N. ventricosa* | Kew living Collection | 2004-2442 | Borneo Exotics | Luzon | Murphy 322 (K) |
| *N. vieilliardii Bauman 15242* | Herbarium (K) |  |  | New Caledonia | Baumann 15241 (K) |
| *N. vieilliardii Buckholz 1464* | Herbarium (K) |  |  | New Caledonia | Buckholz 1463 (K) |
| *N. villosa* | Andreas Wistuba |  |  | Mount Kinabalu |  |
| *N. viridis* | Andreas Wistuba |  |  | Dinagat, Philippines |  |
| *N. vogelii* | Kew living Collection | 2005-2189 | Borneo Exotics | Hose Mountains, Sarawak |  |
| *N. x hookeriana* | Chester Zoo | 2005-1938 | Borneo Exotics | Labi Road, Brunei |  |
| *N. x trichocarpa* | Kew living Collection | *2004-2458* | Borneo Exotics | Labi Road, Brunei | Murphy 301 (K) |
| *N. xiphioides* | Chester Zoo | 2012.0387 | Andreas Wistuba | Toba Region | Cheek 17868 (K) |
| *N. zakriana Gunung Alab* | Chester Zoo | 2011.0513 |  | Gunung Alab, Crocker Range, Sabah | Cheek 17892 (K) |
| *N. zygon* | Kew living Collection | 2004.2413 | Borneo Exotics | Mount Pasian, Mindanao | Murphy 302 (K) |
| *Achatocarpus nigricans* | Herbarium (K) |  |  |  | Garvizu 1104 (K) |
| *Anacampseros papyracea* | Herbarium (K) |  |  |  | Chase 10986 (K) |
| *Ancistrocladus abbreviatus* | Kew Herbarium |  |  |  | Van der Burg 2094 (K) |
| *Ancistrocladus korupensis* | Herbarium (K) |  |  |  | Gereau 5203 (K) |
| *Armeria alliacea* | Herbarium (K) |  |  |  | Chase 20107 (K) |
| *Asteropeia microster* | Herbarium (K) |  |  |  | Civeyrel, L. (K) |
| *Barbeuia madagascariensis* | Herbarium (K) |  |  |  | Zarucetti 7407 (K) |
| *Dianthus fruticosus* | D. Soltis (1KP project) |  |  |  | Soltis&Miles2849(MO) |
| *Drosera dicrosepala* | Kew living Collection | 2001-2740 |  |  |  |
| *Drosophyllum lusitanicum* | Herbarium (K) |  |  |  | Chase 33141 (K) |
| *Emex spinosa* | Herbarium (K) |  |  |  | Chase 1016 (K) |
| *Frankenia laevis* | Herbarium (K) |  |  |  | Chase 38662 (K) |
| *Gisekia africana* | Herbarium (K) |  |  |  | Germhuizen 2382 (K) |
| *Halophytum ameghinoi* | Herbarium (K) |  |  |  | Chase 1753 (K) |
| *Limeum aethhiopicum* | Herbarium (K) |  |  |  | Hugo 2376 (K) |
| *Limonium aborescens* | Herbarium (K) |  |  |  | Chase 1649 (K) |
| *Lophiocarpus burchellii* | Herbarium (K) |  |  |  | de Winter 3156 (K) |
| *Macarthuria kerighenji* | Herbarium (K) |  |  |  | Dixon 1017 (K) |
| *Microtea debilis* | Herbarium (K) |  |  |  | Chuml 301 (K) |
| *Petrorhagia velutina* | Herbarium (K) |  |  |  | Chase 8853 (K) |
| *Plumbago auriculata* | Herbarium (NYBG) |  |  |  | Forest. F.749 NYBG |
| *Polygonum campanulatum* | Kew Living Collection | 1973-1047 |  |  |  |
| *Rhabdodendron amazonicum* | Herbarium (K) |  |  |  | E. Ribeiro 1187 (K) |
| *Rumex induratus* | Herbarium (K) |  |  |  | Chase 925 (K) |
| *Spergularia heldreichii* | Herbarium (K) |  |  |  | Chase 8847 (K) |
| *Stegnosperma cubense* | Herbarium (K) |  |  |  | Herrera 7343 (K) |
| *Sueda vera* | Herbarium (K) |  |  |  | Chase 11007 (K) |
| *Tamarix chinensis* | Herbarium (K) |  |  |  | Chase 33138 (K) |
| *Triphyophyllum peltatum* | Herbarium (K) |  |  |  | Chase 663 (K) |

Appendix 2. Sample origins and voucher specimens. Accession numbers are given for samples collected from cultivated collections at RBG Kew and Chester Zoo. For these samples, source of accession and original wild origin refers to information on the original source of these collections where available. Herbarium acronyms according to Thiers (2019).
