## Supplementary figures and images for "A phylogenomic analysis of *Nepenthes* (Nepenthaceae)"

### Supplementary Figure S1

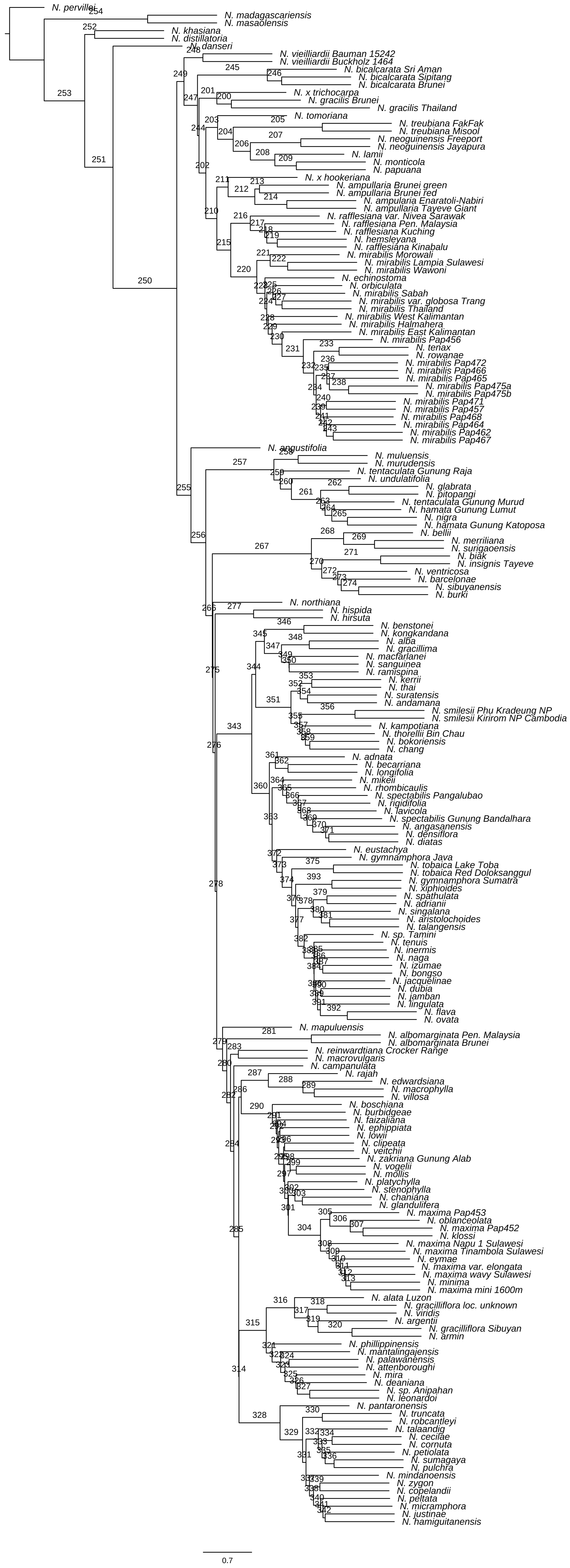

### Supplementary Figure S2

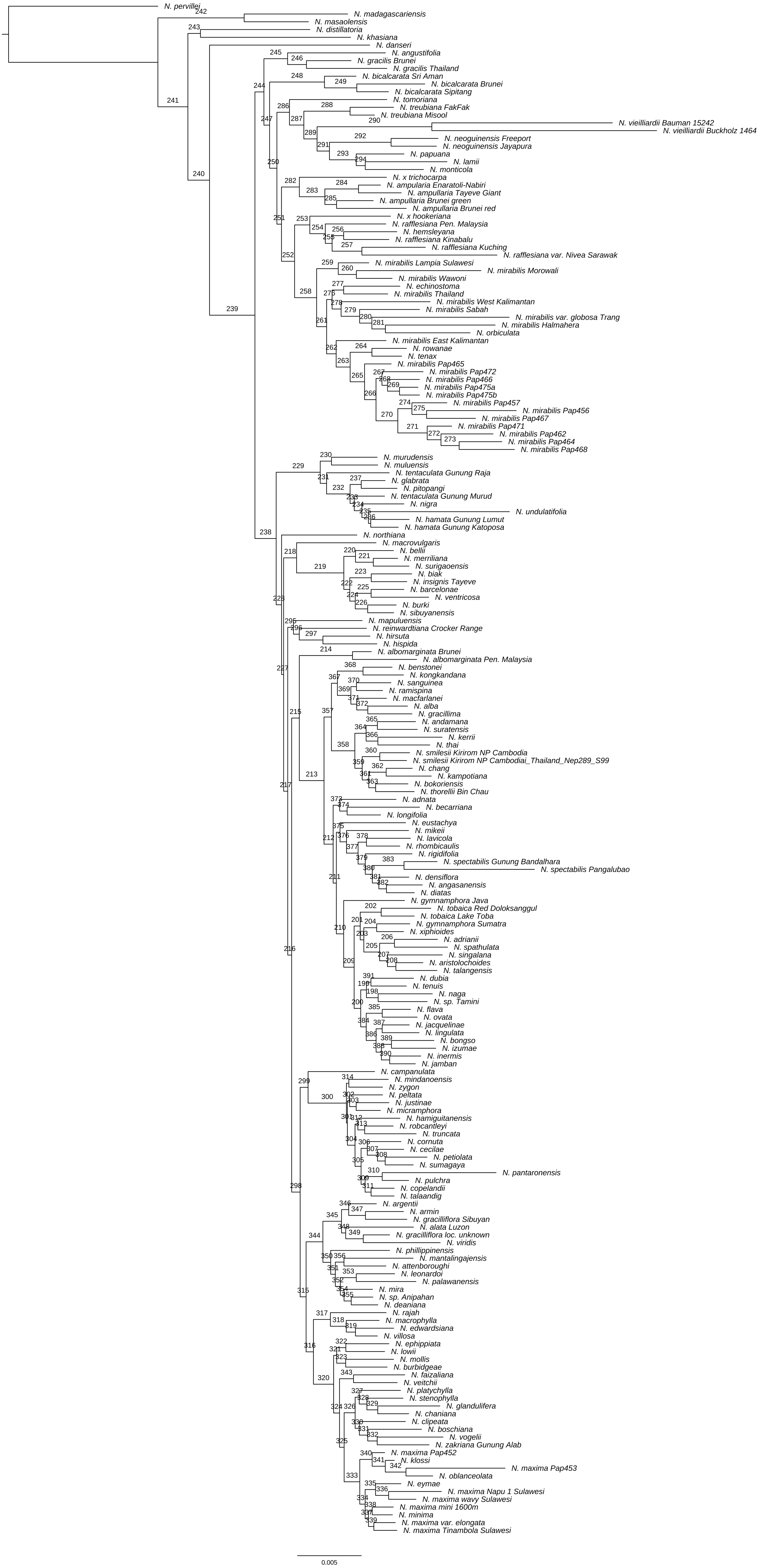

### Supplementary Figure S3

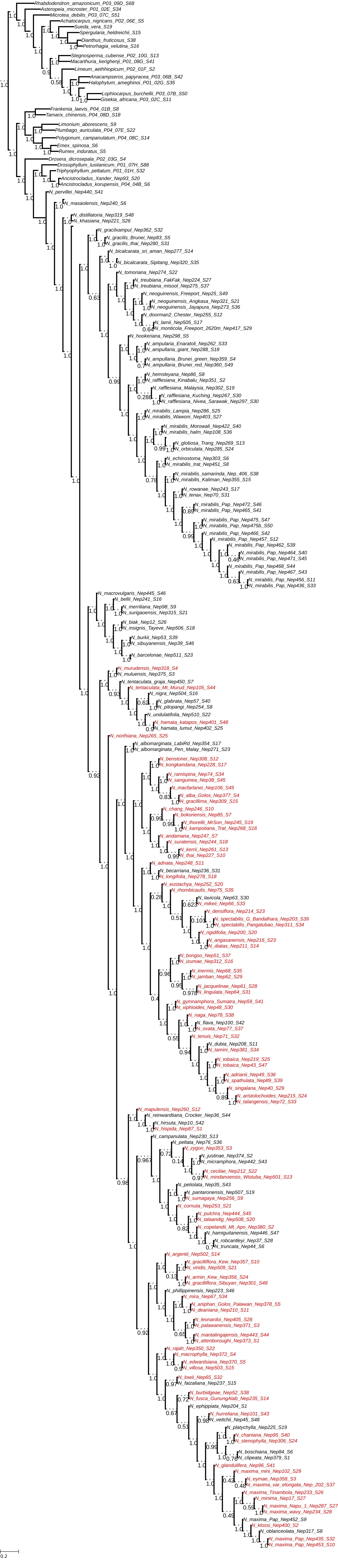
