## Supplementary Table S3 for "A phylogenomic analysis of *Nepenthes* (Nepenthaceae)"

| **ID** | **gCF** | **gDF1** | **gDF2** | **gN** | **sCF** | **sDF1** | **sDF2** | **sN** | **P.P** | **Length** |
| --- | --- | --- | --- | --- | --- | --- | --- | --- | --- | --- |
| 200 | 24.38 | 14.84 | 6.01 | 566 | 44.84 | 27.6 | 27.56 | 2550.44 | 1 | 0.210093 |
| 201 | 11.22 | 0.17 | 0 | 597 | 54.58 | 25.79 | 19.63 | 2040.69 | 1 | 0.25821 |
| 202 | 0.47 | **0.47** | 0 | 642 | 38.14 | 34.64 | 27.23 | 1362.11 | 0.98 | 0.098955 |
| 203 | 3.34 | 0.17 | 0.17 | 598 | 45.78 | 25.26 | 28.96 | 1393.18 | 1 | 0.169645 |
| 204 | 8.29 | 3.05 | 1.86 | 591 | 46.05 | 25.07 | 28.88 | 1300.38 | 1 | 0.226041 |
| 205 | 60.88 | 0.17 | 0.51 | 593 | 79.9 | 10.27 | 9.83 | 2087.66 | 1 | 1.28832 |
| 206 | 15.57 | 3.93 | 5.08 | 610 | 52.63 | 20.32 | 27.05 | 1235.24 | 1 | 0.248166 |
| 207 | 47.53 | 3.14 | 2.69 | 446 | 61.51 | 17.99 | 20.5 | 1193.27 | 1 | 0.729175 |
| 208 | 27.97 | 2.12 | 5.93 | 236 | 53.18 | 27.39 | 19.43 | 530.89 | 1 | 0.353553 |
| 209 | 29.24 | 16.95 | 12.71 | 236 | 54.47 | 26.29 | 19.24 | 486 | 1 | 0.311194 |
| 210 | 0.47 | 0.16 | 0.16 | 644 | 39.33 | 29.71 | 30.95 | 1329.26 | 1 | 0.158999 |
| 211 | 5.7 | 1.79 | 0 | 614 | 48.22 | 30.8 | 20.98 | 1924.81 | 1 | 0.163036 |
| 212 | 14.03 | 2.15 | 2.64 | 606 | 51.63 | 25.95 | 22.42 | 2330.83 | 1 | 0.385772 |
| 213 | 14.31 | 7.33 | 8.52 | 587 | 39.06 | 29.81 | 31.13 | 2097.68 | 0.94 | 0.0679045 |
| 214 | 30.99 | 4.86 | 2.35 | 597 | 63.72 | 18.9 | 17.39 | 2167.54 | 1 | 0.463237 |
| 215 | 2.33 | 0.31 | 0.16 | 644 | 41.33 | 32.11 | 26.55 | 1478.96 | 1 | 0.201137 |
| 216 | 9.28 | 0.26 | 0 | 388 | 57.12 | 24.59 | 18.28 | 1261.04 | 1 | 0.282544 |
| 217 | 7.34 | 5.43 | 4.08 | 368 | 40.69 | 24.08 | 35.23 | 969.26 | 1 | 0.208399 |
| 218 | 7.02 | **7.58** | 6.1 | 541 | 36.02 | 30.85 | 33.13 | 1496.17 | 0.63 | 0.0298473 |
| 219 | 18.38 | 10.12 | 8.26 | 593 | 39.27 | 32.77 | 27.96 | 1722.45 | 1 | 0.152354 |
| 220 | 3.62 | 0.79 | 0 | 636 | 53.94 | 22.15 | 23.9 | 1397.55 | 1 | 0.376724 |
| 221 | 11.55 | 1.44 | 0.36 | 277 | 51.41 | 26.45 | 22.14 | 722.64 | 1 | 0.193211 |
| 222 | 17.41 | 12.22 | 6.67 | 270 | 34.1 | **39.11** | 26.79 | 625.05 | 1 | 0.252891 |
| 223 | 1.6 | 0.64 | 0.32 | 624 | 39.93 | 32.89 | 27.18 | 1540.17 | 1 | 0.117778 |
| 224 | 0.48 | **0.79** | 0.16 | 629 | 34.78 | 32.35 | 32.87 | 1617.64 | 0.61 | 0.0259718 |
| 225 | 1.75 | 0.19 | 0.19 | 513 | 40.24 | 26.87 | 32.89 | 1483.12 | 0.95 | 0.0875008 |
| 226 | 0.8 | **3.19** | **1.99** | 502 | 30.65 | **38.36** | **30.99** | 1741.99 | 0.69 | 0.0425386 |
| 227 | 7.67 | 4.7 | 4.01 | 574 | 37.76 | **40.04** | 22.2 | 1755.41 | 0.99 | 0.0915804 |
| 228 | 0.67 | **0.84** | 0 | 595 | 29.06 | **34.69** | **36.25** | 1551.7 | 0.46 | 0.0208559 |
| 229 | 1.11 | **1.11** | 0.37 | 540 | 42.51 | 29.57 | 27.91 | 1172.86 | 0.51 | 0.0574655 |
| 230 | 1.44 | **1.44** | **2.34** | 556 | 38.22 | 19.38 | **42.41** | 1218.36 | 1 | 0.142402 |
| 231 | 3.24 | 0.27 | 0.81 | 370 | 57.76 | 13.12 | 29.12 | 895.62 | 1 | 0.308223 |
| 232 | 1.13 | 0.85 | 0.85 | 353 | 31.93 | 20.72 | **47.35** | 558.44 | 1 | 0.15018 |
| 233 | 12.64 | 0 | 0 | 356 | 60.82 | 19.12 | 20.06 | 583.04 | 1 | 0.367042 |
| 234 | 0.95 | 0.76 | 0 | 528 | 39.35 | 27.21 | 33.44 | 812.84 | 0.62 | 0.0352902 |
| 235 | 1.04 | **1.22** | 0 | 576 | 37.32 | 33.42 | 29.26 | 998.15 | 1 | 0.154772 |
| 236 | 3.41 | 1.25 | 0.72 | 558 | 39.6 | 24.3 | 36.11 | 1218.84 | 0.56 | 0.0249111 |
| 237 | 2.82 | 1.33 | 1.33 | 602 | 31.37 | **35.24** | **33.39** | 1390 | 0.69 | 0.0371803 |
| 238 | 8.46 | 2.54 | 2.37 | 591 | 47.88 | 19.62 | 32.5 | 1376.27 | 1 | 0.277215 |
| 239 | 1.18 | 0 | 0 | 509 | 48.06 | 26.27 | 25.67 | 851.6 | 0.87 | 0.0634646 |
| 240 | 2.2 | 1.26 | 0 | 318 | 25.6 | **38.26** | **36.14** | 545.08 | 0.85 | 0.0847509 |
| 241 | 1.99 | **2.43** | 0.88 | 453 | 37.05 | 31.91 | 31.04 | 758.76 | 0.79 | 0.0522696 |
| 242 | 2.15 | **2.63** | 1.91 | 418 | 34.9 | 28.74 | 36.36 | 903.47 | 0.59 | 0.0328078 |
| 243 | 3.47 | **4.05** | 2.6 | 346 | 37.63 | 29.52 | 32.85 | 737.65 | 0.96 | 0.0990882 |
| 244 | 0.16 | **0.16** | **0.62** | 643 | 34.95 | **35.02** | 30.02 | 1474.62 | 0.47 | 0.021944 |
| 245 | 48.91 | 0 | 0 | 595 | 75.86 | 14.42 | 9.72 | 2316.9 | 1 | 1.00965 |
| 246 | 32.72 | 21.44 | 15.16 | 541 | 53.85 | 26.34 | 19.81 | 1475.25 | 1 | 0.20737 |
| 247 | 0.44 | **0.44** | 0 | 454 | 28.7 | 25.77 | 45.53 | 269.79 | 1 | 0.195293 |
| 248 | 17.39 | 0 | 0.43 | 230 | 87.94 | 6.74 | 5.31 | 225.73 | 1 | 0.273616 |
| 249 | 1.1 | 0.44 | 0.44 | 454 | 47.76 | 28.01 | 24.23 | 313.79 | 0.8 | 0.104145 |
| 250 | 37.79 | 0 | 0 | 614 | 68.12 | 15.18 | 16.7 | 2920.87 | 1 | 0.92202 |
| 251 | 31.43 | 6.51 | 8.14 | 614 | 51.63 | 21.3 | 27.07 | 2873.12 | 1 | 0.409664 |
| 252 | 28.21 | 8.87 | 14.58 | 631 | 39.31 | 26.71 | 33.98 | 2971.56 | 1 | 0.144237 |
| 253 | 43.99 | 3.32 | 6.33 | 632 | 55.7 | 21 | 23.3 | 3482.17 | 1 | 0.585933 |
| 254 | 79.64 | 4.07 | 3.58 | 614 | 78.85 | 13.42 | 7.73 | 4742.65 | 1 | 1.49612 |
| 255 | 1.83 | 0 | 1.22 | 655 | 43.01 | 21.05 | 35.94 | 2203.65 | 1 | 0.205473 |
| 256 | 1.83 | 0.15 | 0.46 | 655 | 49.26 | 21.91 | 28.83 | 2031.38 | 1 | 0.213894 |
| 257 | 24.73 | 0 | 0 | 651 | 71.22 | 16.12 | 12.66 | 2231.13 | 1 | 0.98497 |
| 258 | 24.72 | 4.25 | 6.3 | 635 | 47.65 | 25.66 | 26.7 | 1833.6 | 1 | 0.350657 |
| 259 | 11.61 | **12.72** | 4.45 | 629 | 36.1 | **38.83** | 25.07 | 1707.26 | 0.82 | 0.0938017 |
| 260 | 15.21 | 2.59 | 5.18 | 309 | 48.51 | 15.78 | 35.71 | 711.59 | 0.65 | 0.159738 |
| 261 | 13.27 | 2.91 | 2.27 | 309 | 54.22 | 18.97 | 26.81 | 591.1 | 1 | 0.421899 |
| 262 | 25.97 | 4.55 | 2.92 | 616 | 55.93 | 18.06 | 26.01 | 1317.23 | 1 | 0.409285 |
| 263 | 6.63 | 5.66 | 5.5 | 618 | 38.89 | 30.63 | 30.48 | 1255.64 | 0.91 | 0.0640406 |
| 264 | 8.26 | **8.93** | 3.97 | 605 | 42.84 | 34.54 | 22.62 | 1084.73 | 0.99 | 0.0978623 |
| 265 | 21.88 | 7.52 | 13.16 | 585 | 42.87 | 18.82 | 38.31 | 1247.48 | 1 | 0.226616 |
| 266 | 0.76 | **0.76** | 0 | 656 | 36.58 | 30.71 | 32.7 | 1757.67 | 0.99 | 0.0945915 |
| 267 | 36.91 | 0 | 0 | 634 | 83.38 | 8.63 | 7.99 | 3066.06 | 1 | 1.43313 |
| 268 | 27.2 | 1.99 | 4.81 | 603 | 57.58 | 22.42 | 20 | 1396.97 | 1 | 0.464839 |
| 269 | 37.18 | 10.87 | 7.81 | 589 | 52.36 | 29.49 | 18.15 | 1367.52 | 1 | 0.447532 |
| 270 | 12.44 | 3.55 | 5.82 | 619 | 37.84 | 34.06 | 28.1 | 1183.22 | 1 | 0.148779 |
| 271 | 46.78 | 0.99 | 1.32 | 605 | 63.02 | 14.89 | 22.09 | 1654.88 | 1 | 0.851853 |
| 272 | 13.52 | 5.03 | 2.95 | 577 | 42.17 | 28.91 | 28.91 | 1165.5 | 1 | 0.231587 |
| 273 | 9.01 | **14.66** | **9.89** | 566 | 42.3 | 35.99 | 21.71 | 1174.81 | 0.81 | 0.0503648 |
| 274 | 24.06 | 6.64 | 9.16 | 557 | 41.53 | 30.74 | 27.73 | 1496.82 | 1 | 0.252336 |
| 275 | 0 | **2.05** | **0.32** | 633 | 31.42 | **35** | **33.58** | 1826.12 | 0.41 | 0.00838823 |
| 276 | 0 | **1.94** | **0** | 617 | 37.82 | 34.52 | 27.65 | 1830.11 | 0.54 | 0.0326788 |
| 277 | 20.93 | 0 | 0 | 602 | 63.33 | 18.91 | 17.76 | 2150.9 | 1 | 0.555614 |
| 278 | 0 | **0** | **0.16** | 632 | 33.78 | 32.71 | 33.51 | 1807.73 | 0.56 | 0.0224745 |
| 279 | 0 | **0.16** | **0** | 642 | 40.35 | 27.77 | 31.87 | 1688.9 | 0.97 | 0.0845703 |
| 280 | 0 | **1.12** | **0** | 624 | 38.35 | 25.14 | 36.51 | 1989.46 | 0.77 | 0.0606467 |
| 281 | 49.35 | 0 | 0 | 616 | 76.94 | 10.83 | 12.23 | 3043.43 | 1 | 1.22323 |
| 282 | **0** | **0.16** | **0.16** | 623 | 37.12 | 26.55 | 36.33 | 1962.28 | 0.78 | 0.0545163 |
| 283 | 5.54 | 0 | 0 | 596 | 39 | 27.52 | 33.48 | 2039.11 | 1 | 0.112785 |
| 284 | 0.32 | **0.64** | 0 | 622 | 38.06 | 29.51 | 32.43 | 2047.75 | 0.76 | 0.0475168 |
| 285 | 0.32 | 0.16 | 0 | 634 | 36.45 | 34.37 | 29.18 | 1798.46 | 0.95 | 0.0709241 |
| 286 | 0.92 | 0 | 0 | 652 | 38.55 | 30.33 | 31.12 | 1931.04 | 0.7 | 0.0360545 |
| 287 | 12.12 | 0.48 | 0.16 | 619 | 55.99 | 22.08 | 21.93 | 1865.09 | 1 | 0.392596 |
| 288 | 24.96 | 5.79 | 2.98 | 605 | 59.3 | 21.13 | 19.57 | 1736.18 | 1 | 0.50013 |
| 289 | 24.67 | 9 | 18.5 | 600 | 43.7 | 24.42 | 31.88 | 1449.63 | 1 | 0.163895 |
| 290 | 2.68 | 0 | 0 | 635 | 58.86 | 18.73 | 22.41 | 2186.32 | 1 | 0.449963 |
| 291 | 0.97 | **0.97** | 0.32 | 621 | 36.84 | 27.81 | 35.35 | 1779.01 | 0.76 | 0.0560695 |
| 292 | 0.16 | **2.59** | **0.32** | 618 | 30.68 | **34.58** | **34.74** | 1810.96 | 0.47 | 0.0153214 |
| 293 | 0 | **1.12** | **0** | 625 | 38.91 | 32.13 | 28.96 | 1649.06 | 0.53 | 0.0211324 |
| 294 | 5.46 | 0 | 0.17 | 604 | 35.7 | 33.49 | 30.82 | 1542.37 | 0.49 | 0.0170535 |
| 295 | 0.48 | 0 | 0 | 628 | 34.64 | 31.75 | 33.61 | 1605.88 | 0.94 | 0.070037 |
| 296 | 2.48 | 0 | 0 | 604 | 39.29 | 24.92 | 35.79 | 1736.4 | 0.42 | 0.0149844 |
| 297 | 0 | **0** | **0.16** | 625 | 36.6 | 29.95 | 33.45 | 1669.6 | 0.49 | 0.0160209 |
| 298 | 1.61 | 0 | 0.16 | 623 | 37.41 | 28.76 | 33.84 | 1448.24 | 0.97 | 0.0794949 |
| 299 | 6.5 | 3 | 4.17 | 600 | 34.56 | 29.81 | **35.63** | 1485.4 | 0.98 | 0.0885131 |
| 300 | 0.16 | **0.47** | **0.16** | 633 | 33.85 | **36.17** | 29.99 | 1526.55 | 0.64 | 0.0466727 |
| 301 | 0.16 | **0.47** | **0.93** | 642 | 38.93 | 31.09 | 29.98 | 1504.53 | 0.4 | 0.00697171 |
| 302 | 2.23 | 0.79 | 0.64 | 629 | 34.21 | 33.28 | 32.52 | 1541.85 | 0.99 | 0.0944078 |
| 303 | 8.29 | 5.01 | 3.8 | 579 | 42.99 | 33.43 | 23.58 | 1619.02 | 0.99 | 0.108126 |
| 304 | 8.41 | 0.16 | 0 | 642 | 59.85 | 17.52 | 22.62 | 1455.32 | 1 | 0.464397 |
| 305 | 9.78 | 0 | 0.82 | 368 | 49.12 | 19.17 | 31.71 | 669.65 | 0.5 | 0.134168 |
| 306 | 8.03 | **8.31** | 2.77 | 361 | 36.28 | **41.91** | 21.8 | 458.9 | 1 | 0.29574 |
| 307 | 9.52 | 5.75 | 9.87 | 557 | 41.95 | 20.89 | 37.17 | 758.39 | 1 | 0.185507 |
| 308 | 2.18 | 0.79 | 1.39 | 505 | 46.56 | 29.04 | 24.4 | 974.73 | 1 | 0.125032 |
| 309 | 0.6 | **2** | **2.99** | 501 | 36.09 | 25.11 | **38.8** | 950.49 | 0.94 | 0.100805 |
| 310 | 0.98 | **2.77** | 0.49 | 613 | 36.3 | 27.03 | **36.67** | 1088.07 | 0.88 | 0.0635979 |
| 311 | 0.33 | **1.8** | 0.16 | 611 | 41.27 | 22.31 | 36.43 | 1021.56 | 0.79 | 0.0584892 |
| 312 | 0.51 | **4.23** | 1.52 | 591 | 37.89 | 31.99 | 30.12 | 946.44 | 0.44 | 0.0185475 |
| 313 | 6.2 | 2.85 | 4.69 | 597 | 31.14 | **35.78** | **33.08** | 944.76 | 0.9 | 0.0697446 |
| 314 | 0.15 | **0.3** | **0.15** | 661 | 34.33 | 32.91 | 32.76 | 1824.74 | 0.38 | 0.00440233 |
| 315 | 2.9 | 0 | 0 | 655 | 53.93 | 24.08 | 21.99 | 1934.92 | 1 | 0.393993 |
| 316 | 14.33 | 0 | 1.42 | 635 | 57.82 | 18.17 | 24.02 | 1840.3 | 1 | 0.408839 |
| 317 | 8.32 | 5.12 | 3.36 | 625 | 42.74 | 35.15 | 22.11 | 1429.69 | 1 | 0.196753 |
| 318 | 25.33 | 2.89 | 1.56 | 450 | 50.3 | 27.06 | 22.64 | 1188.44 | 1 | 0.274105 |
| 319 | 13.46 | 2.13 | 9.52 | 609 | 37.24 | 20.15 | **42.62** | 1373.65 | 1 | 0.142229 |
| 320 | 28.87 | 6.34 | 6.16 | 568 | 65.25 | 18.93 | 15.81 | 1645.14 | 1 | 0.492232 |
| 321 | 1.25 | 0.94 | **1.56** | 641 | 42.01 | 25.18 | 32.81 | 1685.77 | 0.97 | 0.0839308 |
| 322 | 0.81 | **2.59** | 0.49 | 618 | 39.54 | 28.01 | 32.45 | 1660.04 | 1 | 0.123746 |
| 323 | 1.13 | **2.74** | 0.32 | 620 | 37.87 | **38.5** | 23.62 | 1599.64 | 0.83 | 0.0630259 |
| 324 | 7.56 | 1.18 | 0.84 | 595 | 35.65 | 34.02 | 30.33 | 1555.04 | 0.86 | 0.0589137 |
| 325 | 3.63 | 0.79 | 0.95 | 633 | 41.64 | 24.97 | 33.39 | 1543.96 | 1 | 0.15853 |
| 326 | 4.4 | **7.69** | 1.88 | 637 | 33.08 | **33.51** | **33.41** | 1429.31 | 0.44 | 0.0204965 |
| 327 | 13.36 | 6.19 | 6.68 | 614 | 41.75 | 34.24 | 24.02 | 1351.38 | 1 | 0.178994 |
| 328 | 23.51 | 0 | 0.4 | 251 | 68.65 | 13.61 | 17.74 | 914.28 | 1 | 0.593404 |
| 329 | 11.16 | 0 | 0.8 | 251 | 42.45 | 25.55 | 32 | 462.63 | 1 | 0.325294 |
| 330 | 17.56 | 0.98 | 2.28 | 615 | 58.37 | 20.11 | 21.52 | 1326.7 | 1 | 0.285637 |
| 331 | 1.09 | 0.16 | 0.62 | 642 | 34.17 | 27.6 | **38.23** | 1081.44 | 0.55 | 0.0497915 |
| 332 | 1.76 | 0.16 | 0 | 626 | 47.42 | 25.87 | 26.71 | 1121.59 | 1 | 0.17873 |
| 333 | 1.46 | **2.1** | **1.46** | 618 | 37.8 | 31.59 | 30.61 | 1133.18 | 0.79 | 0.0522177 |
| 334 | 8.58 | 0.83 | 1.49 | 606 | 41.71 | 25.71 | 32.58 | 1172.96 | 1 | 0.146508 |
| 335 | 3.46 | 1.48 | 0.99 | 607 | 39.01 | 29.31 | 31.68 | 1185.84 | 0.51 | 0.0460312 |
| 336 | 6.84 | 4.34 | 6.01 | 599 | 33.59 | 33.15 | 33.26 | 1022.56 | 1 | 0.129375 |
| 337 | 0.32 | 0.16 | 0 | 631 | 38.95 | 28.65 | 32.39 | 973.14 | 0.72 | 0.050131 |
| 338 | 0.48 | **0.48** | **0.48** | 629 | 34.35 | **36.27** | 29.39 | 1026.2 | 0.8 | 0.0583927 |
| 339 | 4.58 | 0.32 | 1.9 | 633 | 39.99 | 31.72 | 28.29 | 1055.27 | 0.98 | 0.0927266 |
| 340 | 1.59 | 0.64 | 0 | 627 | 42.97 | 32.63 | 24.4 | 1112.19 | 0.99 | 0.102916 |
| 341 | 2.28 | **5.71** | **3.43** | 613 | 32.6 | **38.21** | 29.19 | 965.5 | 0.65 | 0.034272 |
| 342 | 9.05 | 5.36 | 6.2 | 597 | 37.51 | 37.41 | 25.08 | 1048.74 | 0.6 | 0.0318051 |
| 343 | 1.94 | 0 | 0 | 669 | 57.35 | 20.11 | 22.54 | 2016.94 | 1 | 0.506971 |
| 344 | 0.76 | 0.15 | 0.3 | 661 | 38.07 | 34.33 | 27.6 | 1781.88 | 0.85 | 0.0563376 |
| 345 | 1.37 | 0.61 | 0.15 | 657 | 45.59 | 26.63 | 27.78 | 1612.67 | 1 | 0.126055 |
| 346 | 27.7 | 0.16 | 0.47 | 639 | 63.84 | 17.07 | 19.09 | 1897.82 | 1 | 0.571376 |
| 347 | 4.95 | 0.15 | 0.46 | 647 | 49.71 | 27.07 | 23.21 | 1671.43 | 1 | 0.247173 |
| 348 | 20.7 | 0.63 | 1.26 | 633 | 52.83 | 25.19 | 21.97 | 1737.42 | 1 | 0.401842 |
| 349 | 5.78 | 4.38 | 1.88 | 640 | 36.5 | **38.76** | 24.74 | 1592.14 | 0.99 | 0.109039 |
| 350 | 9.92 | 8.78 | 8.62 | 615 | 29.08 | **29.92** | **41** | 1662.35 | 0.36 | 0.00242572 |
| 351 | 8.54 | 0.16 | 0 | 644 | 59.49 | 21.24 | 19.27 | 2069.97 | 1 | 0.510383 |
| 352 | 4.35 | 0.81 | 0.97 | 620 | 39.25 | 29.62 | 31.13 | 1919.03 | 1 | 0.13351 |
| 353 | 11.31 | 2.01 | 2.55 | 548 | 55.76 | 21.63 | 22.61 | 1569.59 | 1 | 0.167141 |
| 354 | 8.56 | 6.04 | 4.36 | 596 | 39.35 | 35.11 | 25.55 | 1731.94 | 0.99 | 0.106049 |
| 355 | 2.36 | 0.47 | 2.36 | 635 | 44.99 | 28.52 | 26.5 | 1734.16 | 1 | 0.12822 |
| 356 | 28.59 | 0.32 | 0.32 | 633 | 87.73 | 6.4 | 5.87 | 3150.12 | 1 | 0.797943 |
| 357 | 0.87 | **3.05** | **1.96** | 459 | 31.58 | 31.9 | 36.52 | 1126.02 | 0.55 | 0.0296089 |
| 358 | 1.99 | **4.42** | 1.55 | 452 | 31.04 | 32.63 | 36.32 | 1193.8 | 0.76 | 0.0507469 |
| 359 | 6.38 | 4.31 | 4.66 | 580 | 37.49 | 31.96 | 30.55 | 1497.98 | 0.89 | 0.0636962 |
| 360 | 1.07 | 0 | 0 | 655 | 47.6 | 27.21 | 25.19 | 1707.46 | 1 | 0.254142 |
| 361 | 2.37 | 0 | 0.32 | 634 | 40.47 | 27.02 | 32.5 | 1381.98 | 0.98 | 0.0874477 |
| 362 | 9.79 | 2.64 | 3.77 | 531 | 45.21 | 26.3 | 28.49 | 1223.21 | 1 | 0.182212 |
| 363 | 0.15 | **0.46** | 0 | 652 | 33.43 | 30.99 | **35.58** | 1371.31 | 0.7 | 0.0399833 |
| 364 | 0.8 | 0 | 0 | 627 | 42.12 | 27.74 | 30.15 | 1434.21 | 1 | 0.152709 |
| 365 | 1.15 | **1.15** | **1.15** | 607 | 38.74 | 27.49 | 33.78 | 1410.3 | 0.82 | 0.057901 |
| 366 | 2.05 | 1.37 | 0.46 | 438 | 46.08 | 18.61 | 35.31 | 1045.51 | 1 | 0.166484 |
| 367 | 2.08 | **6.47** | **2.77** | 433 | 34.25 | 28.91 | **36.85** | 1010.52 | 0.67 | 0.0396326 |
| 368 | 2.95 | 1.69 | 1.9 | 474 | 38.36 | 23.88 | 37.75 | 1184.92 | 0.92 | 0.0897483 |
| 369 | 2.56 | **3.42** | 1.71 | 468 | 38.18 | 29.43 | 32.4 | 1155.98 | 0.93 | 0.0803685 |
| 370 | 9.11 | 5.06 | 3.04 | 593 | 46.88 | 26.63 | 26.49 | 1421.42 | 1 | 0.186945 |
| 371 | 11.18 | 9.27 | 10.7 | 626 | 33.8 | 30.83 | **35.37** | 1253.29 | 0.76 | 0.0463641 |
| 372 | 0.17 | **0.34** | **0.17** | 593 | 34.68 | 32.68 | 32.65 | 1414.2 | 0.89 | 0.0662082 |
| 373 | 0.34 | **1.71** | 0.17 | 586 | 40.46 | 30.25 | 29.29 | 1734.98 | 0.94 | 0.0794697 |
| 374 | 0.16 | **0.49** | **0.32** | 617 | 37.24 | 29.75 | 33.01 | 1697.22 | 1 | 0.13728 |
| 375 | 27.21 | 0 | 0 | 588 | 65.64 | 16.63 | 17.72 | 1681.64 | 1 | 0.604237 |
| 376 | 0 | **0.32** | **0.16** | 620 | 35.83 | 33.15 | 31.02 | 1375.3 | 0.86 | 0.0570667 |
| 377 | 0.48 | 0.16 | 0 | 628 | 35.26 | 33.7 | 31.04 | 1355.3 | 0.56 | 0.0336972 |
| 378 | 2.37 | 0 | 0 | 591 | 55.13 | 19.32 | 25.55 | 1498.33 | 1 | 0.220905 |
| 379 | 17.33 | 1.08 | 0.72 | 554 | 51.39 | 23.11 | 25.5 | 1581.56 | 1 | 0.198506 |
| 380 | 4.98 | 2.41 | 1.72 | 582 | 37.03 | 25.02 | **37.95** | 1467.59 | 1 | 0.117582 |
| 381 | 11.15 | 7.2 | 6.17 | 583 | 38.99 | 29.41 | 31.59 | 1373.33 | 1 | 0.120031 |
| 382 | 0.51 | 0 | 0 | 590 | 38.61 | 32.05 | 29.33 | 1368.9 | 0.97 | 0.0893493 |
| 383 | 0.51 | **1.36** | 0 | 588 | 42.08 | 24.8 | 33.12 | 1378.59 | 1 | 0.141435 |
| 384 | 0 | **0** | **0.16** | 632 | 33.96 | 33.16 | 32.87 | 1429.35 | 0.3 | 0.00213732 |
| 385 | 0 | **0.16** | **0** | 626 | 32.96 | **34.56** | 32.48 | 1427.13 | 0.79 | 0.0474248 |
| 386 | 1.3 | **2.76** | 0.97 | 617 | 38.24 | 25.89 | 35.87 | 1416.16 | 0.62 | 0.0312812 |
| 387 | 5.79 | 3.48 | 3.48 | 604 | 33.19 | **36.68** | 30.13 | 1441.7 | 0.74 | 0.0467207 |
| 388 | 0.16 | 0 | 0 | 624 | 34.97 | 32.8 | 32.23 | 1441.19 | 0.57 | 0.0243587 |
| 389 | 0.32 | **1.46** | **0.32** | 618 | 34.09 | 32.69 | 33.21 | 1514.91 | 0.58 | 0.0261524 |
| 390 | 5.68 | 0.17 | 1 | 599 | 33.84 | 31.78 | **34.38** | 1381.56 | 0.76 | 0.0488014 |
| 391 | 0.96 | **1.28** | 0.64 | 624 | 32.2 | 31.48 | **36.32** | 1408 | 0.71 | 0.0405001 |
| 392 | 19.44 | 2.64 | 1.98 | 607 | 59.6 | 19.91 | 20.49 | 1479.38 | 1 | 0.389622 |
| 393 | 23.9 | 0 | 0.33 | 615 | 65.71 | 18.58 | 15.71 | 1571.82 | 1 | 0.524664 |

Supplementary table 1. Branch IDs and concordance factors for MQS-all-loci species tree (figure 1).

ID : BranchID (see figure S1); gCF: Gene concordance factor (%); gDF1: Gene discordance factor (%) for NNI-1 branch; gDF2: Gene discordance factor (%) for NNI-2 branch; gN: Number of trees decisive for the branch; sCF: Site concordance factor (%) averaged over 100 quartets; sDF1: Site discordance factor (%) for alternative quartet 1; sDF2: Site discordance factor (%) for alternative quartet 2; sN: Number of informative sites averaged over 100 quartets; PP: ASTRAL posterior probability support (-t3 option); Length: Branch length. Gene and site discordance factors in bold are >= GCF or SCF for that branch.
