## Supplementary Table S4 for "A phylogenomic analysis of *Nepenthes* (Nepenthaceae)"

| **ID** | **gCF** | **gDF1** | **gDF2** | **gN** | **sCF** | **sDF1** | **sDF2** | **sN** | **Label** | **Length** |
| --- | --- | --- | --- | --- | --- | --- | --- | --- | --- | --- |
| 198 | 0.75 | 0 | 0 | 266 | 36.36 | 30.97 | 32.68 | 297.74 |  | 0.000918848 |
| 199 | 0 | 0 | 0 | 266 | 38.2 | 32.43 | 29.37 | 304.48 | 0.998 | 0.000447726 |
| 200 | 0.38 | 0 | 0 | 266 | 41.34 | 24.38 | 34.27 | 293.85 | 0.961 | 0.000514909 |
| 201 | 0 | 0 | 0 | 266 | 37.26 | 35.17 | 27.57 | 273.59 | 0.962 | 0.000476102 |
| 202 | 18.49 | 0 | 0 | 265 | 61 | 20.71 | 18.29 | 298.36 | 1 | 0.00164318 |
| 203 | 0 | 0.38 | 0 | 266 | 38.93 | 28.46 | 32.62 | 249.56 | 0.986 | 0.000281257 |
| 204 | 16.23 | 0 | 0 | 265 | 59.45 | 20.9 | 19.65 | 274.45 | 1 | 0.00100734 |
| 205 | 1.5 | 0.38 | 0 | 266 | 53.17 | 19.45 | 27.38 | 300.37 | 1 | 0.00123651 |
| 206 | 13.36 | 0.76 | 0.38 | 262 | 40.41 | 33.95 | 25.65 | 304.33 | 1 | 0.00116189 |
| 207 | 3.4 | 1.13 | 0.38 | 265 | 41.35 | 24.65 | 34 | 286.46 | 0.58 | 0.000616866 |
| 208 | 6.84 | 7.98 | 4.94 | 263 | 41.21 | 33.09 | 25.69 | 265.18 | 0.951 | 0.000660227 |
| 209 | 0 | 0 | 0 | 265 | 40.3 | 31.08 | 28.62 | 310.22 | 1 | 0.000819512 |
| 210 | 0 | 0 | 0 | 265 | 44.45 | 29.72 | 25.83 | 305.79 | 1 | 0.000573787 |
| 211 | 0 | 0 | 0 | 266 | 36.36 | 34.38 | 29.26 | 281.08 | 0.902 | 0.000251478 |
| 212 | 1.13 | 0 | 0 | 266 | 43.64 | 28.08 | 28.28 | 317.78 | 1 | 0.000714915 |
| 213 | 1.13 | 0 | 0 | 266 | 52.74 | 25.14 | 22.12 | 398.26 | 1 | 0.00193454 |
| 214 | 43.23 | 0 | 0 | 266 | 77.26 | 11.06 | 11.68 | 577.05 | 1 | 0.00416599 |
| 215 | 0.38 | 0.38 | 0.38 | 266 | 37.7 | 23.63 | 38.67 | 373.78 | 0.999 | 0.000600446 |
| 216 | 0 | 0 | 0 | 266 | 33.43 | 30.01 | 36.55 | 349.36 | 0.359 | 0.000307461 |
| 217 | 0 | 0 | 0 | 266 | 39.12 | 30.65 | 30.23 | 354.72 | 0.229 | 0.000322392 |
| 218 | 0.75 | 0 | 0 | 265 | 35.12 | 26.39 | 38.49 | 367.02 | 0.998 | 0.00101545 |
| 219 | 29.81 | 0 | 0 | 265 | 82.8 | 10.14 | 7.06 | 604.83 | 1 | 0.00370016 |
| 220 | 23.31 | 2.26 | 3.38 | 266 | 59.28 | 18.93 | 21.79 | 271.62 | 1 | 0.000915167 |
| 221 | 37.97 | 10.53 | 6.39 | 266 | 67.13 | 18.84 | 14.03 | 247.31 | 1 | 0.00139704 |
| 222 | 8.65 | 4.14 | 5.64 | 266 | 40.81 | 25.79 | 33.41 | 239.59 | 1 | 0.000466535 |
| 223 | 43.02 | 2.26 | 1.89 | 265 | 63.92 | 17.13 | 18.95 | 310.19 | 1 | 0.00168298 |
| 224 | 12.45 | 2.26 | 1.89 | 265 | 41.07 | 28.21 | 30.71 | 243.85 | 1 | 0.000430483 |
| 225 | 13.36 | 7.25 | 9.92 | 262 | 50.04 | 26.62 | 23.34 | 254.33 | 1 | 0.00124164 |
| 226 | 19.16 | 6.13 | 3.07 | 261 | 54.24 | 30.34 | 15.42 | 262.61 | 1 | 0.000972086 |
| 227 | 0 | 0 | 0 | 265 | 31.04 | 36.1 | 32.86 | 350.04 | 0.374 | 0.000160588 |
| 228 | 0.75 | 0.75 | 0 | 265 | 38.93 | 31.15 | 29.92 | 315.4 | 0.992 | 0.00042176 |
| 229 | 18.42 | 0 | 0 | 266 | 74.88 | 10.67 | 14.46 | 464.05 | 1 | 0.00345083 |
| 230 | 22.93 | 4.89 | 3.38 | 266 | 48.95 | 22.74 | 28.31 | 323.92 | 1 | 0.000881746 |
| 231 | 8.27 | 10.9 | 3.76 | 266 | 34.53 | 41.22 | 24.25 | 313.6 | 0.621 | 0.000496973 |
| 232 | 18.42 | 0.38 | 1.13 | 266 | 67.71 | 16.49 | 15.8 | 327.64 | 1 | 0.00184159 |
| 233 | 7.89 | 4.14 | 3.38 | 266 | 27.23 | 30.3 | 42.47 | 188.18 | 0.973 | 0.000328394 |
| 234 | 5.66 | 2.64 | 2.64 | 265 | 49.06 | 15.85 | 35.09 | 164.36 | 1 | 0.000597063 |
| 235 | 2.63 | 3.95 | 4.61 | 152 | 26.95 | 23.14 | 49.91 | 69.73 | 0.995 | 0.000510846 |
| 236 | 13.82 | 6.58 | 1.97 | 152 | 39.51 | 30.57 | 29.92 | 71.93 | 0.565 | 0.000180348 |
| 237 | 18.42 | 4.14 | 2.26 | 266 | 64.33 | 17.31 | 18.35 | 197.33 | 1 | 0.000873635 |
| 238 | 2.63 | 0 | 0 | 266 | 53.78 | 25.64 | 20.58 | 414.72 | 1 | 0.00166419 |
| 239 | 28.57 | 0 | 0 | 266 | 69.43 | 16.74 | 13.82 | 618.79 | 1 | 0.00355631 |
| 240 | 23.68 | 9.4 | 4.51 | 266 | 51.55 | 27.44 | 21.01 | 580.13 | 1 | 0.00169725 |
| 241 | 38.78 | 2.66 | 6.84 | 263 | 54.86 | 20.43 | 24.71 | 639.64 | 1 | 0.0023493 |
| 242 | 79.39 | 3.05 | 3.44 | 262 | 82.63 | 10.75 | 6.62 | 986.1 | 1 | 0.00675913 |
| 243 | 27.38 | 12.17 | 7.22 | 263 | 39.64 | 35.8 | 24.56 | 541.2 | 0.992 | 0.000984036 |
| 244 | 0.38 | 0.38 | 0 | 266 | 40.04 | 27.5 | 32.46 | 377.86 | 1 | 0.00070308 |
| 245 | 7.17 | 0 | 1.51 | 265 | 51.65 | 10.33 | 38.03 | 459.42 | 1 | 0.001855 |
| 246 | 24.32 | 12.74 | 2.7 | 259 | 47.02 | 43.38 | 9.6 | 529.24 | 1 | 0.00147717 |
| 247 | 0 | 0 | 0 | 266 | 43.03 | 26.81 | 30.17 | 333.64 | 0.656 | 0.000445449 |
| 248 | 38.26 | 0 | 0 | 264 | 72.26 | 9.99 | 17.75 | 565.92 | 1 | 0.00429893 |
| 249 | 38.98 | 11.42 | 17.32 | 254 | 55.53 | 20.82 | 23.66 | 356.3 | 1 | 0.00253181 |
| 250 | 0 | 0.38 | 0.38 | 266 | 40.37 | 30.84 | 28.79 | 260.19 | 0.995 | 0.000571608 |
| 251 | 0.75 | 0 | 0 | 266 | 38.85 | 33.42 | 27.73 | 250.86 | 0.871 | 0.000360175 |
| 252 | 0 | 0.38 | 0.38 | 266 | 46.8 | 28.47 | 24.73 | 337.86 | 1 | 0.00102556 |
| 253 | 3.83 | 0 | 0.77 | 261 | 43.81 | 21.24 | 34.95 | 341.97 | 1 | 0.00119304 |
| 254 | 6.15 | 1.54 | 1.15 | 260 | 51.55 | 21.53 | 26.92 | 349.53 | 1 | 0.00121195 |
| 255 | 4.15 | 6.79 | 2.64 | 265 | 36.91 | 34.67 | 28.43 | 313.53 | 0.201 | 0.000534418 |
| 256 | 17.74 | 4.91 | 3.02 | 265 | 50.37 | 30.14 | 19.49 | 298.76 | 0.998 | 0.000902816 |
| 257 | 10.53 | 4.74 | 2.11 | 190 | 46.2 | 34.18 | 19.62 | 269.01 | 1 | 0.00233268 |
| 258 | 2.26 | 0.38 | 0 | 266 | 54.05 | 23.94 | 22 | 314.79 | 1 | 0.00172729 |
| 259 | 10.53 | 0.38 | 0.38 | 266 | 57.97 | 18.41 | 23.62 | 251.25 | 1 | 0.00169446 |
| 260 | 8.27 | 3.76 | 15.04 | 133 | 36.51 | 28.32 | 35.17 | 124.76 | 0.946 | 0.00141731 |
| 261 | 0.75 | 0 | 0 | 266 | 40.77 | 26.15 | 33.07 | 311.21 | 1 | 0.000783696 |
| 262 | 1.13 | 0 | 0.38 | 266 | 36.61 | 32.43 | 30.96 | 347.53 | 0.986 | 0.000746303 |
| 263 | 1.88 | 1.5 | 1.13 | 266 | 50.63 | 31.12 | 18.25 | 334.9 | 1 | 0.00109492 |
| 264 | 9.96 | 0 | 0 | 261 | 60.9 | 22.57 | 16.53 | 265.21 | 1 | 0.00170628 |
| 265 | 1.13 | 0 | 0 | 265 | 39.93 | 23.06 | 37 | 281.15 | 1 | 0.00112893 |
| 266 | 0 | 1.14 | 0 | 263 | 41.05 | 32.56 | 26.39 | 229.82 | 0.96 | 0.000897746 |
| 267 | 1.14 | 0 | 0 | 264 | 41.67 | 26.9 | 31.44 | 219.55 | 0.566 | 0.000463913 |
| 268 | 1.88 | 3.01 | 2.63 | 266 | 36.9 | 30.21 | 32.88 | 258.41 | 0.842 | 0.000436809 |
| 269 | 8.3 | 4.53 | 1.13 | 265 | 50.68 | 28.57 | 20.75 | 223.64 | 1 | 0.00094035 |
| 270 | 0 | 0 | 0 | 260 | 45.32 | 22.8 | 31.88 | 248.26 | 1 | 0.0017075 |
| 271 | 1.29 | 0 | 0 | 232 | 37.58 | 21.59 | 40.83 | 238.89 | 1 | 0.00230386 |
| 272 | 2 | 4.5 | 0.5 | 200 | 36.41 | 34.77 | 28.82 | 251.38 | 0.998 | 0.00107669 |
| 273 | 4.12 | 3.61 | 3.61 | 194 | 40.24 | 35.82 | 23.93 | 261.63 | 1 | 0.00142425 |
| 274 | 1.3 | 0.87 | 0 | 231 | 39.9 | 25.1 | 34.99 | 218.7 | 1 | 0.00110712 |
| 275 | 4.43 | 2.53 | 1.27 | 158 | 38.95 | 37.69 | 23.36 | 140.05 | 0.966 | 0.00114706 |
| 276 | 0 | 0 | 0 | 266 | 31.5 | 27.02 | 41.48 | 371.04 | 0.524 | 0.000426326 |
| 277 | 5.32 | 0 | 0 | 263 | 39.33 | 26.37 | 34.3 | 384.15 | 0.999 | 0.000907678 |
| 278 | 0 | 0.38 | 0 | 262 | 36.69 | 33.98 | 29.34 | 371.27 | 0.499 | 0.000705602 |
| 279 | 0 | 3.83 | 0 | 261 | 41.76 | 32.36 | 25.89 | 372.75 | 1 | 0.00144974 |
| 280 | 0.38 | 0 | 4.58 | 262 | 32.82 | 33.26 | 33.92 | 339.75 | 0.905 | 0.000961768 |
| 281 | 0.82 | 1.23 | 5.76 | 243 | 38.32 | 21.79 | 39.89 | 321.89 | 0.157 | 0.00105895 |
| 282 | 5.3 | 0 | 0 | 264 | 49.23 | 21.94 | 28.83 | 366.37 | 1 | 0.00139937 |
| 283 | 10.98 | 1.14 | 2.65 | 264 | 57.55 | 21.74 | 20.71 | 444.32 | 1 | 0.00199765 |
| 284 | 30.68 | 3.79 | 4.17 | 264 | 66.62 | 17.16 | 16.22 | 451.18 | 1 | 0.00263525 |
| 285 | 11.32 | 6.42 | 7.92 | 265 | 39.45 | 28.67 | 31.87 | 392.49 | 0.877 | 0.00091707 |
| 286 | 0.75 | 0 | 0 | 265 | 43.04 | 26.44 | 30.52 | 277.12 | 1 | 0.000984741 |
| 287 | 1.89 | 3.4 | 1.89 | 265 | 40.28 | 29.17 | 30.55 | 221.22 | 1 | 0.00112589 |
| 288 | 49.24 | 0.38 | 0 | 262 | 77.79 | 12.8 | 9.41 | 364.56 | 1 | 0.0036211 |
| 289 | 0.45 | 0.9 | 4.93 | 223 | 51.64 | 14.83 | 33.52 | 73.86 | 0.994 | 0.00103585 |
| 290 | 17.05 | 1.55 | 1.55 | 129 | 75.64 | 11.74 | 12.62 | 68.63 | 1 | 0.00899072 |
| 291 | 10.76 | 1.79 | 1.35 | 223 | 41.92 | 33.25 | 24.82 | 72.39 | 0.997 | 0.000970982 |
| 292 | 41.47 | 2.3 | 1.38 | 217 | 80.88 | 8.88 | 10.24 | 269.85 | 1 | 0.0048268 |
| 293 | 27.65 | 2.65 | 1.89 | 264 | 69.55 | 17.15 | 13.3 | 220.14 | 1 | 0.00209528 |
| 294 | 16.67 | 10.83 | 27.5 | 120 | 32.88 | 33.7 | 33.43 | 80.27 | 0.838 | 0.000708815 |
| 295 | 0 | 0 | 0 | 266 | 39.51 | 29.78 | 30.71 | 346.54 | 0.967 | 0.000450852 |
| 296 | 1.51 | 2.64 | 0.38 | 265 | 35.09 | 32.17 | 32.75 | 352.93 | 0.989 | 0.000453097 |
| 297 | 13.58 | 0.75 | 0.75 | 265 | 60.59 | 18.42 | 20.99 | 413.61 | 1 | 0.00185402 |
| 298 | 0.38 | 0 | 0 | 266 | 46.08 | 26.8 | 27.12 | 346.59 | 1 | 0.000665731 |
| 299 | 1.88 | 0 | 0.38 | 266 | 37.24 | 31.11 | 31.65 | 342.02 | 0.988 | 0.000620314 |
| 300 | 20.68 | 0 | 0 | 266 | 79.42 | 7.99 | 12.59 | 463.45 | 1 | 0.0030032 |
| 301 | 0 | 0 | 0.75 | 266 | 29.71 | 42.1 | 28.18 | 186.75 | 0.826 | 7.822E-05 |
| 302 | 1.88 | 0 | 0 | 266 | 49.48 | 23.45 | 27.07 | 194.09 | 0.921 | 0.000169993 |
| 303 | 3.38 | 4.89 | 4.89 | 266 | 38.96 | 32.67 | 28.37 | 176.15 | 0.608 | 0.000517889 |
| 304 | 0 | 0 | 0.38 | 266 | 32.12 | 31.57 | 36.31 | 192.96 | 1 | 0.000596396 |
| 305 | 1.13 | 0 | 0 | 266 | 40.92 | 29.31 | 29.77 | 186.72 | 0.999 | 0.000474542 |
| 306 | 0.38 | 0 | 0 | 266 | 40.51 | 32.82 | 26.67 | 217.02 | 1 | 0.000487808 |
| 307 | 1.88 | 6.02 | 1.13 | 266 | 33.35 | 36.88 | 29.76 | 243.73 | 1 | 0.000796642 |
| 308 | 7.14 | 4.14 | 4.14 | 266 | 34.83 | 27.75 | 37.42 | 228.64 | 0.992 | 0.000605239 |
| 309 | 0 | 1.13 | 0 | 266 | 33.3 | 37.19 | 29.52 | 165.76 | 0.121 | 0.000314175 |
| 310 | 2.52 | 0 | 0 | 119 | 46.5 | 15.77 | 37.73 | 94 | 0.93 | 0.00134061 |
| 311 | 1.88 | 0 | 3.01 | 266 | 49.64 | 25.86 | 24.51 | 119.2 | 0.999 | 0.000469254 |
| 312 | 1.13 | 0 | 0 | 266 | 42.73 | 30.52 | 26.75 | 182.52 | 0.953 | 0.000222678 |
| 313 | 13.53 | 3.01 | 2.26 | 266 | 47.15 | 29.07 | 23.78 | 202.02 | 1 | 0.000534631 |
| 314 | 4.14 | 0.75 | 1.13 | 266 | 43.06 | 25.64 | 31.31 | 168.82 | 0.912 | 0.000190663 |
| 315 | 0.38 | 0 | 0 | 266 | 32.23 | 33.83 | 33.93 | 343.66 | 0.963 | 0.000466152 |
| 316 | 0.75 | 0 | 0 | 266 | 37.14 | 31.18 | 31.69 | 349.55 | 1 | 0.000575026 |
| 317 | 8.3 | 0.38 | 0 | 265 | 58.35 | 18.85 | 22.79 | 359.37 | 1 | 0.00132825 |
| 318 | 18.87 | 4.53 | 3.02 | 265 | 58.95 | 14.41 | 26.64 | 322.83 | 1 | 0.0012415 |
| 319 | 13.53 | 9.4 | 19.17 | 266 | 43.35 | 20.91 | 35.73 | 285.84 | 0.761 | 0.000724467 |
| 320 | 3.01 | 0 | 0 | 266 | 58.52 | 23.12 | 18.36 | 382.47 | 1 | 0.00158049 |
| 321 | 0 | 0 | 0 | 266 | 34.03 | 34.56 | 31.41 | 294.39 | 0.693 | 0.000239789 |
| 322 | 6.79 | 0 | 0.38 | 265 | 41.81 | 28.89 | 29.3 | 279.6 | 1 | 0.00071828 |
| 323 | 0.38 | 0.38 | 0.76 | 264 | 38.44 | 36.68 | 24.88 | 280.16 | 1 | 0.000705352 |
| 324 | 0 | 0 | 0 | 266 | 34.08 | 33.25 | 32.67 | 321.26 | 0.999 | 0.000462185 |
| 325 | 0 | 0 | 0 | 266 | 39.14 | 27.25 | 33.61 | 317.63 | 0.998 | 0.000361218 |
| 326 | 0 | 0 | 0 | 266 | 28.92 | 44.02 | 27.06 | 315.39 | 1 | 0.00086251 |
| 327 | 0 | 0 | 0 | 266 | 36.11 | 27.7 | 36.18 | 357.82 | 0.994 | 0.000321472 |
| 328 | 1.88 | 0.38 | 0.75 | 266 | 41.49 | 29.5 | 29.02 | 315.86 | 1 | 0.000598862 |
| 329 | 5.26 | 3.01 | 3.01 | 266 | 36.43 | 23.79 | 39.78 | 316.84 | 1 | 0.000851036 |
| 330 | 0 | 0 | 0 | 265 | 32.44 | 35.47 | 32.09 | 335.39 | 0.769 | 0.00033207 |
| 331 | 0.38 | 0 | 2.64 | 265 | 32.12 | 34.1 | 33.79 | 328.67 | 1 | 0.000653537 |
| 332 | 1.88 | 1.13 | 1.88 | 266 | 42.53 | 33.38 | 24.09 | 338.59 | 1 | 0.000741301 |
| 333 | 6.02 | 0 | 0 | 266 | 61.39 | 21.04 | 17.57 | 307.79 | 1 | 0.00122568 |
| 334 | 1.13 | 0 | 0 | 265 | 47.67 | 27.39 | 24.94 | 177.83 | 1 | 0.000420583 |
| 335 | 1.9 | 0 | 0 | 263 | 34.23 | 27.12 | 38.65 | 169.93 | 1 | 0.000814595 |
| 336 | 7.73 | 4.29 | 1.29 | 233 | 35.81 | 18.61 | 45.58 | 151.07 | 1 | 0.00102357 |
| 337 | 0.38 | 0.75 | 0 | 265 | 35.8 | 38.3 | 25.9 | 158.44 | 0.37 | 0.000263136 |
| 338 | 6.04 | 0 | 0 | 265 | 44.72 | 30.43 | 24.84 | 146.51 | 0.512 | 0.00029058 |
| 339 | 1.13 | 1.89 | 1.13 | 265 | 34.7 | 32.98 | 32.32 | 150.51 | 0.998 | 0.000419343 |
| 340 | 7.52 | 0 | 0 | 266 | 57.21 | 25.76 | 17.03 | 176.56 | 1 | 0.000968876 |
| 341 | 3.85 | 5 | 1.92 | 260 | 40.46 | 47.64 | 11.9 | 106.89 | 1 | 0.00103065 |
| 342 | 7.34 | 2.82 | 9.6 | 177 | 41.49 | 14.79 | 43.71 | 77.69 | 0.981 | 0.00162061 |
| 343 | 1.5 | 0 | 0 | 266 | 37.89 | 26.33 | 35.79 | 371.28 | 1 | 0.00109537 |
| 344 | 2.63 | 0 | 0 | 266 | 52.2 | 23.02 | 24.78 | 350.16 | 1 | 0.00131485 |
| 345 | 11.28 | 0 | 0 | 266 | 66.15 | 15.92 | 17.93 | 349.11 | 1 | 0.00145957 |
| 346 | 12.03 | 0.75 | 2.63 | 266 | 40.95 | 25.51 | 33.54 | 240.59 | 1 | 0.000562457 |
| 347 | 26.82 | 8.81 | 4.98 | 261 | 51.08 | 28.68 | 20.24 | 308.77 | 1 | 0.00131203 |
| 348 | 4.89 | 6.02 | 3.01 | 266 | 36.37 | 37.04 | 26.59 | 223.65 | 0.995 | 0.000327168 |
| 349 | 18.87 | 3.3 | 2.83 | 212 | 55.31 | 23.37 | 21.33 | 236.75 | 1 | 0.00137518 |
| 350 | 2.26 | 1.5 | 1.13 | 266 | 40.44 | 27.55 | 32.01 | 304.69 | 0.994 | 0.000610428 |
| 351 | 0.75 | 0 | 0.75 | 266 | 41.82 | 33.99 | 24.19 | 310.95 | 0.977 | 0.000364322 |
| 352 | 0.38 | 0.38 | 0 | 266 | 37.31 | 37.97 | 24.72 | 294.63 | 0.538 | 0.000392899 |
| 353 | 2.26 | 3.02 | 0 | 265 | 43.17 | 34.63 | 22.2 | 317.28 | 1 | 0.00124773 |
| 354 | 1.88 | 0.75 | 0.75 | 266 | 51.53 | 19.96 | 28.51 | 271.37 | 0.777 | 0.000288618 |
| 355 | 7.89 | 4.14 | 6.77 | 266 | 35.27 | 35.99 | 28.74 | 223.75 | 0.408 | 0.000563687 |
| 356 | 3.01 | 0 | 0.75 | 266 | 36.77 | 26.52 | 36.7 | 292.99 | 1 | 0.000676802 |
| 357 | 0.38 | 0 | 0 | 266 | 39.51 | 31.55 | 28.94 | 320.32 | 0.992 | 0.000581815 |
| 358 | 6.39 | 0 | 0.38 | 266 | 60.72 | 21.57 | 17.71 | 396.08 | 1 | 0.00183252 |
| 359 | 2.63 | 0 | 0.38 | 266 | 43.95 | 26.04 | 30.01 | 300.77 | 0.999 | 0.00057656 |
| 360 | 13.58 | 1.13 | 0.38 | 265 | 67.66 | 15.31 | 17.02 | 291.94 | 1 | 0.00137827 |
| 361 | 0.75 | 1.5 | 0.38 | 266 | 33.91 | 33.7 | 32.39 | 281.05 | 1 | 0.000515886 |
| 362 | 4.89 | 1.13 | 1.5 | 266 | 46.56 | 30.22 | 23.22 | 344.98 | 1 | 0.00136189 |
| 363 | 5.28 | 2.64 | 0.75 | 265 | 30.18 | 34 | 35.82 | 295.58 | 0.886 | 0.00053725 |
| 364 | 1.5 | 0 | 0 | 266 | 42.28 | 33.02 | 24.7 | 331.15 | 1 | 0.000882619 |
| 365 | 7.89 | 1.88 | 5.64 | 266 | 41.49 | 24.03 | 34.48 | 312.67 | 1 | 0.000882731 |
| 366 | 11.9 | 1.59 | 2.38 | 252 | 37.35 | 31.65 | 31 | 304.99 | 1 | 0.000922295 |
| 367 | 0.75 | 0 | 0 | 266 | 39.7 | 29.55 | 30.75 | 308.15 | 0.886 | 0.000468046 |
| 368 | 20.83 | 0.38 | 0.38 | 264 | 66.19 | 14.78 | 19.03 | 345.59 | 1 | 0.00200447 |
| 369 | 5.26 | 0 | 0.38 | 266 | 58.23 | 20.4 | 21.37 | 318.09 | 1 | 0.00105195 |
| 370 | 8.27 | 3.01 | 1.13 | 266 | 38.07 | 31.42 | 30.51 | 243.26 | 0.514 | 0.000437083 |
| 371 | 3.01 | 3.38 | 3.01 | 266 | 38.26 | 41.11 | 20.64 | 250.08 | 0.859 | 0.000458259 |
| 372 | 16.67 | 3.03 | 3.41 | 264 | 43.82 | 30.01 | 26.16 | 256.06 | 1 | 0.000870846 |
| 373 | 1.5 | 0 | 0 | 266 | 44.23 | 30.12 | 25.65 | 251.69 | 1 | 0.000511228 |
| 374 | 6.83 | 1.2 | 2.41 | 249 | 47.02 | 19.13 | 33.85 | 210.79 | 1 | 0.000584011 |
| 375 | 0 | 0 | 0 | 265 | 33.29 | 31.06 | 35.64 | 281.46 | 0.674 | 0.000286073 |
| 376 | 0.38 | 0.38 | 0.38 | 264 | 45.82 | 30.93 | 23.25 | 261.66 | 1 | 0.0005077 |
| 377 | 0.75 | 0.38 | 0.38 | 265 | 40.09 | 30.99 | 28.92 | 292.36 | 1 | 0.000908762 |
| 378 | 1.88 | 2.26 | 0 | 266 | 35.74 | 41.17 | 23.09 | 276.84 | 0.037 | 0.0006566 |
| 379 | 1.5 | 0 | 0.38 | 266 | 41.13 | 31.58 | 27.28 | 287.43 | 0.997 | 0.000562698 |
| 380 | 0.75 | 1.89 | 3.4 | 265 | 31.33 | 39.83 | 28.84 | 283.39 | 0.998 | 0.000553925 |
| 381 | 6.79 | 3.4 | 0.75 | 265 | 51.16 | 24.94 | 23.9 | 264.48 | 0.999 | 0.000508591 |
| 382 | 8.27 | 9.4 | 7.14 | 266 | 32.09 | 30.4 | 37.51 | 309.49 | 0.101 | 0.000597943 |
| 383 | 5.71 | 1.9 | 0.48 | 210 | 57.02 | 26.54 | 16.44 | 238.88 | 1 | 0.00248002 |
| 384 | 0 | 0 | 0 | 266 | 34.7 | 25.48 | 39.81 | 296.05 | 0.991 | 0.000432295 |
| 385 | 14.72 | 0 | 0 | 265 | 63.93 | 14.47 | 21.6 | 311.52 | 1 | 0.00125417 |
| 386 | 0 | 0.38 | 0 | 266 | 36.15 | 33.14 | 30.71 | 273.41 | 1 | 0.000772025 |
| 387 | 3.02 | 0 | 0 | 265 | 40.59 | 31.55 | 27.85 | 237.45 | 0.98 | 0.000476591 |
| 388 | 0 | 0 | 0 | 266 | 31.46 | 33.74 | 34.8 | 254.98 | 0.983 | 0.000415161 |
| 389 | 3.8 | 0 | 0.76 | 263 | 42.42 | 30.92 | 26.66 | 237.57 | 0.999 | 0.00079111 |
| 390 | 4.15 | 1.13 | 0 | 265 | 41.02 | 34.76 | 24.22 | 240.69 | 0.777 | 0.00064998 |
| 391 | 1.14 | 0 | 0 | 264 | 37.32 | 35.26 | 27.41 | 257.12 | 0.753 | 0.00035239 |

Supplementary table 2. Branch IDs and concordance factors for 177-supermatrix tree (figure 1) based on 177 gene alignments used to build the supermatrix. ID : BranchID (see figure S1); gCF: Gene concordance factor (%); gDF1: Gene discordance factor (%) for NNI-1 branch; gDF2: Gene discordance factor (%) for NNI-2 branch; gN: Number of trees decisive for the branch; sCF: Site concordance factor (%) averaged over 100 quartets; sDF1: Site discordance factor (%) for alternative quartet 1; sDF2: Site discordance factor (%) for alternative quartet 2; sN: Number of informative sites averaged over 100 quartets; PP: ASTRAL posterior probability support (-t3 option); Length: Branch length. Gene and site discordance factors in bold are >= GCF or SCF for that branch.
